# Mechanistic model-driven exometabolomic characterisation of human dopaminergic neuronal metabolism

**DOI:** 10.1101/2021.06.30.450562

**Authors:** German Preciat, Agnieszka B. Wegrzyn, Edinson Lucumi Moreno, Cornelius C.W. Willacey, Jennifer Modamio, Fatima L. Monteiro, Diana El Assal, Alissa Schurink, Miguel A.P. Oliveira, Zhi Zhang, Ben Cousins, Hulda S. Haraldsdóttir, Siham Hachi, Susanne Zach, German Leparc, Yin Tat Lee, Bastian Hengerer, Santosh Vempala, Michael A. Saunders, Amy Harms, Enrico Glaab, Jens C. Schwamborn, Ines Thiele, Thomas Hankemeier, Ronan M.T. Fleming

## Abstract

Starting with a comprehensive generic reconstruction of human metabolism, we generated high-quality, constraint-based, genome-scale, cell-type and condition specific models of metabolism in human dopaminergic neurons, the cell type most vulnerable to degeneration in Parkinson*’*s disease. They are a synthesis of extensive manual curation of the biochemical literature on neuronal metabolism, together with novel, quantitative, transcriptomic and targeted exometabolomic data from human stem cell-derived, midbrain-specific, dopaminergic neurons *in vitro*. Thermodynamic constraint-based modelling enabled qualitatively accurate and moderately quantitatively accurate prediction of dopaminergic neuronal metabolite exchange fluxes, including predicting the consequences of metabolic perturbations in a manner also consistent with literature on monogenic mitochondrial Parkinson*’*s disease. These dopaminergic neurons models provide a foundation for a quantitative systems biochemistry approach to metabolic dysfunction in Parkinson*’*s disease. Moreover, the plethora of novel mathematical and computational approaches required to develop them are generalisable to study any other disease associated with metabolic dysfunction.

## Introduction

Parkinson*’*s disease (PD) is a complex multifactorial disease with an incidence range between 5 and 35 per 100,000 population [1]. In sporadic PD, it is well established that degeneration followed by cell loss is selective for certain brainstem nuclei [2, 3, 4, 5]. For example, the motor symptoms of PD are mainly caused by the dysfunction, degeneration and death of substantia nigra dopaminergic neurons [6]. A combination of intrinsic anatomical, morphological, physiological and biochemical characteristics has been proposed to be shared by neurons selectively vulnerable to degeneration in PD [7]. They tend to have long unmyelinated axons, large axonal trees, a high number of synapses, autonomous pace-making activity, broad action potentials [8] and calcium-mediated feed-forward stimulation of mitochondrial oxidative phosphorylation [9]. These features are thought to place high material and energetic demands on such neurons for axonal trafficking, cycling of synaptic vesicles, re-establishment of ionic gradients following firing and maintenance of cellular structural integrity, including proteome homeostasis [10]. Therefore, it is of great interest to characterise the balance of metabolic supply and demand in substantia nigra dopaminergic neurons, both in health and in disease.

One of the significant challenges to research neuronal disease mechanisms is the requirement for accessible and faithful cellular models that recapitulate the main cellular features of healthy and diseased cells. Since the publication of the seminal work describing the generation of human-induced pluripotent stem cells (iPSCs) from human dermal fibroblasts [11], reprogramming of differentiated cells into iPSCs has become an essential tool to study neurodegenerative diseases, such as PD [12, 13, 14, 15, 16, 17]. From iPSCs, it is possible to derive robust, stable human neuroepithelial stem cells, which can be differentiated clonally and efficiently into neural tube lineages, midbrain dopaminergic neurons [18], offering an accessible approach to study the normal and abnormal metabolism of midbrain dopaminergic neurons *in vitro*. Therefore, genome-scale characterisation of the metabolic status of such midbrain dopaminergic neurons is of major interest but has not yet been reported.

Constraint-based reconstruction and analysis (COBRA) is a genome-scale computational modelling approach [19] that provides a molecular mechanistic framework for experimental design, integrative analysis of prior biochemical knowledge with experimental data and quantitative prediction of physicochemical and biochemical feasible phenotypic states [20]. In particular, quantitative bioanalytical chemistry [21, 22, 23] has been combined with constraint-based modelling of metabolism [24] to enable context-specific biochemical interpretation of metabolomic data, e.g., to discover differences in glycolytic versus oxidative metabolism in different lymphoblastic leukaemia cell lines [25], and to characterise metabolic changes influencing pluripotency and cell fate in stem cells [26].

Constraint-based modelling of neuronal metabolism is challenging, but progress has been made. Cakir et al. [27] developed a stoichiometric model of central metabolic interactions between astrocytes and neurons, encompassing 217 reactions among 216 metabolites. It was further updated to include 630 metabolic reactions and 570 genes detected in the transcriptomic data of six neurodegenerative diseases [28]. It was able to predict major metabolic fluxes that were in agreement with literature data and was further used to predict potential biomarkers for several neurodegenerative diseases; however, they have yet to be experimentally validated. Most recently, Abdik et al. has developed a mouse brain-specific genome-scale metabolic model that allows the study of different mouse models of PD in terms of their metabolism [29]. Lewis et al. [30] developed a stoichiometric model of metabolism and mitochondrial function in astrocytes and either glutamatergic, GABAergic or cholinergic neurons. The emphasis was on cerebral energy metabolism, including central metabolism, mitochondrial metabolic pathways, and pathways relevant to anabolism and catabolism of the neurotransmitters glutamate, GABA, and acetylcholine.

In parallel, dynamic kinetic models have been developed that focus on particular aspects of dopaminergic neuronal function. Extending a model [31], Cloutier et al. [32] developed a phenomenological kinetic model of central metabolic brain energy metabolism including capillary, neuronal, astrocyte and subcellular mitochondrial compartments. Qi et al. [33] developed a model of the nigrostriatal dopamine pathway, representing processes as products of power-law functions [34], and used it to predict key determinants of dopamine metabolism associated with the dysregulation of dopamine homeostasis.

We present two genome-scale constraint-based models of midbrain-specific dopaminergic neuronal metabolism giving preference to either *in vitro* experimental data (condition specific; iDopaNeuroC) or dopaminergic neuronal literature curation (cell-type specific; iDopaNeuroCT). Manual literature curation was used to establish the activity or inactivity of a core set of metabolic genes and reactions, as well as presence or absence of metabolites characteristic of dopaminergic neuronal metabolism. In parallel, exometabolomic and trancriptomic data was generated from midbrain-specific dopaminergic neuronal cultures, obtained by differentiation of normal human neuroepithelial stem cells. Exometabolomic data from four platforms, including liquid chromatography-mass spectrometry and gas chromatography-mass spectrometry, were used to quantify energy-related metabolites including biogenic amines and organic acids, as well as neurochemical metabolites in fresh and spent culture media.

The combined results of literature curation and omics data generation were integrated with a stoichiometrically consistent and flux consistent derivative of a Recon3D [35], a comprehensive reconstruction of human metabolism, using a novel model generation pipeline [36]. An ensemble of thermodynamically flux consistent context-specific models were generated and the models with the highest predictive accuracy, evaluated against exometabolomic data from dopaminergic neuronal cultures, were designated iDopaNeuroC and iDopaNeuroCT.A breakthrough in predictive fidelity demonstrated by the iDopaNeuro models was demonstrated by comparison with independent exometabolomic data on pharmacologically perturbed dopaminergic neuronal cultures. Furthermore, we developed and applied a novel approach to predict the most informative metabolites to measure, by future exometabolomic experiments, in order to maximally reduce uncertainty in the feasible steady-state flux solution space. These models are a quantitative, interdisciplinary characterisation of dopaminergic neuronal metabolism at genome-scale. They provide a validated platform for experimental data-driven mechanistic computational modelling and optimal design of experiments, and ultimately provide an objective, quantitative framework for development of drugs targeted toward the aetiopathogeneses of Parkinson*’*s Disease.

## Materials and Methods

A summary of the materials and methods is provided below and all details are provided as supplementary information. An overview of the generation, validation and application of the dopaminergic neuronal metabolic models, iDopaNeuroC and iDopaNeuroCT, is shown on Fig. 1.

**Figure 1:**
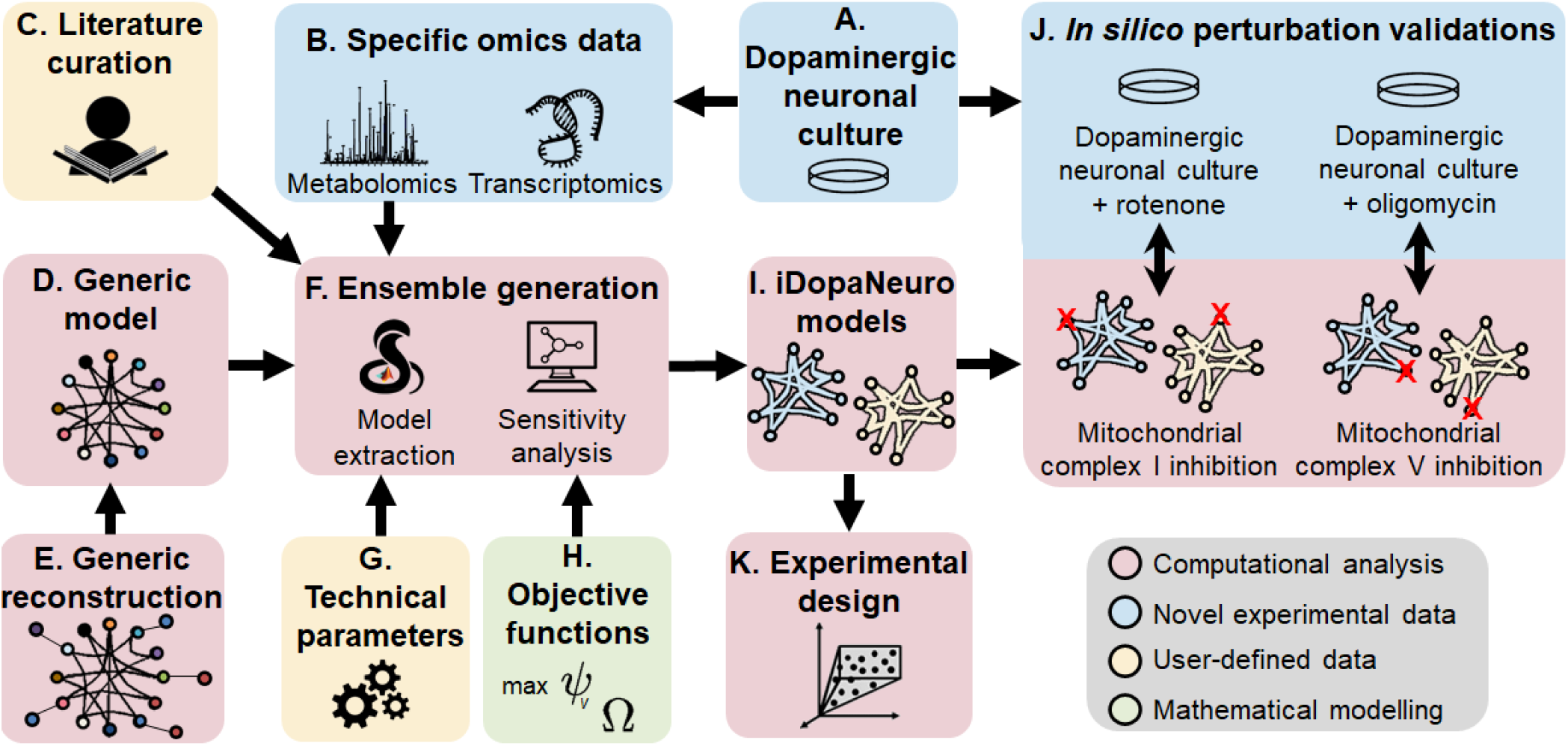
Overview of generation, validation and application of the iDopaNeuro models. Human neuroepithelial stem cells were differentiated into midbrain-specific dopaminergic neuronal cultures *in vitro* (**A**). Transcriptomic and targeted exometabolomic data were generated from fresh and spent media samples (**B**). This condition-specific omics data together with cell-type-specific data derived from manual curation of the literature on dopaminergic neuronal metabolism (**C**) were integrated with a generic metabolic model (**D**) derived from a comprehensive reconstruction of human metabolism (**E**). An ensemble of candidate dopaminergic neuronal metabolic models were generated (**F**), as function of technical parameters (**G**) and mathematical modelling approaches (**H**). The models with the highest predictive fidelity were identified (**I**), which gave preference to either *in vitro* experimental data (iDopaNeuro condition specific; iDopaNeuroC) or dopaminergic neuronal literature curation (iDopaNeuro cell-type specific; iDopaNeuroCT). The predictive fidelity of the iDopaNeuro models was validated by comparison of predictions with independent exometabolomic data generated on perturbations to normal dopaminergic neuronal metabolism *in vitro* (**J**). Finally, the iDopaNeuro models were used prospectively to design exometabolomic experiments to generate new constraints on the variables currently most uncertain in the model (**K**).

### 1 Experiments

Fig.2 provides an overview of the experimental protocol, illustrating the timing of events.

**Figure 2:**
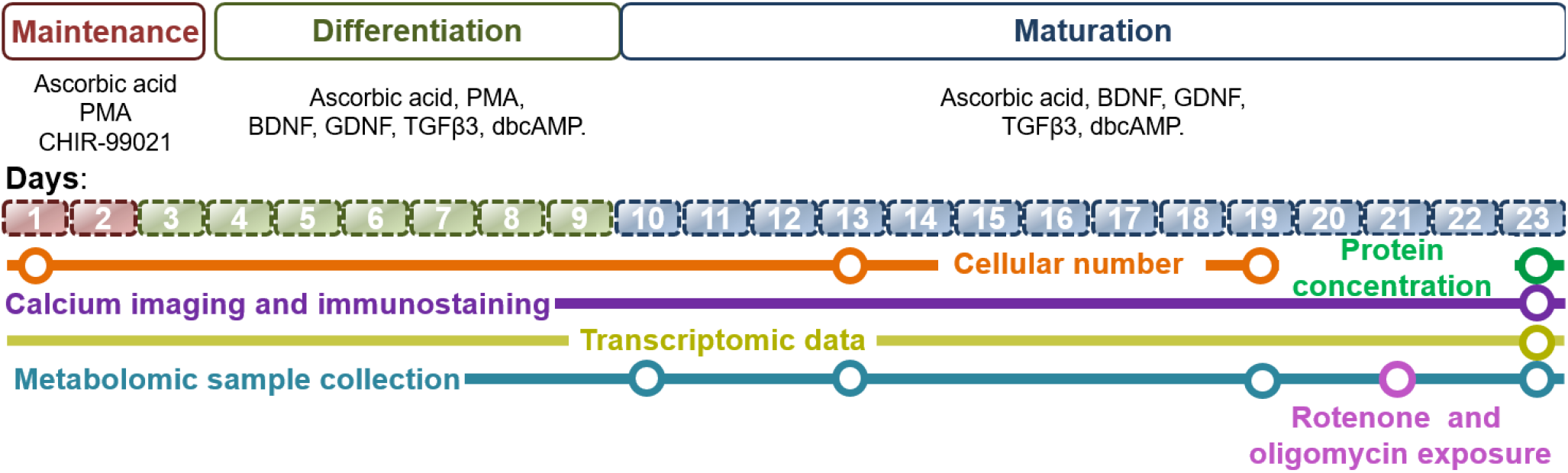
Experimental protocol overview. Human neuroepithelial stem cells (hNESC) were differentiated into midbrain dopaminergic neurons. The cell number in each culture well was counted on day 1, 13, 19 and estimated for day 23. Spent media samples for metabolomic analyses were collected at days 10, 13, 19 and 23. Samples were analysed with both GC-MS or LC-MS. At day 23, live cells were subjected to calcium imaging followed by immunostaining assays, and collection of parallel samples for transcriptomic analysis. The media composition at the various stages of cell culture were as follows. Maintenance stage (red): maintenance medium containing ascorbic acid, purmorphamine (PMA) and the aminopyrimidine CHIR-99021 (CHIR). Differentiation stage (green): differentiation medium containing ascorbic acid, Brain-derived neurotrophic factor (BDNF), glial cell-derived neurotrophic factor (GDNF), Transforming Growth Factor Beta 3 (TGF*β*3), dbcAMP and PMA. Maturation stage (blue): differentiation media without PMA.

#### 1.1 *In vitro* cell culture

##### Summary

A human neuroepithelial stem cell line from a healthy human donor was maintained and differentiated toward midbrain-specific dopaminergic neurons using an established protocol [18], summarised in Section 1.1.1. Cellular morphology was monitored during differentiation and after sufficient time had elapsed (23 days), calcium imaging and automated image analysis were used to assess electrophysiological activity, using an established pipeline [37], summarised in Section 1.1.2. Immunofluorescent staining was used to identify differentiated cell types (Section 1.1.3). Normal cultures were also perturbed metabolically, either pharmacologically, or by changing the fresh medium carbon source (Section 1.1.4).

###### 1.1.1 Dopaminergic neuronal maintenance and differentiation

Generation of an *in vitro* culture of midbrain-specific dopaminergic neurons followed an established protocol [18], with the following adaptions. The human neuroepithelial stem cells were cultivated in mTESR1 medium (StemCell technologies, #05850) on 6-well dishes coated with Matrigel (Corning, #354263). The composition of each fresh medium, to the extent that it has been defined by the manufacturer, is detailed in Table S-1.

###### N2B27 medium preparation

The culture medium, denoted *N2B27 medium*, was used as the basis to prepare both maintenance and differentiation media. 50.25 mL of culture medium was obtained by mixing 24 mL Neurobasal medium (Invitrogen/Life Technologies), 24 mL of DMEM/F12 medium (Invitrogen/Life Technologies) supplemented with 0.5 mL of 1% penicillin and streptomycin (Life Technologies), 0.5 mL of 200 mM L-glutamine (Life Technologies), 0.5 mL of B27 supplement without Vitamin A (Life Technologies) and 0.25 mL of N2 supplement (Life Technologies). The final concentration of the media composition is fully detailed in Table S-1.

###### Plate coating

Nunc cell-culture treated 6-well plates (ThermoFisher scientific, Roskilde, Denmark) were coated with 1% Matrigel (Discovery Labware, Inc., Two Oak Park, Bedford, MA, USA, Catalogue number 354277, lot number 3318549) in 600*μ*L of knockout DMEM (1X) medium.

###### Cell seeding and maintenance

At the time of cell seeding, the knockout DMEM (1X) medium from the coating step was removed from each well and the K7 hNESC line was seeded in three replicate wells. The medium to maintain the hNESC in culture, denoted *maintenance medium* (red in Fig. 2), is based on N2B27 medium with 0.5 *μ*M PMA (Enzo life sciences), 3 *μ*M CHIR (Axon Medchem) and 150 *μ*M ascorbic acid (Sigma Aldrich). The cell seeding was done by preparing 5 × 10^6^ cells/mL in 50% matrigel in maintenance medium and adding 200 *μ*L of this preparation to obtain approximately 0.2 mm thick layer of cells in three dimensions within Matrigel, with 4 × 10^5^cells per well. After the Matrigel and cell mixture was added to the well, the plate was incubated for 2 min at 37 °C to gelate the matrigel layer, the plate was then taken out of the incubator and 2.8 mL of maintenance medium was added and the plate was incubated at 37 °C and 5% CO_2_ for 48 h.

###### Neuronal differentiation and maturation

The *differentiation medium with PMA* (green in Fig. 2) preparation to induce the differentiation of hNESC towards midbrain dopaminergic neurons consisted of N2B27 medium with 200 *μ*M ascorbic acid, 0.01 ng/*μ*L BDNF (Peprotech), 0.01 ng/*μ*L GDNF (Peprotech), 0.001 ng/*μ*L TGF*β*3 (Peprotech), 2.5 *μ*M dbcAMP (Sigma Aldrich) and 1 *μ*M PMA. This medium preparation was completely replaced every 2 days during the next 6 days of culture in the differentiation process. For the maturation of differentiated neurons, PMA is required to be absent from the differentiation medium. This *differentiation medium without PMA* (Blue in Fig. 2) was used from day 9 onwards and 50% media replacement every 2 days for 3 weeks. Protein concentration (mg/mL) was measured using a bicinchoninic acid protein assay [38].

###### 1.1.2 Microscopy and calcium imaging

To monitor cellular morphology during differentiation, bright field images were acquired every 48 h for 23 days of differentiation using a Zeiss Axiovert 40 CFL microscope equipped with a cooled charge-coupled device based camera (Zeiss AxioCam MRm, Zeiss). At day 23 in culture, calcium imaging was done with a Fluo-4 AM green-fluorescent calcium indicator dye. After the differentiation medium was removed, 1 mL of 5*μ*M cell permeant Fluo-4 AM (Invitrogen/Life Technologies, F-14201) in neurobasal medium was added to selected wells of a 6-well plate at room temperature. Full frame fluorescence images of size 2560×2160 pixels were acquired using an epifluorescence microscope (Leica DMI6000 B, Germany) equipped with a cooled sCMOS camera (Neo 5.5, Andor technology, UK) and both were controlled with Micro-manager (version 1.4) [39]. Images were sampled at a rate of approximately 10 Hz for about 2 min, stored as image stacks and analysed off-line using an established pipeline for automated calcium image analyses [37]. For each segmented neuron, we measured fluorescence traces as relative changes in fluorescence intensity over time.

###### 1.1.3 Immunofluorescence staining assay

Immunostaining for a dopaminergic marker, tyrosine hydroxylase (TH) and a pan neuronal marker, Class III *β*-tubulin (TUBbIII) were used to identify differentiated dopaminergic neurons. Immunostaining for tyrosine hydroxylase (TH) positive differentiated neurons was performed on wells of a 6-well plate after day 25 of differentiation. Differentiated cells were fixed with 4 % PFA in 1× phosphate-buffered saline (PBS) (15 min), followed by permeabilisation with 0.05% Triton-X 100 in 1× PBS (3 min on ice), and blocking with 10% fetal calf serum (FCS) in 1× PBS (1 h). After washing with 1× PBS, the primary antibodies mouse anti-TUB*β*III (1:1000, Covance, Germany), rabbit anti-TH (1:1000, Santa Cruz biotechnology, Germany) and chicken anti-GFAP (1:1000, Merck Millipore, Germany), were incubated for 90 min at 25 °C. After washing with 1× PBS, the secondary antibodies Alexa Fluor 488 Goat Anti-Rabbit (1:1000, Invitrogen), Alexa Fluor 568 Goat Anti-Mouse (1:1000, Invitrogen), Alexa Fluor 647 Goat Anti-chicken (1:1000, Invitrogen) and Hoechst 33342 to stain DNA (1:10000, Invitrogen), were incubated overnight at 4 °C. After washing with 1× PBS, confocal images of areas of selected wells were acquired, using a confocal microscope (Zeiss LSM 710).

###### 1.1.4 Metabolic perturbations

At day 23 of differentiation, midbrain-specific dopaminergic neurons were exposed for 24hrs to two different mitochondrial inhibitors, 12.5 *μ*M Rotenone (Merck) and 12.5 *μ*M Oligomycin (Abcam). Triplicate experiments resulted in nine samples per perturbation. Maintenance media and maintenance media containing 1% DMSO (place) were used as controls. After 24 hours the extracellular spent media was collected and snap-frozen using liquid nitrogen. Furthermore, a galactose perturbation was performed by replacing glucose with galactose concentration at 3 g/L at day 23 as the primary sugar source. Exometabolomic analysis of fresh and spent culture media was used to measure exchange of metabolites between the media and perturbed dopaminergic neuronal cell culture, as described previously.

#### 1.2 Transcriptomics

##### Summary

A human neuroepithelial stem cell line from a healthy human donor was maintained and differentiated toward midbrain-specific dopaminergic neurons using the same protocol [18] as described in Section 1.1. After sufficient differentiation (Fig. 2), cell culture sample RNA was extracted and sequenced (Section 1.2.1) from *in vitro* cell culture samples, RNA-sequencing was employed and raw data was analysed (Section 1.2.2), such that the output was quantitative expression for each gene, in units of Fragments Per Kilobase of transcript per Million mapped reads (FPKM).

###### 1.2.1 RNA extraction and sequencing

A kit (Ambion Magmax™-96 total RNA isolation kit, Life Sciences) was used for RNA extraction. Magnetic beads were used to isolate nucleic acids. Afterwards, the samples were washed and purified with DNAase. The RNA obtained was eluted in 50*μM* elution buffer. Fragment Analyzer (Aligent Technologies Inc.) was used to measure RNA quality and concentration.

The sequencing library preparation was done using 200 ng of total RNA input with the TrueSeq RNA Sample Prep Kit v3-Set B (RS-122-2002, Illumina Inc, San Diego, CA) producing a 275 bp fragment including adapters in average size. In the final step before sequencing, twelve individual libraries were normalised and pooled together using the adapter indices supplied by the manufacturer. Pooled libraries were then clustered on the cBot Instrument (Illumina Inc, San Diego, CA) using the TruSeq SR Cluster Kit v3-cBot-HS (GD-401-3001, Illumina, Inc, San Diego, CA) sequencing was then performed as 78 bp, single reads and 7 bases index read on an Illumina HiSeq300 instrument using the TruSeq SBS Kit HS-v3 (50-cycle) (FC-401-3002, Illumina Inc, San Diego, CA).

###### 1.2.2 RNA sequence analysis

The raw RNA-seq data were analysed with a custom-made RNA-seq analysis pipeline, which included publicly available software (SAMtools, version 0.1.18; FASTX-Toolkit, version 0.0.14) [40] and custom-made python scripts. The current version of the pipeline is available at https://git-r3lab.uni.lu/zhi.zhang/rnaseqhs. The RNA-seq analysis pipeline consists of six main steps: (i) quality control for the raw RNA-seq reads; (ii) preprocessing of the raw RNA-seq reads to remove adapters and low-quality sequences; (iii) alignment of the reads to the human reference genome; (iv) assembly of the alignments into transcripts and (v) quantification of the expression levels of each gene. Briefly, the raw RNA-seq reads (length 52 nucleotides, single-end) of each sample were checked using FastQC (version 0.11.2) to determine the read quality. Adapter sequences and low-quality sequences were removed using cutadapt (version 1.10) [41] with default settings. Reads with less than 25 nucleotides were excluded from further analysis. Next, the alignment of RNA-seq reads against the human reference genome (NCBI build37.2, downloaded from iGenome of Illumina) was performed using TopHat2 (version 2.0.13) [42]. Alignment results were processed using Cufflinks (version 2.2.1) [43] for assembly of transcripts with default parameter settings. The quantification of transcript expression was estimated by normalised FPKM (Fragments Per Kilobase of transcript per Million mapped reads) and counts at gene level by cuffnorm (version 2.2.1) [43]. In order to obtain one expression value per gene, we used the transcript with the highest average expression as representative for the corresponding gene, since measurements for low-abundance transcripts are less reliable.

#### 1.3 Analytical chemistry

##### Summary

Targeted, quantitative, exometabolomic data was generated from fresh and spent culture media using four partially overlapping metabolomic platforms described below as AccQ-Tag method, benzoyl chloride (BzCl), dimethylaminophenacyl bromide (DmPaBr), and GC-MS [44, 45, 46, 47] (Section 1). Table S-2 contains a list of target metabolites analysed with three LC-MS and one GC-MS platforms covering biogenic amines, amino acids, organic acids and glucose with a total of 77 metabolites targeted. Where a metabolite was quantified by more than one platform, its concentration was assumed to be that measured by the platform with the lowest relative standard deviation in the repeated measurements of a quality control sample [48]. Subsequently, differences in metabolite concentrations over time were either used to generate a context-specific model (Section 3.1.5) or kept independent from the model generation pipeline and used to test *in silico* model predictions (Section 3.2.2).

###### 1.3.1 LC-MS profiling of biogenic amines and amino acids using the AccQ-Tag method

The analysis of 75 biogenic amines (Table S-2) was performed with an established LC-MS method [44]. Briefly, 15 *μ*L of culture medium was extracted by adding 400 *μ*L of ice-cold methanol, 55 *μ*L of ice-cold milliQ water, 10*μ*L of tris(2-carboxyethyl)phosphine (TCEP; 1*μ*g/*μ*L) and 10 *μ*L of a mixture of stable isotope labelled internal standards (IS; Table S-2). After centrifugation, all supernatants were transferred into 1.5 mL tubes and the liquid extracts were evaporated in a vacuum concentrator (Labconco, Kansas City, MO, USA) to dryness. The dried extracts were dissolved in 80 *μ*\*μ*L borate buffer (pH 9) and mixed with 20 *μ*L of pure acetonitrile containing 3 *μ*g/*μ*L AccQ-Tag derivatisation reagent (Waters, Etten-Leur, Netherlands) for the chemical derivatisation of the primary and/or secondary amine groups. The derivatisation reaction was performed at 55 °C for 30 min after which the samples were centrifuged at 16000×g and 4 °C for 2 min and 80 *μ*L of the supernatant was transferred into LC vials for sample injection. 1 *μ*L of the liquid extract was injected onto the analytical column for the analysis. Measurements were performed with a Waters Acquity ultra-high pressure liquid chromatography (UPLC) (Milford, MA, USA) hyphenated with Agilent 6460 triple quadrupole mass spectrometer (Palo Alto, CA, USA). Chromatographic separation was achieved on a Water Acquity HSS T3 C18 UPLC column (2.1×100 mm, 1.7*μ*m) and the metabolites were identified based on their retention time and multiple reaction monitoring (MRM) transitions from their protonated precursor ions of the AccQ-Tag derivates into common product ion of 171 m/z.

###### 1.3.2 Analysis of amines and neurochemical metabolites using LC-MS/MS derivatized by benzoyl chloride (BzCl)

The samples were analyzed following the protocol by [45] for the quantitative targeted analysis of amines and neurochemical metabolites. The dried samples were reconstituted in 10 *μ*L H2O whilst maintained on ice. To start the derivatization reaction, 10 *μ*L of 100 mM sodium carbonate (pH 9.4) was added, followed by 10 *μ*L of 2% benzoyl chloride in ACN (v/v), and vortexed immediately for 10 seconds triggering the spontaneous Sn1 reaction at room temperature. The reaction was quenched by the addition of 20 *μ*L H2O with 1 % sulphuric acid after 5 minutes. Isotopically labelled internal standards were synthesized by derivatizing a mixture of all targeted metabolites using the 13C labelled BzCl reagent. The internal standard mix was added to the quenched derivatized samples and 50 *μ*L H2O was added to reduce the organic content. Samples were analyzed by LC-MS/MS using a Waters Acquity UPLC Class II (Milford, USA) coupled to an ABSciex QTrap 6500 series (Framingham, USA). The analytical column used was a Waters BEH-C18 column (1mm×100 mm, 1.8 *μ*m, 180 Å) with an injection of 5 *μ*L, maintained at 60°C. A gradient from 0.1% v/v formic acid and 10 mM ammonium formate in water to 100% acetonitrile over 20 minutes with a flow rate of 100 *μ*L/min was used for the separation of metabolites. Samples were automatically integrated using the vendor software AbSciex MultiQuant Workstation Quantitative Analysis for QTrap 6500 series.

###### 1.3.3 Analysis of energy-related metabolites using LC-MS/MS derivatized by dimethylaminophenacyl bromide (DmPABr)

The samples were analyzed following the protocol by [47] for the quantitative targeted analysis of energy-related metabolites The dried content was reconstituted in 10 *μ*L of DMSO/DMF to dissolve the remaining content. Then, 10 *μ*L of triethanolamine (750 mM) was added to the vial, followed by 10 *μ*L of DmPABr (82 mM). The sealed Eppendorf vial was placed into a shaking incubator for 60 minutes at 65°C to complete the derivatization. A total of 10 *μ*L of formic acid (30 mg/mL) was added to the vial to quench the reaction with an additional 30 minutes in the shaking incubator. Then, 5 *μ*L of DmPABr-D_6_-labelled metabolites (DmPABr-2D) or DmPA-^13^C_2_ (DmPABr-13C)[49] were added. Before vortexing, 45 *μ*L of ACN was added and transferred to an HPLC vial for analysis. Samples were analyzed by LC-MS/MS using a Waters Acquity UPLC Class II (Milford, USA) coupled to an ABSciex QTrap 6500 series (Framingham, USA). The analytical column used was a Waters AccQ-tag column (2.1mm×100 mm, 1.8 *μ*m, 180 Å) with an injection of 1 *μ*L, maintained at 60°C. A gradient from 0.1% v/v formic acid and 10 mM ammonium formate in water to 100% acetonitrile over 15 minutes with a flow rate of 700 *μ*L/min was used for the separation of metabolites. Samples were automatically integrated using the vendor software AbSciex MultiQuant Workstation Quantitative Analysis for QTrap 6500 series.

###### 1.3.4 GC-MS profiling of polar metabolites

Twenty-four polar metabolites (Table S-2) were analysed in culture media using a modified version of an in-house GC-MS platform [46]. Because of the high abundance of D-glucose and L-lactic acid in culture media, samples were diluted 1:299 (v/v) in milliQ water. Fifty microliters of both diluted and non-diluted culture medium were extracted with 425 *μ*L of an extraction solvent (methanol/water, 94%/6%; volume/volume) containing stable isotope labelled internal standards (Table S-2). Four hundred microliters of the supernatant were transferred into a 1.5 mL tube and the solvent was evaporated in a vacuum concentrator (Labconco, Kansas City, MO, USA). Dry samples were resuspended for the oximation reaction in 35 *μ*L of pyridine containing methoxyamine hydrochloride (15 *μ*g/*μ*L) and kept at 30 °C for 90 min. After the oximation of the aldehyde groups on reducing sugars and organic acids, samples were further derivatised with silylation reaction for 60 min in an orbital shaker (VWR, Germany). This reaction was carried out by adding 40 *μ*L of MSTFA (N-methyl-N-trimethylsilylacetamide) into the samples. Subsequently, samples were centrifuged at 16000×g and room temperature for 5 min and 70 *μ*L of the supernatant was transferred into silanized glass inserts. The GC-MS measurements were performed on an Agilent 7890A GC System coupled to a single quadruple 5975C Mass Selective Detector. One microliter of the sample was injected with splitless injection. The analytes were separated on an Agilent HP-5MS Ultra Inert capillary GC column (30 m, 250 *μ*m ID, 0.25 *μ*m film thickness). Metabolite identification was carried out by using the retention time of the chemical standards and mass spectral similarity of the fragmentation pattern with NIST MS Search Software (v2.0). The metabolite quantification was performed based on the specific fragment ion for each polar metabolite (Table S-2). Both peak extraction and integration were performed by using the vendor*’*s software (Agilent MassHunter Quantitative software v5.0).

###### 1.3.5 Concentration determination

To determine the concentration, calibration samples were measured covering a wide concentration range. The peak areas for all analytes measured in the samples were converted to peak area ratios using

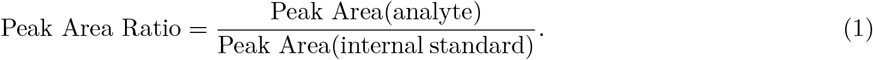

The analytes were assigned an exact isotope labelled internal standard, where possible, and if the internal standard was not available then the closest eluting peak was assigned (internal standard selection table detailed below). The internal standard was used to correct for differences in extraction efficiency, matrix effect, ion suppression, and instrument response (Table S-2). For absolute quantification, a linear regression was calculated for the calibration lines by fitting a linear model *y* = *ax* + *b* in RStudio. If the intercept was not significant for the fit, then a linear model was changed to *y* = *ax* + 0 (Table S-2).

###### 1.3.6 Quality control assessment

To ensure that the data meets the quality standards throughout the experimental procedure, quality control (QC -pooled samples) samples were taken through the experiment process. The quality control samples were injected every 10 samples throughout the batch. The relative standard deviation of the quality control samples *RSD* (*qc*) was assessed for quality per metabolite by using

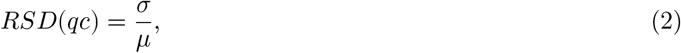

where *σ* is the standard deviation of the measured quality control samples, and *μ* is the mean of the control samples measurements. *RSD*(*qc*) was then used as a measurement uncertainty for all the measured samples. Furthermore, for analytes covered by more than one analytical method, measurement with the lowest *RSD*(*qc*) value was chosen for further analysis.

## 2 Reconstruction

### Summary

An established generic human metabolic reconstruction, Recon3D [35], was used as a foundation to generate the iDopaNeuro models. Literature review was performed to define active and inactive reactions and genes (Tables S-3, S-4, S-5, S-6 and S-7), transport reactions (Table S-3), degradation pathways and quantitative constraints necessary to represent the requirement for molecular turnover in a non-growing, non-dividing dopaminergic neuron (Tables S-8 and S-9). When specific information on human substantia nigra pars compacta dopaminergic neurons was not present in the literature, information from other neuronal types, cerebral tissue, or rodent data was used (Tables S-4 and S-6). Additionally, neuron-specific biomass maintenance requirements (section 2.2) were estimated for each biomass constituent and the first reaction in the corresponding degradation pathway, or pathways, for each biomass constituent was identified. This enabled the generation of turnover constraints to ensure that the material requirements for maintenance of a dopaminergic neuron were met. The bounds on the rate of each exchange reaction, corresponding to a constituent of the defined fresh cell culture medium, plus reversible extracellular transport reactions, for water, carbon dioxide and oxygen, were qualitatively set to eliminate uptake of all other metabolites (Table S-1).

#### 2.1 Generic human metabolic reconstruction

Recon2 [50] and its successor Recon3D [35] are increasingly comprehensive, genome-scale reconstructions of human metabolism. They also provide information about gene-protein-reaction associations that associate each metabolic gene with the corresponding enzyme or enzyme complex and reaction in a Boolean manner. They are generic reconstructions formed by amalgamation and manual curation of metabolic reactions across human metabolism occurring in many cell types. The metabolic identity of a cell is strongly influenced by its ability to transport metabolites across its extracellular membrane. Therefore, particular emphasis was placed on curation of dopaminergic neuronal transport reactions. Furthermore, a key characteristic of dopaminergic neurons is their ability to synthesise, degrade and release dopamine. Any novel neuronal reactions that were identified by literature review, but not present in Recon2 [50], were either incorporated into Recon3D [35], or added to Recon3D prior to model generation (Table S-10).

#### 2.1.1 Dopaminergic neuronal transporters

To identify transporters specific to dopaminergic neurons, we began with the 1550 human extracellular transport reactions from the Virtual Metabolic Human database [51], which correspond to 255 genes as identified by gene-protein-reaction associations. Extracellular transport genes were manually checked for association with dopaminergeic neurons by manual literature curation. This primarily involved the identification of transporters present in human substantia nigra pars compacta tissue or cell cultures of dopaminergic neurons through *in situ* hybridisation, reverse transcription polymerase chain reaction, immunohistochemistry or immunoblotting. When human data was not found, data from rat or mouse was included instead. Additionally, when data specific for dopaminergic neurons or substantia nigra pars compacta was not found, evidence for transporters being present in neurons in general, astrocytes or in the blood brain barrier was used instead.

#### 2.1.2 Dopamine metabolism

In Recon2 [50], there were already 75 tyrosine-related reactions. This content was extended with dopaminergic neuronal specific information from a comprehensive literature review of dopamine metabolism [52] and additional manual curation of the literature (Table S-3), the results of which were incorporated into Recon3D [35].

### 2.2 Neuronal biomass constituent turnover constraints

#### Summary

Stoichiometric specification of biomass composition [53], as well as cellular synthesis and turnover requirements, is an essential component for the formulation of the objective function in constraint-based modelling. However, fully differentiated dopaminergic neurons do not replicate and therefore, it is sufficient if lipid, nucleic acid, and amino acid synthesis fluxes meet their turnover demand. We adapted an established methodology [54] to define the minimal biomass maintenance and turnover requirements for dopaminergic neurons. This required manual curation of the neurochemical literature to extract (i) neuronal biomass composition, (ii) biomass constituent turnover fluxes, and (iii) key biomass constituent degradation reactions in dopaminergic neurons. This information was then combined to establish the minimum biomass constituent turnover requirements for a dopaminergic neuron, which were then used as constraints during the dopaminergic neuronal model generation process.

##### 2.2.1 Biomass constituent turnover constraints

###### Neuronal biomass composition

The fractional composition (*μmol*/*gDW*) of biomass constituents in a human substantia nigra pars compacta dopaminergic neuron was obtained as follows. First, the percentage composition of lipid and water of a human substantia nigra pars compacta dopaminergic neuron was assumed to be the same as that reported for 55 year human cerebral cortex grey matter, that is 39.6% dry weight of lipids, 60.4% dry weight of non-lipid residues, and 82.3% wet weight water content [55]. Furthermore, we used the protein wet weight (WW) composition for human substantia nigra (99 mg/g WW) [56] to calculate the protein dry weight (DW) composition. RNA and DNA dry weight fractional compositions for human substantia nigra (grey matter) were obtained from the literature (3.29 *μ*g/mg DW of RNA and 1.81 *μ*g/mg DW of DNA) [57]. Based on the relative concentrations of the different neuronal lipids, amino acids, and nucleic acids, the dry weight percentage composition was estimated to be 39.60% lipid, 55.93% protein, 0.18% DNA, 0.33% RNA, and 3.96% others [58]. Using these percentages for each biomass constituent class, the percentage of each biomass constituent was then calculated by assuming that the amount of each constituent, within each constituent class, was the same as that stoichiometrically specified in the Recon3D [35] biomass maintenance reaction (biomass_maintenance_noTrTr).

The fractional composition (*μmol*/*gDW*) of biomass constituents *in vitro* was calculated by assuming that the *in vivo* biomass composition is the same as the *in vitro* biomass composition. The *in vivo* percentage of each biomass constituent was converted into an *in vitro* fractional composition (*μmol*/*gDW*) using

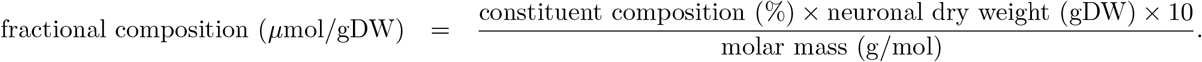

The neuronal dry weight (gDW) was calculated using

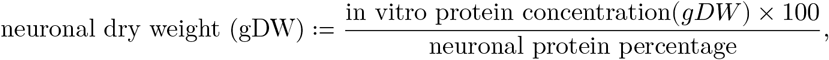

where we measured the *in vitro* protein concentration using a bicinchoninic acid assay (0.0002459 to 0.00047053 *μ*g/cell) and divided it by a count of the number of cell nuclei, leading to an estimated average dry weight of a single dopaminergic neuron to be equal to 6.4 × 10^−10^ (gDW/cell), with a range of 4.4 − 8.41 × 10^−10^, and a neuronal protein percentage of 55.93% [58] was obtained from the literature. The result is a coarse-grained approximation of individual neuronal lipid, amino acid and nucleic acid constituent compositions in *μ*mol/gDW.

###### Biomass constituent turnover fluxes

For each neuronal biomass constituent, half-lives (*hr*^−1^) were collected from the literature [59] and used to calculate the *turnover flux* (Table S-9), defined by

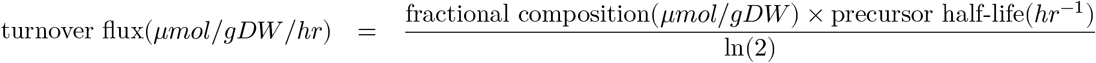

where the fractional composition of key metabolites (*μmol*/*gDW*) was obtained from the literature [57].

###### Biomass constituent degradation reactions

Manual curation of the literature was used to identify the degradation pathway(s) for each biomass constituent and the first reaction in each degradation pathways was identified in Recon3D. For example, as reviewed in [60], phosphatidylserine is exclusively localised in the cytoplasmic leaflet of neuronal and astrocytic membranes, forming protein docking sites for signalling pathways. The phosphatidylserine decarboxylase enzyme is able to decarboxylate the serine moiety of phosphatidylserine to form phosphatidylethanolamine. Although one of the fatty acyl groups of phosphatidylserine can also be hydrolysed to convert phosphatidylserine into lysophosphatidylserine, this is quantitatively a minor pathway [60].

###### Biomass constituent turnover constraints

To enforce biomass constituent turnover for each biomass constituent, constraints were applied on one or more degradation reaction rates. When a biomass constituent was associated with a single degradation reaction, this reaction was set to irreversible in the direction of degradation, and the lower bound was set to 0.75 times the turnover flux. A 25% relaxation of the lower bounds from the estimated degradation rate was used as standard to account for uncertainty in the data [61]. For the example of phosphatidylserine (Section 2.2.1), a lower bound on the degradation rate was implemented by setting a lower bound on the phosphatidylserine decarboxylase reaction (PSDm_hs). When a biomass constituent could be metabolised through a reversible reaction, the turnover constraint was applied to the catabolic direction.

When a biomass constituent could be degraded by more than one reaction, the sum total fluxes of degradation by all degradation reactions was set to be greater than 0.75 times the turnover flux, via an inequality of the form

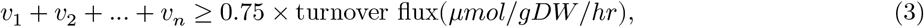

with due consideration of reaction directionality. Support for inequalities such as Equation 3 within constraint-based modelling problems has been fully implemented within the Constraint-Based Reconstruction and Analysis Toolbox COBRA Toolbox [20]. This approach resulted in 21 turnover constraints (Table S-9) on single degradation reactions, and a further 11 turnover constraints each on a set of degradation reactions (Table S-8), when the metabolite could be degraded by more than one pathway.

### 2.3 Active and inactive genes and reactions

A *context-specific metabolic model* should contain only the set of reactions active in a particular context. Therefore, we assembled a core set of genes and metabolic reactions known to be active or inactive in dopaminergic neurons *in vivo* or in dopaminergic neurons in culture. A core set of active genes (Table S-5) and inactive genes (Table S-6), as well as active and inactive reactions (Table S-3 and S-6) and present or absent metabolites (Table S-7), was obtained either from manual curation of the literature or from transcriptomic data (SM5). Manual literature curation was focused on the physiological and biochemical literature on dopamine metabolism, dopaminergic neuronal transporters, central carbon metabolism, mitochondria-associated reactions and genes. In addition, manual curation of the literature was used to determine the need for addition or deletion of external reactions that are required for modelling non-equilibrium steady-state fluxes in dopaminergic neuronal metabolism. The list of genes, established by manual literature curation to be metabolically active, was combined with the aforementioned transcriptomic data and used to generate the context-specific model through gene-protein-reaction associations [54].

## 3 Modelling

### 3.1 Dopaminergic neuronal model generation

Context-specific models were generated from this generic model using the XomicsToModel pipeline [36], a novel, flexible COBRA Toolbox extension enabling modular integration of context-specific data derived from literature curation and multi-omic data, including transcriptomic, metabolomic and proteomic data. A complete description of the XomicsToModel pipeline is provided elsewhere [36]. Briefly, the XomicsToModel pipeline can use two model extraction algorithms, fastcore, an established and widely used algorithm for extracting a minimal flux consistent model, and thermoKernel [36] a novel model extraction algorithm for extracting a minimal thermodynamically flux consistent model. The XomicsToModel pipeline enables flexible specification of technical model extraction parameters, such as the algorithm to use for model extraction, the transcriptomic expression threshold, the magnitude of the largest anticipated flux, the tolerance beneath which a predicted numerical flux is considered zero, whether to close or open unspecified external reactions, whether or not to close ionic transport reactions for sodium, calcium, potassium and iron, as well as solver-specific parameters and debugging options to enable evaluation of the intermediate results of each major consecutive step within the pipeline. In order to identify the optimal technical parameters [62] to extract the iDopaNeuro models with maximum predictive fidelity, an ensemble of context-specific models was generated, by varying all uncertain technical parameters. The key steps are as follows.

#### 3.1.1 Generic model generation

Given the Recon3D reconstruction, plus 21 additional reactions added by subsequent manual literature curation (Table S-10), a generic model of human metabolism was generated by extracting the largest stoichiometrically and flux consistent subset of the reconstruction [20]. This resulted in a generic model of human metabolism, with 10,621 metabolic reactions, 5,835 unique metabolites, and representing the activity of 2,248 open reading frames.

#### 3.1.2 Integration of context-specific data

A metabolic network formed from the set of core reactions alone is not necessarily flux consistent, that is, some reactions may not admit a non-zero steady-state flux. Therefore, we used the thermoKernel algorithm Preciat et al. [36], implemented in the COBRA Toolbox Heirendt et al. [20] to generate a compact, flux and thermodynamic-consistent model. This algorithm returns a minimal number of additional reactions, beyond the core set, which are required to ensure the flux and thermodynamic-consistency of the model. Therefore, the output is a context-specific, flux and thermodynamic-consistent model.

#### 3.1.3 Maximum metabolite uptake constraints

Only the constituents of the fresh medium, plus some reversible extracellular transport reactions including water, carbon dioxide and oxygen, were permitted to be taken up by the model (Table S-1). That is, lower bounds on the corresponding exchange reactions were set by assuming that the maximum uptake rate is equal to the metabolite concentration in the fresh medium, divided by the duration of the interval being modelled (Table S-1). This maximum uptake rate was then used to set the lower bound on the corresponding exchange reaction. This is always an overestimate of the actual metabolite uptake rate, because it effectively assumes that the concentration of each metabolite taken up is zero at the end of the time interval. The fresh culture medium was primarily composed of defined medium, however, certain supplements required for maintenance of neurons in culture consist of proprietary formulations where the concentration of each metabolite is not publicly available. Where certain supplements were known to contain unspecified amounts of key metabolites, the corresponding exchange reactions were opened to uptake only. For example, AlbuMAX™ II Lipid-Rich BSA (ThermoFisher Inc.) is a commercially available serum substitute which was used for maintenance of neuronal cultures. Although its composition is a trade secret, it is known to contain triglycerides and cholesterol [63], and therefore the corresponding exchange reactions were opened to represent growth in cell culture. The lower bounds on all other metabolite uptake reactions were set to zero, to reflect the assumption that no other metabolites were accessible to the *in vitro* culture.

#### 3.1.4 Model extraction

In general, a model extraction algorithm extracts a context-specific subset of a generic model given certain context-specific input data. In our case, this input data were: (i) The set of genes, metabolites and reactions that were identified by manual curation of biochemical literature as being active or inactive in human dopaminergic neurons from the substantia nigra (Section 2), and (ii) RNA-sequencing data from a dopaminergic neuronal *in vitro* cell culture, was mapped into the generic human metabolic model to identify the genes that should be active or inactive in a dopaminergic neuronal reconstruction (Section 1.2), based on quantitative expression levels above or below a user-specified threshold. (iii) Bounds on otherwise reversible exchange reactions that were manually set to satisfy specific characteristics of a (neuronal) cell culture, e.g., the production of oxygen and glucose were disallowed by setting the upper bound on the corresponding exchange reaction to zero, so that only uptake became possible in the model. (iv) A set of reactions associated to dopaminergic neuronal metabolism were qualitatively constrained to make sure that the model could produce neuromelanin, ATP, dopamine and gamma-aminobutyric acid.

#### 3.1.5 Exometabolomic data integration

The XomicsToModel pipeline uses quantitative exometabolomic data (Section 1.3) of fresh and spent media to constrain external reaction fluxes in each candidate dopaminergic neuronal model (Table S-2). This approach assumes that the differentiated dopaminergic neurons are (a) not growing, and (b) at a metabolic steady state. Our justification is that, in contrast to earlier stages, the cell number does not alter significantly in the last five days in culture (<3-4% increase, Supplementary Fig. 20). Also, it is known that the rate of neuronal differentiation reaches a plateau toward the end of the period in culture [18], consistent with our microscopic observations. Furthermore, the ratio of intracellular cell volume to extracellular media in macroscopic cell culture is sufficiently large that metabolite concentration changes in the surrounding medium are ameliorated.

By assuming that metabolite exchange fluxes are constant with respect to time, between replacement with fresh medium and collection of spent medium, the measured rate of exchange is the difference between spent and fresh medium concentrations, divided by the time interval the fresh medium was exposed to the cells before being sampled. In the model, the unit of flux is *μ*mol/gDW/hr, while the unit of metabolite concentration change is *μ*mol/L. In order to transform an extracellular metabolite concentration change into a mean exchange reaction flux, we used

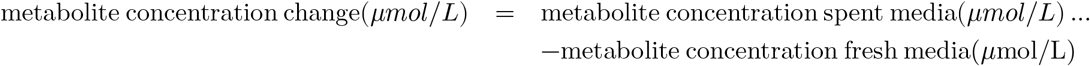

and

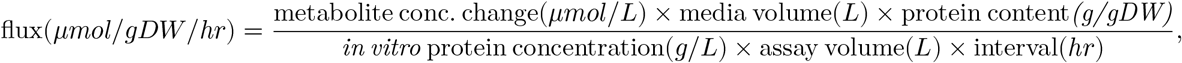

where cell dry weight was estimated as described in Section 2.2.1. By convention, a negative flux value represents uptake and a positive value represents secretion.

The measurements of metabolite concentrations and the measurement of cell culture parameters, e.g., protein concentration, are associated with measurement uncertainty. This measurement uncertainty was propagated to represent each measurement with a mean and standard deviation. The result is a vector of mean measured exchange reaction fluxes *ν*_*exp*_ ∈ ℝ^*n*^ accompanied by a standard deviation vector *σ*_*exp*_ (Eq. (5)). Both of these vectors are incorporated into the mathematical formalism to fit model bounds to measured exchange fluxes. However, they cannot be directly incorporated to an any subset of a generic model with arbitrary exchange reaction bounds, because, either due to model misspecification or experimental error, *ν*_*exp*_ may not be consistent with the feasible set of steady state fluxes, as defined in Fig. 3. Specifically, it may be inconsistent with the set defined by the steady state constraint and the reaction bounds, or the set defined by the coupling constraints and the reaction bounds.

**Figure 3:** Amalgamation of mathematical modelling approaches. This optimisation problem is an amalgamation of all of the objectives and constraints used by each model derived from the context-specific model ensemble, the validation of the iDopaNeuro models and their prospective applications. The variables are net *ν* ∈ ℝ^*n*^, forward *ν*_*f*_ ∈ ℝ^*n*^ and reverse *ν*_*r*_ ∈ ℝ^*n*^ internal reaction flux as well as external reaction net flux *w* ∈ ℝ^*k*^. In the objective the data are linear objective coefficients *a* ∈ ℝ^*n*^ on net fluxes, weights *q* ∈ ℝ^*n*^ on the *p*-norm of internal net fluxes and measured exchange reaction fluxes *w*_*exp*_ ∈ ℝ^*k*^ (for a subset of exchanges). Note that here, in a slight departure from established notation, ‖·‖_*p*_ denotes the component-wise norm, and ln the component-wise natural logarithm. That is, when *p* = 0, 1, 2 the *p*-norm denotes the zero-, one- and two-norm, respectively. The entropy function is −*x*^*T*^ ln *x*, so entropy maximisation is minimisation of *x*^*T*^ ln *x*. The ◊ denote the objective terms that represent maximisation of flux entropy. The constraint data are the internal *N* ∈ ℤ^*m*×*n*^ and external *B* ∈ ℤ^*k*×*n*^ reaction stoichiometric matrix, while the data *C* ∈ ℤ^*s*×*n*^ and *d* ∈ ℤ^*s*^ enforce coupling between net reaction fluxes, e.g., to represent a constraint on cumulative turnover of a metabolite by a set of degradation reactions. The other constraints are bounds on net internal flux, net external flux and the bounds on each of the variables.

Therefore, we assumed that inconsistency would occur and fit the bounds of the candidate dopaminergic neuronal model to the experimental data while allowing relaxation of the bounds on net flux in Fig. 3, using the following quadratic optimisation problem:

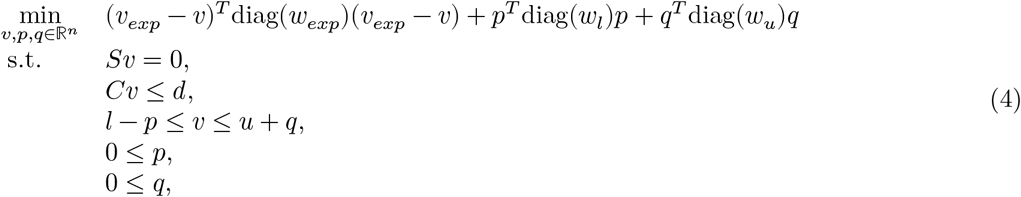

where *S* ≔ [*N, B*], *ν* ≔ [*ν*_*f*_ − *ν*_*r*_; *w*], *l* ≔ [*l*_*ν*_; *l*_*w*_] and *u* ≔ [*u*_*ν*_; *u*_*w*_] in Fig. 3 and where 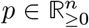 and 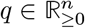 are non-negative variables that permit relaxation of the lower and upper bound constraints, respectively.

Problem 4 always admits a steady state flux *ν* ∈ ℝ^*n*^ and also allows for different weights to be input as parameters. With 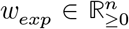 to penalise deviation from experimentally measured mean fluxes, 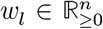 to penalise relaxation of lower bounds, and 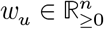 to penalise relaxation of upper bounds.

For example, we set the penalty on deviation from experimental measurement to be the inverse of one plus the variance:

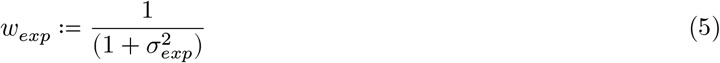

where 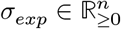 is the standard deviation. This approach increases the penalty on deviation from an experimentally measured mean flux where the variance is lower. Note that, certain lower or upper bounds might not be realistic to be relaxed, e.g., an essential amino acid can always be taken up but never secreted, therefore the upper bound on the corresponding exchange reaction must be zero. This can be specified a priori, using the technical parameters input to the XomicsToModel pipeline.

#### 3.1.6 ATP maintenance requirements

At a steady state, the rate of ATP production equals the rate of consumption. The ATP maintenance reaction rate (ATPM) was estimated by assuming that nigrostriatal dopaminergic neurons have both a basal and an electrophysiological signaling-related ATP requirement. The residual energy consumption rate is the ATP requirement in the absence of electrophysiological activity and can be obtained by blocking the voltage-gated sodium-potassium pump or inducing a coma [64]. Basic cell activities are not tightly coupled to electrophysiological signaling, therefore, they can be sustained by residual energy consumption. This energy is required for mitochondrial proton exchange, axoplasmic transport, and the biomass maintenance of lipids, oligonucleotides, or proteins. The lower bound on ATPM represents the residual energy consumption rate of human grey matter, which has been estimated to be 186 *μmol*/*gDW* /*hr* [64] (Table S-9). This, however, does not account for electrophysiological signaling-related ATP consumption.

### 3.2 Prediction of reaction fluxes

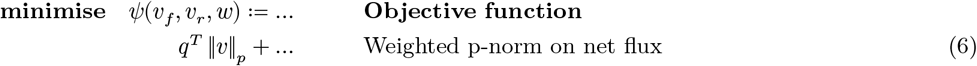

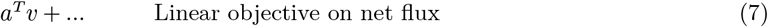

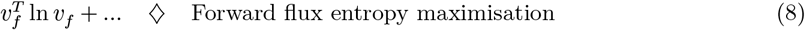

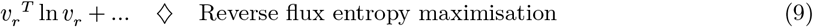

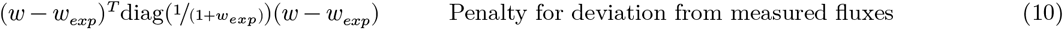

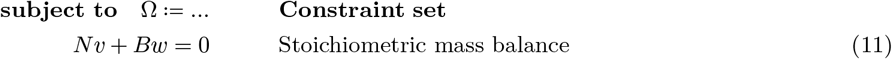

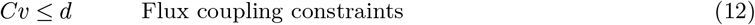

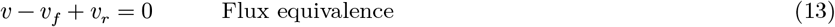

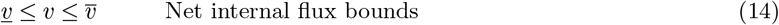

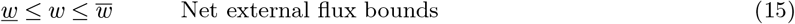

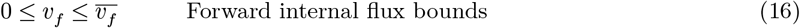

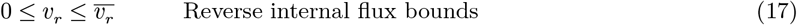

Several mathematical modelling innovations were necessary to adapt constraint-based modelling to predict reaction fluxes in non-replicating cells, such as neurons, where maximisation of growth rate is not appropriate. Fig. 3 provides an amalgamation of the mathematical modelling approaches we employed when predicting reaction fluxes. Many of these approaches were already implemented within the COBRA Toolbox, and any new approaches were implemented within it. We generated an ensemble of flux predictions by applying a set of different objectives (Table 1) to each model in the aforementioned ensemble of context-specific models.

**Table 1.** Candidate objective functions. A set of candidate objective functions were used to predict net reaction fluxes. sup(*x*) denotes the supremum of *x* and □ denotes the set of external reactions. Note that here, in a slight departure from established notation, ‖·‖_*p*_ denotes the component-wise norm, and log the component-wise logarithm.

| Objective | Data | Objective, $\psi$ |
| --- | --- | --- |
| Min. zero-norm of net flux | $p = 0, q = 1$ | $\psi_0 = 1^T \ v\ _0$ |
| Min. weighted zero norm of net flux | $p = 0, q = \begin{cases} x_j \in \mathbb{R} & -\frac{\log(x)}{\log(\sup(x))} \\ x_j \notin \mathbb{R} & 0 \end{cases}$ | $\psi_0(q) = q^T \ v\ _0$ |
| Min. one-norm of net flux | $p = 1, q = 1$ | $\psi_1 = 1^T \ v\ _1$ |
| Min. weighted one-norm of net flux | $p = 1,$<br>$q = \begin{cases} x_j \in \mathbb{R}, & \log\left(\frac{\sup(x)}{x}\right) \\ x_j \notin \mathbb{R}, j \notin \square, & \text{mean}\left(\log\left(\frac{\sup(x)}{x}\right)\right) \\ x_j \notin \mathbb{R}, j \in \square, & 0 \end{cases}$ | $\psi_1(q) = q^T \ v\ _1$ |
| Min. two-norm of net flux | $p = 2, q = 1$ | $\psi_2 = 1^T \ v\ _2$ |
| Min. weighted two-norm of net flux | $p = 2, q \propto 1/\text{expression}$ | $\psi_2(q) = q^T \ v\ _2$ |
| Max. entropy of forward and reverse flux | $p = 0, q = 0$ | $\psi_v = v_f^T \ln v_f + v_r^T \ln v_r$ |

**Table 2.** Evaluation of predictive fidelity of different objectives, averaged over the model ensemble. Variation of model extraction parameters resulted in generation of 128 models. From each of these models, an uptake constrained model modelUpt and a secretion constrained model modelSec were generated to predict measured secretion and uptake reaction fluxes, respectively. With each model, reaction fluxes were predicted using 7 different objectives. The average qualitative, semi-quantitative and quantitative predictive fidelity, over all 256 = 2 × 128 models, is given.

| Objectives | Correct/total predictions | Spearman $\rho$ | Euclidean distance score | Average score |
| --- | --- | --- | --- | --- |
| $\psi_v = v_f^T \ln v_f + v_r^T \ln v_r$ | 0.72559 | 0.54587 | 0.6022 | 0.62455 |
| $\psi_1(q) = q^T \ v\ _1$ | 0.43919 | 0.6162 | 0.78005 | 0.61181 |
| $\psi_1 = 1^T \ v\ _1$ | 0.39719 | 0.5888 | 0.79773 | 0.59457 |
| $\psi_2(q) = q^T \ v\ _2$ | 0.71875 | 0.54776 | 0.35054 | 0.53902 |
| $\psi_2 = 1^T \ v\ _2$ | 0.75133 | 0.50338 | 0.31766 | 0.52413 |
| $\psi_0 = 1^T \ v\ _0$ | 0.32369 | 0.6107 | 0.47556 | 0.46998 |
| $\psi_0(q) = q^T \ v\ _0$ | 0.58935 | 0.28903 | 0.17235 | 0.35024 |

#### 3.2.1 Evaluation of predictive fidelity

Each model within the ensemble of context-specific models, was generated with quantitative exometabolomic data on metabolites taken up and secreted by midbrain-specific dopaminergic neurons in culture, represented by constraints on a subset of external reaction fluxes. An underestimate of the predictive fidelity of each of model was obtained by generating a pair of derived models. First, an uptake constrained model (modelUpt) was derived by replacing quantitative bounds on secreted metabolites with arbitrarily large reversibility constraints. Second, a secretion constrained model (modelSec) was derived by replacing quantitative bounds on up-taken metabolites with arbitrarily large reversibility constraints. A metabolite was defined as secreted (uptaken) if the mean measured exchange, minus (plus) standard deviation, was greater (less) than zero. A metabolite was defined as unchanged (i.e. neither taken up nor secreted) if the magnitude of the mean was less than or equal to the standard deviation.

Setting arbitrarily large reversibility constraints includes removal of constraints forcing uptake or secretion of measured metabolites, which were previously applied while the models were being generated by the XomicsToModel pipeline (Section 3.1.4). That is, these constraints were replaced by lower and upper bounds set to the default maximum magnitude flux, e.g. [-1e5, 1e5] *μmol*/*gDW* /*hr*, before the predictive capacity of the models were tested. To put it another way, constraining uptake or secretion during model construction ensures that a model can uptake or secrete, but once such constraints are removed and replaced with reversibility constraints, only the set of uptakes, or secretions, together with the chosen objective determines whether a metabolite is actually secreted, or taken up, respectively.

Then each pair of models, modelUpt and modelSec, was combined with each objective and the predicted flux was evaluated for its qualitative and quantitative ability to predict the fluxes of measured secretion and uptake reactions, respectively. Predicted exchange flux was considered zero if it was less than the numerical tolerance of the solver (1*e* − 6). Qualitative accuracy was measured as the fraction of of correct predictions of uptake or secretion. Semi quantitative accuracy was measured by the Spearman rank correlation coefficient between predicted and all measured exchange fluxes. Quantitative predictive fidelity was initially measured by a weighted Euclidean distance between predicted and all measured exchange fluxes, specifically

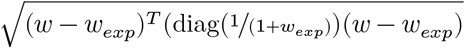

where *w* and *w*_*exp*_ denote predicted and mean measured exchange flux. Subsequently, this metric was converted into a measure of quantitative predictive accuracy in a unit interval for each prediction that was given by the descending rank order of the weighted Euclidean distance, divided by the number of such predictions. The average of qualitative, semi-quantitative and quantitative predictive accuracy was used to define the average accuracy of each prediction. The average accuracy over all models was used to identify the objective with the highest predictive fidelity. The average accuracy over all models, using only the objective with the highest predictive fidelity, was used to identify iDopaNeuro models from the ensemble with the highest predictive fidelity.

#### 3.2.2 Model Validation

The effects of three different perturbations to the iDopaNeuro models were predicted: (i) inhibition of mitochondrial complex V, (ii) inhibition of mitochondrial complex I, and (iii) replacement of glucose with galactose as the primary sugar source. The control for these experiments was considered to be culture within the *differentiation medium without PMA*, which contains glucose as the main carbon source, as described in Subsection *1*.*1*.*1*. To predict control and perturbed fluxes, flux entropy maximisation was used as the objective for internal reactions, while exchange reaction bounds derived from exometabolomic data were replaced with the more relaxed generic bounds from the same reactions in Recon3, and a quadratic term was added to the objective to penalise deviation from each measured exchange flux (Fig 3), except for one measured flux left out within each iteration of a leave-one-out cross validation. Where an exchange flux was not acquired for a metabolite in a perturbed experimental condition, the corresponding measured exchange flux from the control experimental condition was used, were it was acquired. Inhibition of mitochondrial complex I and V were simulated by setting the bounds on the ATP Synthase (ATPS4m) and Mitochondrial NADH Dehydrogenase (NADH2_u10m) reactions to zero, respectively. Measured and predicted metabolomic exchange were compared qualitatively, semiquantiatively and quantitatively, as described previously. The consequences of deleting the glucocerebrosidase (GBA1) gene, the gene most commonly associated with PD, was also predicted, for validation in an independent study.

#### 3.2.3 Prospective research design

All reactions in the iDopaNeuro models are thermodynamically flux consistent, so all metabolites should be considered as targets for future development of quantitative metabolomic platforms for characterisation of intracellular dopaminergic neuronal metabolism. However, in practice, a prioritised list of metabolites exchanged with the environment would enable optimal design of exometabolomic platforms targeted to a manageable subset of metabolites. Similarly, under constrained external reactions represent an opportunity for further reconstruction efforts, or refinement. Therefore, we developed a novel *uncertainty reduction* pipeline (Fig. 31) that predicted the metabolites whose corresponding external reactions would be the most important to constrain in future research, in order to maximally shrink the feasible set of external reaction fluxes of the iDopaNeuro models.

First, uniform sampling [65] was used to obtain an unbiased assessment of the fluxes satisfying the constraints on the solution space, *ψ* = 0 and Ω ≔ {*l*_*c*_ = *u*_*c*_ = 0} in Fig. 3. Uniform sampling was implemented using the Riemannian Hamiltonian Monte Carlo (RHMC) algorithm, within the COBRA Toolbox [20], a Markov chain that, at each step, picks a random direction and uses an Ordinary Differential Equation to move along a curve determined by the starting point and direction. RHMC is able to take longer steps in polytopes than traditional random walks such as the ball walk and hit-and-run [66]. The parameters used were *nSamples* = 8 × dim(Ω), which represent the total number of samples returned. This resulted in a set of *z* steady state flux vectors, *V* ∈ ℝ^*n*+*k*×*z*^, from which a covariance matrix, *Q* ∈ ℝ^*k*×*k*^, restricted to the set of external reactions, was computed. The first most informative external reaction was defined to be the one corresponding to the largest Euclidean norm of this covariance matrix. Thereafter, we used a heuristic, iterative method that greedily selected the row of the covariance matrix that had the maximum Euclidean distance to the subspace spanned by the rows selected already. This approach ensures that the variance reduction due to cumulative measurement of higher-ranked exchange reactions was taken into account in the ranking of subsequent most informative metabolites.

## Results

### 4 Experiments

#### Summary

A human neuroepithelial stem cell line from a healthy donor was maintained and differentiated into dopaminergic neurons (Section 4.1), identified by immunoreactivity for tyrosine hydroxylase indicated the presence of neurons capable of converting tyrosine to L-DOPA (Fig. 4a), the penultimate step in dopamine synthesis. RNA sequencing and data processing quantified the expression of 8,204 genes but only 820 corresponded to genes present in the generic model of human metabolism. All genes expressed above a threshold were considered active, and the rest considered inactive, unless manual curation of the literature dictated otherwise for the iDopaNeuroCT model, while giving preference to our *in vitro* experimental data in the iDopaNeuroC model.

**Figure 4:**
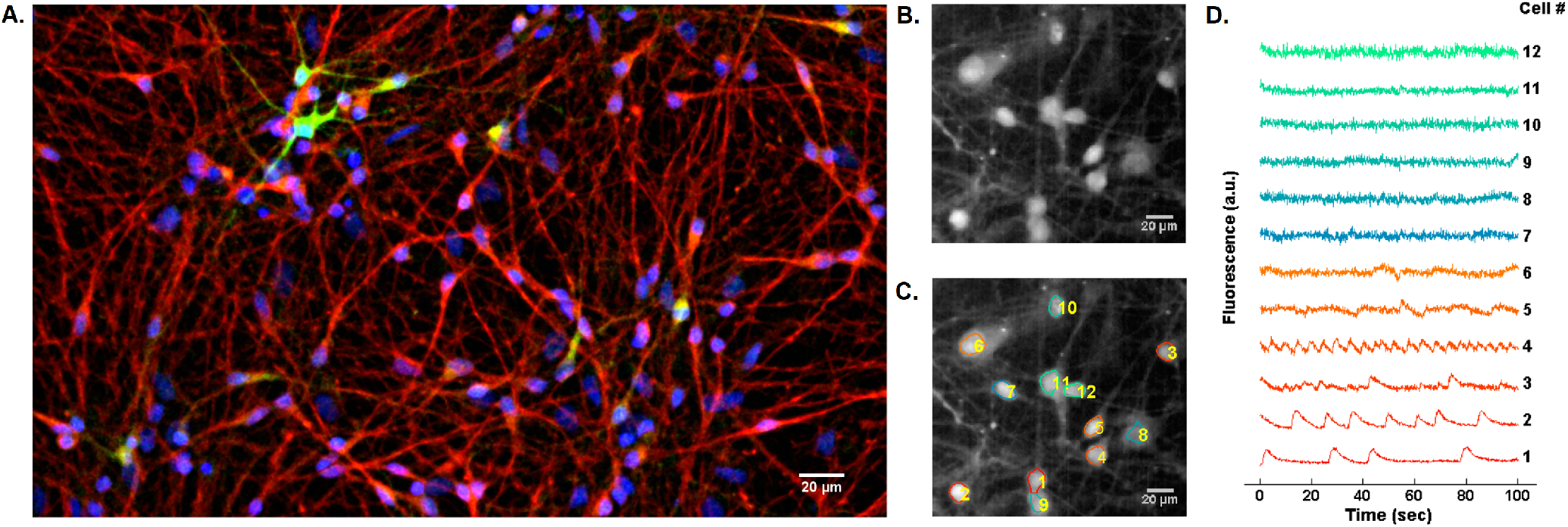
Immunostaining and calcium imaging. Immunostaining of differentiated neurons and calcium imaging of spontaneously firing human neuroepithelial stem cell differentiated into dopaminergic neurons. (**A**) Immunostaining of a representative well at day 23, showing neurons positive for nuclei with Hoechst (blue), TUB*β*III (red) and TH (green); scale bar 20*μ*m. (**B**) Mean frame of a field of view of representative neurons. (**C**) Automatic segmentation of neurons. (**D**) Fluorescence traces showing the spontaneous activity of individual segmented neurons.

##### 4.1 Cell culture

A human neuroepithelial stem cell line from a healthy donor was maintained and differentiated into dopaminergic neurons, using an established protocol [18] (Section 1.1.1). The differentiation of human neuroepithelial stem cells into neurons was verified by identified by immunofluorescent staining for TUB*β*III. At 23 days of the protocol (Fig. 2), neurons positive for tyrosine hydroxylase, the penultimate step in dopamine synthesis, confirmed the presence of neurons capable of converting tyrosine to L-DOPA, (Fig. 4a). The percentage of tyrosine hydroxylase positive cells was between 15-20%, depending on the well. Furthermore, analysis of calcium imaging data revealed spontaneously electrophysiologically active neurons, that is, neurons firing without a requirement for extrinsic electrophysiological stimulation (Fig. 4b, c, d).

##### 4.2 Transcriptomic analysis

In the transcriptomic data of the differentiated midbrain-specific dopaminergic neuronal culture, fragments were detected from 18,530 genes, but only 8,204 of these were sufficiently abundant to be considered expressed. That is, above a threshold of one Fragment Per Kilobase of exon per Million reads (FPKM) [67]. Of the expressed genes, 1,110 could be mapped to metabolic genes in Recon3D and were considered active, unless manual curation of the literature revealed otherwise for the iDopaNeuroCT model, while giving preference to experimental data in the iDopaNeuroC model. Finally, 790 genes were present in the iDopaNeuroC model and 621 genes for the iDopaNeuroCT model. Transcriptomic data contains a range of gene expression values. Some of the low expression values are attributable to experimental noise or aborted transcripts, but for borderline expression values, it is a challenge to divide the corresponding genes into expressed or not expressed. By default, each gene with less than zero FPKM, on base-two logarithmic scale, was considered not expressed [67]. Each gene with FPKM higher than this threshold was considered expressed. Out of the 18,530 unique genes with expression levels reported in the transcriptomic data, 8,204 were considered to be expressed, based on the aforementioned threshold. However, only 1,110 were mapped into Recon3D (metabolic genes) and therefore included in the model.

To test the viability of the selected transcriptomic data expressed in the *in vitro* culture and selected in Recon3D, a receiver-operating characteristic curve [68] was generated to qualitatively compare the expressed and not-expressed assignments from our transcriptomic data on dopaminergic neurons, against the active and inactive assignments for manually curated dopaminergic neuronal genes (cf. Section 2.3 below), which we assume to be a true representation of dopaminergic neuronal gene expression (Fig. 5). If a gene was considered to be active by manual literature curation and was also found to be expressed in transcriptomic data, it was considered a true positive. The true and false positive fluxes can vary depending on the threshold applied to distinguish between a gene that is expressed or not. In the reconstruction, genes expressed above the threshold were assigned to be metabolically active and genes expressed below the threshold were not included as in the model, unless the corresponding reactions had to be included to generate a flux consistent model.

**Figure 5:**
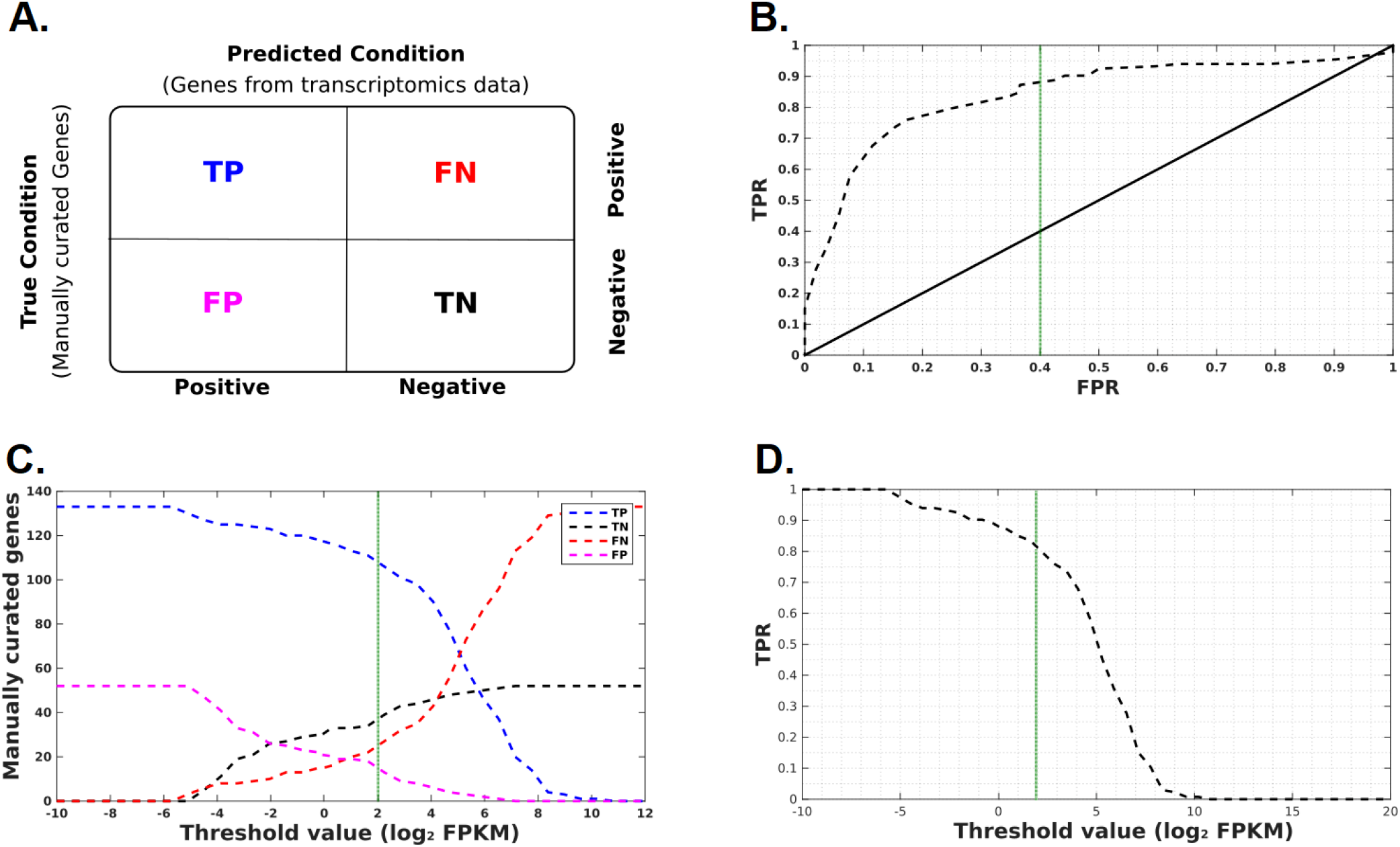
Manually curated genes compared with transcriptomic data. (**A**) Confusion matrix illustrating the performance of the transcriptomic classification into active and inactive genes. TP - True Positive, TN - True Negative, FN - False Negative, FP - False Positive. (**B**) Receiver operating characteristic (ROC) curve. True 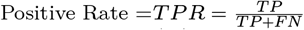, False 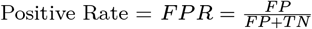. (**C**) Number of manually curated genes per threshold for each condition. (**D**) A true positive rate of 0.9 corresponded to a threshold value of zero Fragments Per Kilobase of transcript per Million mapped reads (FPKM), on base-two logarithmic scale (green vertical line).

##### 4.3 Analytical chemistry

Fresh and spent cell culture media samples were analysed using four complementary mass spectrometry platforms (three different LC-MS platforms and one GC-MS platform), resulting in quantification of a total of 49 metabolites (Fig. 6). Of the 49 metabolites, 6 were quantified by DmPABr-2D [47], 18 from BzCl [45] 17 from GC-MS [46] and 8 were quantified by Accq-Tag [69]. Where platforms measured the same metabolite, the measurement from the platform with the lowest relative standard deviation was employed for comparison with model predictions.

**Figure 6:**
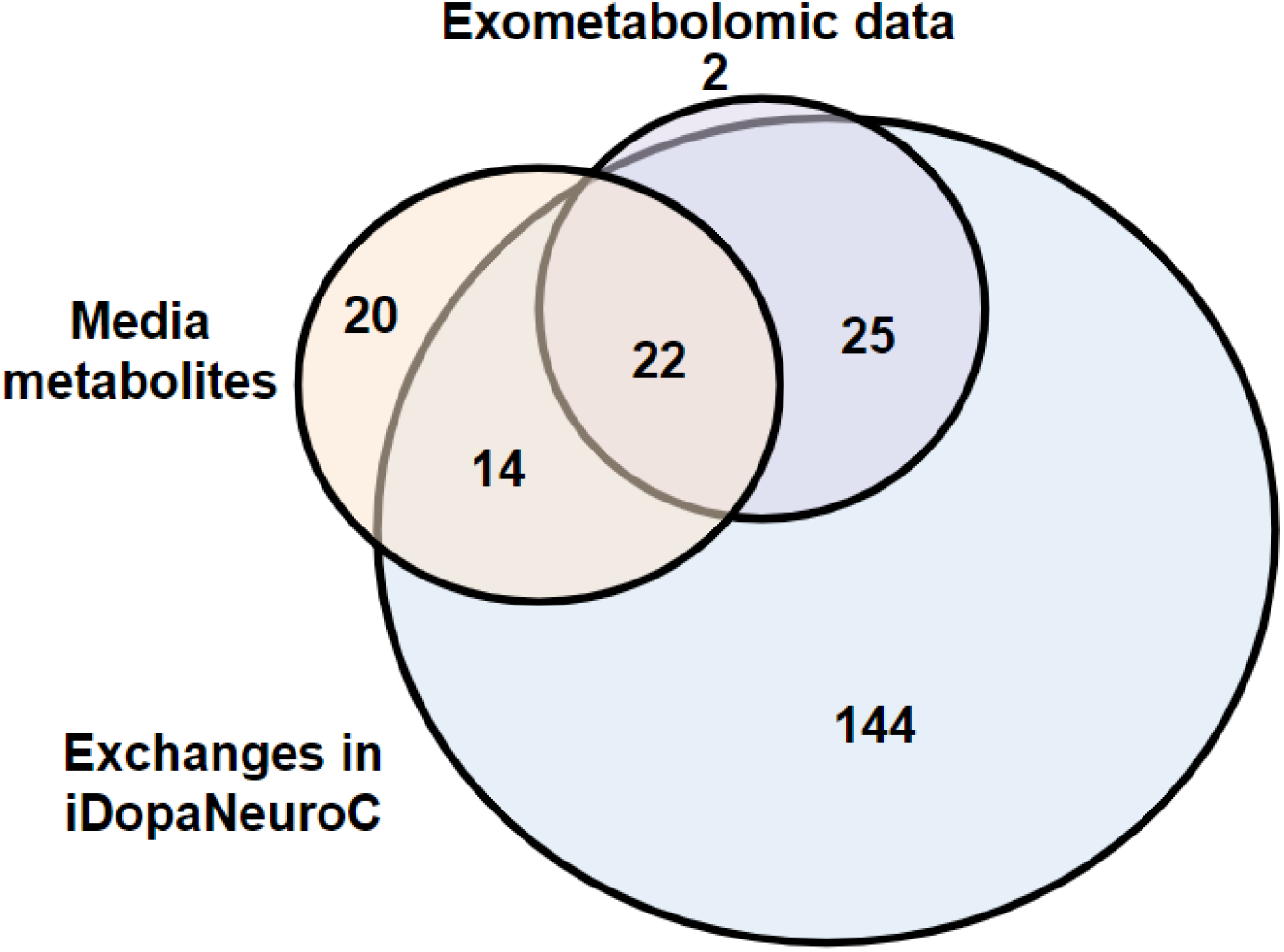
Venndiagram summarising metabolomic measurements. A total of 49 unique metabolites were targeted by the selected metabolomic platforms (purple). Of these, 47 could be used to constrain the model as 2 metabolites (glutarate, methylmalonate) were not present in the stoichiometrically and thermodynamically flux consistent subset of Recon3D. Of these 47 metabolites, 22 were present in the fresh medium (orange) and 25 were synthesised by the cells and secreted into the spent medium. The iDopaNeuroC model contains exchange reactions for 205 metabolites (blue), therefore there still remains 144 metabolites to target with additional exometabolomic platforms.

###### 4.3.1 Metabolomic analysis of fresh medium

We compared measured metabolite concentrations with manufacturers specification of fresh media. The manufacturers specification identifies a total of 57 different metabolites and molecules in the fresh culture medium: 24 amines, 12 vitamins, 16 inorganic salts, 1 lipid, 2 nucleotides and 2 organic acids (Table S-1). Of the fresh medium metabolites and molecules almost all (50/57) were present in the stoichiometrically and flux consistent subset of Recon3D. The remaining 7 molecules were omitted from further consideration as they were inorganic salts or were not present in Recon3D, and a three others (magnesium, cyanocobalamin and selenium trioxide) did not correspond to any stoichiometrically and flux consistent reaction. Analysis of fresh medium samples in multiple GC-MS and LC-MS platforms enabled measurement of the absolute concentrations of glucose and pyruvic acid and 22 of the 24 amines, known to be in the medium. This enabled us to test the concordance between the specifications of the medium manufacturer and the actual concentrations (Fig. 7). Reduced glutathione and L-cystine are two amines that could not be detected by the LC-MS platform.

**Figure 7:**
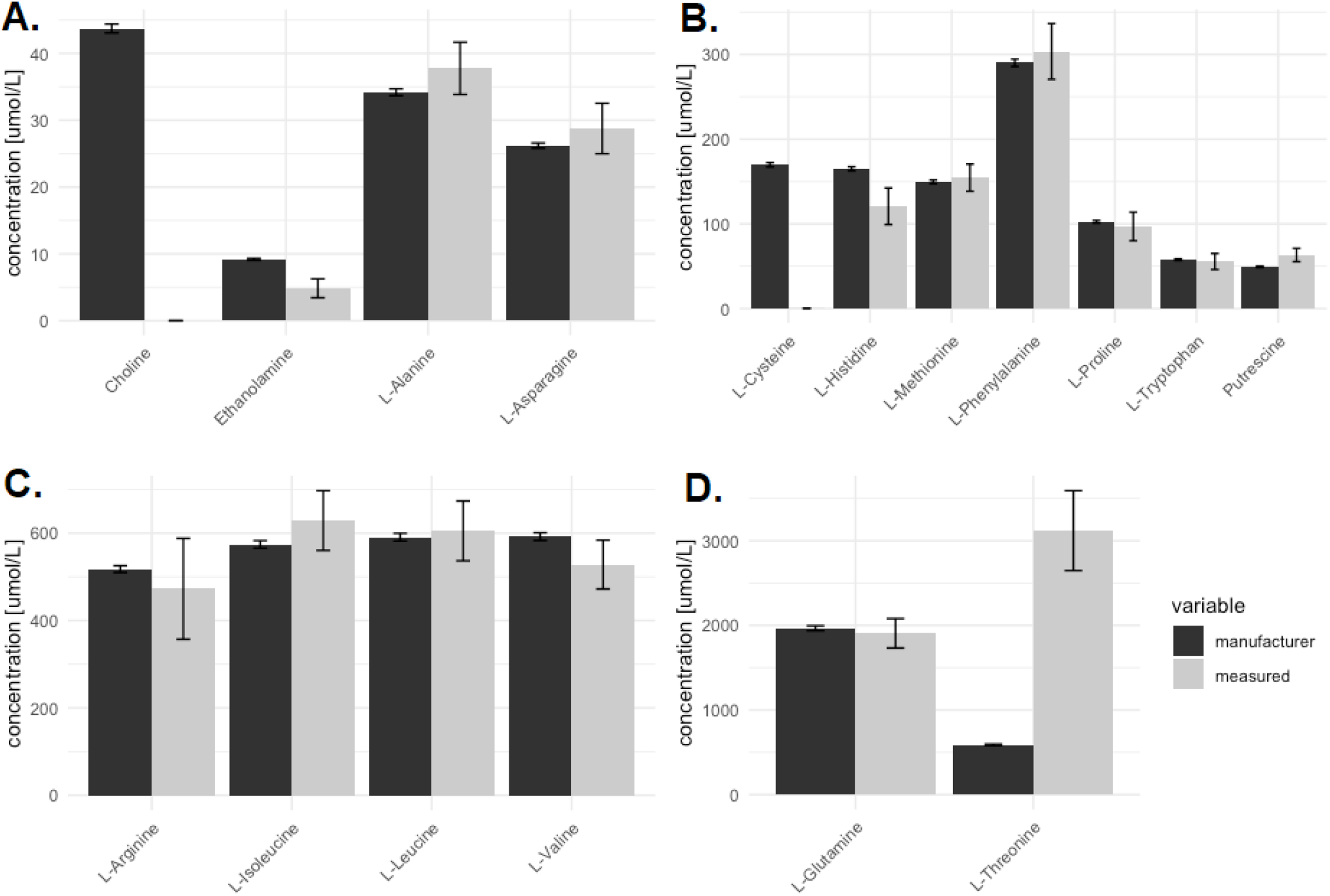
Validation of specified fresh medium concentrations. Metabolite concentrations specified by the medium manufacturer (blue) compared to the absolute concentrations measured by mass spectrometry (grey) for different concentration ranges (A-D). Some quantified metabolite concentrations, e.g., for cysteine, pyruvic acid, valine, aspartic acid and putrescine, appear to significantly deviate from the manufacturers specifications.

**Figure 8:**
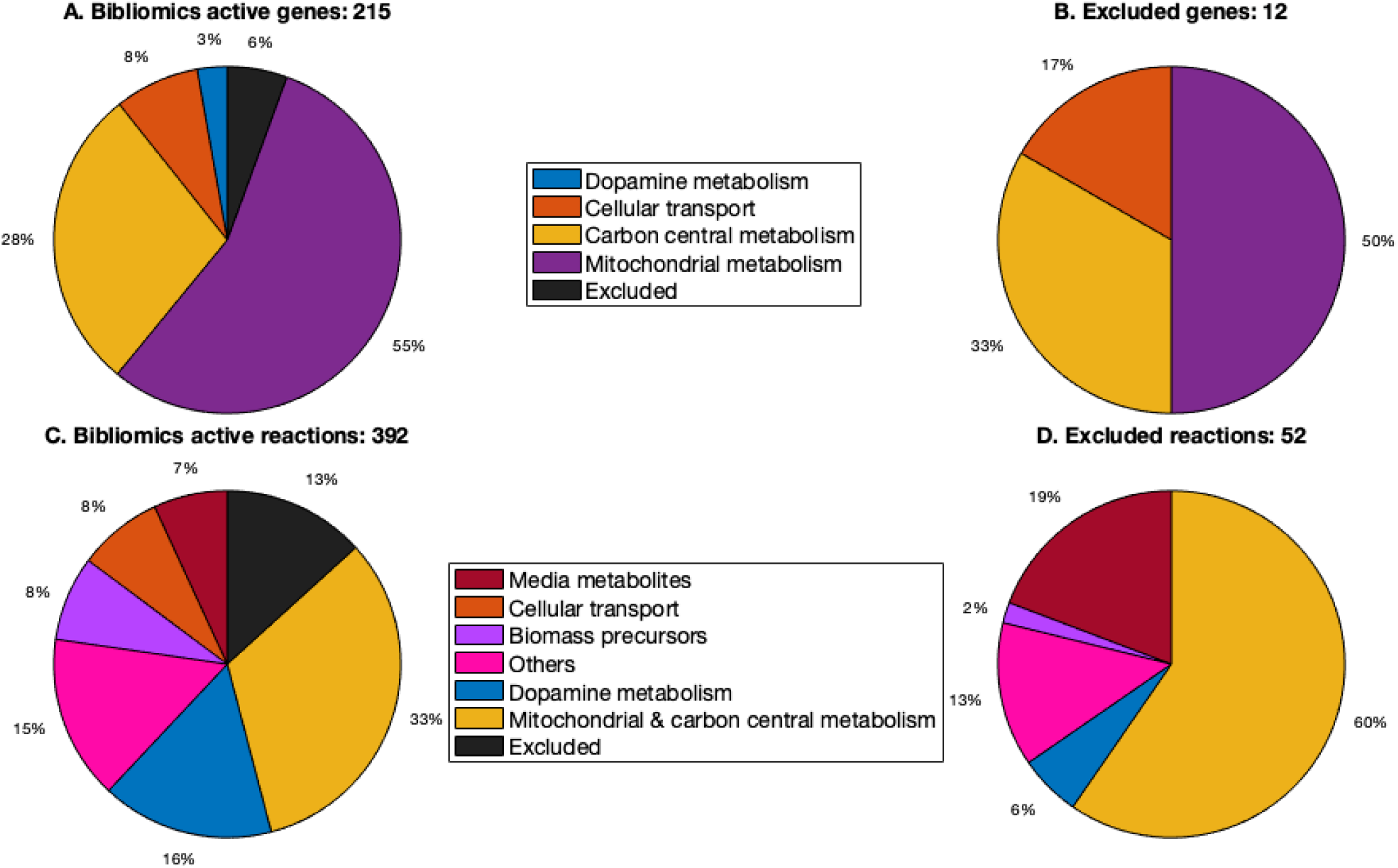
Classification of active reactions and genes by manual literature curation. Classification of active genes (**A**) and reactions (**C**) is partly a reflection of the availability of biochemical information on certain pathways, e.g., central metabolism, and partly a reflection of the pathways that were targeted for curation, e.g., dopamine metabolism. Classification of the subset of the active genes that were excluded (**B**), and the subset of active reactions that were excluded (**D**) from the context-specific dopaminergic neuronal model as they did not correspond to a stoichiometrically and thermodynamically flux consistent subset of Recon3D.

## 5 Reconstruction

Following manual curation of the literature [52, 70, 71, 72, 73, 74, 75, 76, 77, 78], 9 metabolites and 49 reactions were added to the subsystem dopamine metabolism within Recon3D. These are 11 transport, 11 exchange, 19 internal and 8 demand reactions. In Recon3D, dopamine metabolism includes 122 reactions in total (Supp. Fig. 21). Out of these 122 reactions, we were able to collect evidence for the occurrence of 77 reactions in dopaminergic neurons that were also specified as active reactions: 42/49 newly added reactions and 35/73 dopamine-related reactions already present in Recon 2.04. For many reactions (45/122) no clear information was found, therefore they were not specified as active (Table S-3) or inactive (Table S-4).

Manual literature curation established the activity of 239 genes in dopaminergic neurons. In Recon3D, each gene is associated with one or more reactions. Out of these 239 genes, 24 were not associated with any stoichiometric and flux consistent reaction in Recon3D, so were excluded from further analyses (Table S-6 and Fig. 28). Significant effort was made to manually curate transport reactions as their presence or absence help to establish the idiosyncratic boundary conditions for any particular cell type. Accordingly, literature evidence established the presence of 20 transport reactions in human, mouse, or rat substantia nigra. Manual literature curation revealed that 334 unique reactions should be active in dopaminergic neurons of which 68 are specific for dopamine metabolism. Nevertheless, 2 of the active reactions had to be excluded from the model generation process as they were not associated with any stoichiometric and flux consistent reaction in Recon3D (Table S-3 and Fig. 28).

Based on literature curation, a total of 61 genes were deemed to be inactive in dopaminergic neurons and these genes corresponded to 34 inactive reactions (Table S-6). A reaction uniquely encoded by an inactive gene was removed from the model, unless that gene participated in a reaction that was biochemically established to be active. A total of 233 metabolic reactions were specified to be inactive in the brain (Table S-4) and were therefore excluded from the model. Taken together, manual literature curation revealed evidence for the activity, or inactivity, of 300 metabolic genes (Table S-5 and S-6) and 567 metabolic reactions (Table S-3 and S-4) in dopaminergic neurons. Manual literature curation of neuron-specific biomass maintenance requirements resulted in 32 turnover constraints, each of which was either on a single degradation reaction or a set of degradation reactions, when the metabolite could be degraded by more than one pathway.

## 6 Modelling

### 6.1 Identification of condition-specific and cell-type specific dopaminergic models

To identify the best approach to construct a dopaminergic neuronal metabolic model, an ensemble of 135 candidate dopaminergic neuronal metabolic models was generated from a generic model of human metabolism, by varying the parameters of a novel model generation pipeline, XomicsToModel [36], that was supplied with the results of manual literature curation, as well as our *in vitro* transcriptomic and metabolomic data from midbrain-specific dopaminergic neurons. An underestimate of the predictive fidelity of each model was obtained by evaluating the predictive fidelity of a pair of derived models, an uptake-constrained model and a secretion-constrained model, without exometabolomically derived constraints on secretion and uptake reactions, respectively.

To identify a suitable metabolic objective, a set of 7 candidate cellular objectives (Table 1) were optimised on each uptake constrained model and each secretion constrained model, subject to additional constraints representing steady state mass balance, biomass turnover and reaction directionality (Fig. 3). The effects of changing cellular objectives on predictive fidelity, averaged over all 135 models, was evaluated qualitatively (correct/total), semi-quantitatively (Spearman rho) and qualitatively (weighted Euclidean distance). Maximisation of the entropy of all internal forward and reverse fluxes (*ψ*_*ν*_ in 1 and Ω ≔ {*c* = 0, *c*_0_ = 0} in Eq. 3), averaged over all models, was identified as the best performing objective (Supplementary Table 2).

Using maximisation of flux entropy as the objective, the effect of changing pipeline parameters on predictive fidelity, was evaluated qualitatively, semi-qualitatively and qualitatively. If a gene is specified to be active, then requiring that at least one corresponding reaction to be active (1-rxn), rather than all corresponding reactions be active (all-rxn) results in smaller dopaminergic neuronal models (Fig. 9C), with greater qualitative (Fig. 9A) and quantitative accuracy (Fig. 9B). Ensuring that context-specific models are thermodynamically flux consistent (thermoKernel), rather than just flux consistent (fastCore, [79]) results in models of comparable size (Fig. 9C) with an increase in semi-quantitative predictive accuracy (in Fig. 9B,E) as does specifying that all reactions corresponding to genes expressed below a transcriptomic threshold should be inactive (Fig. 9A,B).

**Figure 9:**
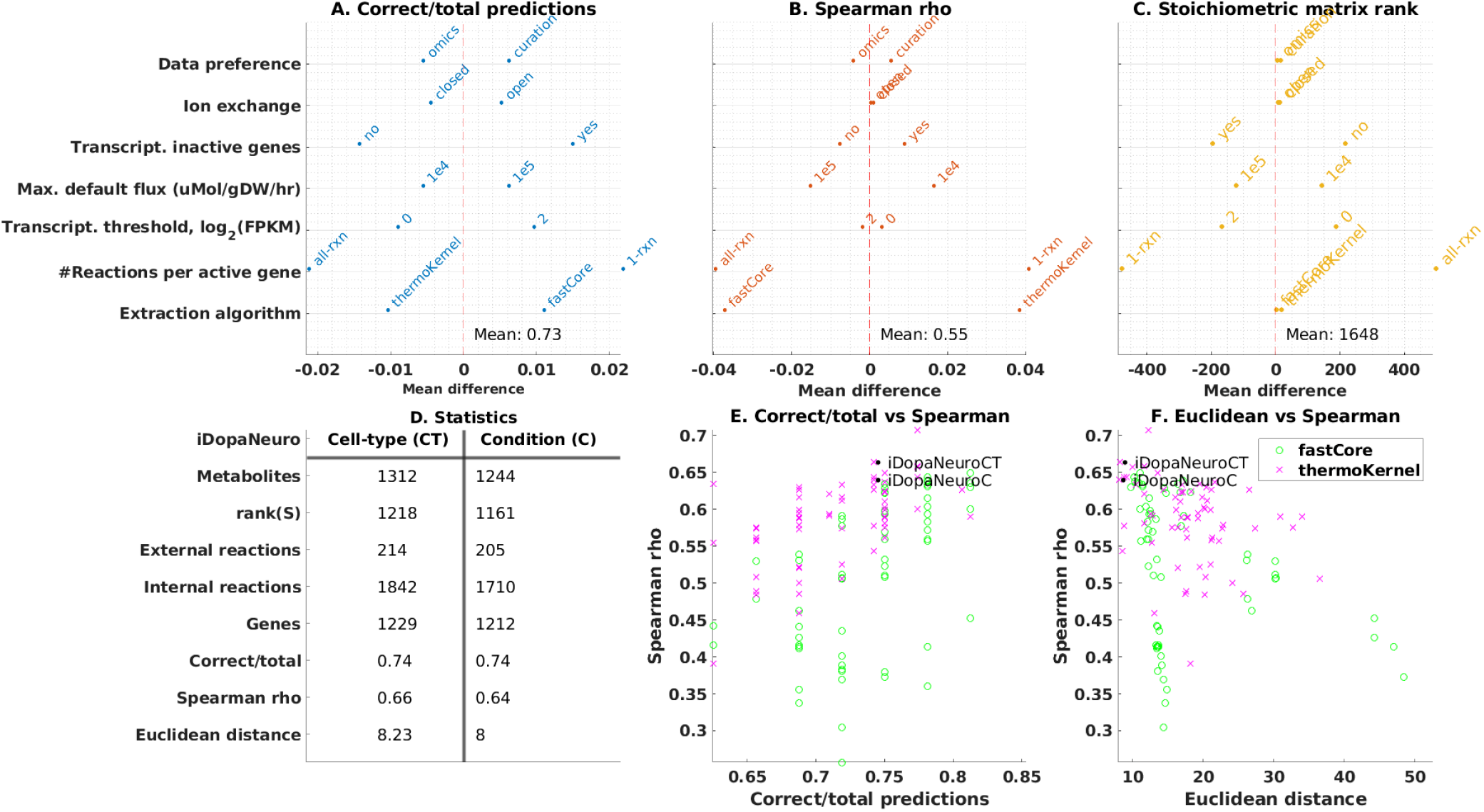
Exometabolomic data driven identification of iDopaNeuro models. (**A-C**) The effects on predictive accuracy and model size of changing XomicsToModel pipeline parameters with maximisation of flux entropy as the objective. Specifically, (**A**) Qualitative accuracy of predictions, given by the number of correct predictions of secretion/uptake divided by the total number of predictions. (**B**) Semi-quantitative accuracy of predictions, given by the Spearman rank correlation coefficient between the predicted and measured exchange fluxes. (**C**) Average model size, given by the rank of the stoichiometric matrix (*S* ≔ [*N, B*] in Fig. 3). D: Cell-type specific and condition-specific (iDopaNeuroCT and iDopaNeuroC, respectively). (**D**) Dopaminergic models were defined as those with the best compromise between qualitative (correct/total), semi-quantitative (Spearman rho) and quantitative (Euclidean distance) predictive accuracy. (**E, F**): Using maximisation of flux entropy as the objective, there were substantial differences in predictive accuracy, depending on the choice of XomicsToModel pipeline parameters, such as thermoKernel or fastCore as the model extraction algorithm.

Using maximisation of flux entropy as the objective, from the aforementioned model ensemble, the final dopaminergic neuronal models were identified as the two models, each corresponding to a pair of derived uptake-constrained and secretion-constrained models with the highest predictive accuracy, which prioritised either in vitro experimental data (condition-specific model, iDopaNeuroC) or manual curation of the literature on dopaminergic neuronal metabolism (cell-type specific model, iDopaNeuroCT). Fig. 9D provides basic statistics on the properties and predictive accuracy of these models. Both models used the same model generation parameters, including thermoKernel as the model extraction algorithm, at least one active reaction per active gene (1-rxn in Fig. 9) as the method to include reactions corresponding to active genes, two fragments per kilobase of transcript per million mapped reads as the transcriptomic threshold (on logarithmic scale), below which genes were designated as inactive, and ±1 × 10^5^*μmol*/*gDW* /*hr* as the default maximum and minimum flux bounds.

### 6.2 Dopaminergic neuronal metabolic model exploration

The condition-specific and cell-type specific dopaminergic neuronal models largely overlap in their content as they share 1,667 reactions, with 248 reactions exclusive to the condition-specific model and 389 reactions exclusive to the cell-type-specific model (Fig. **10**A). Using maximisation of internal flux entropy and quadratic penalisation of deviation from experimentally measured exchanges as the objective function, both models generate similar phenotypic predictions. Figure 10B-E illustrates the predicted rate of certain key metabolic reactions as a function of varying ATP demand. In both models, the most active metabolic pathways are related to energy production (glycolysis, oxidative phosphorylation) and both use glucose as the main carbon source (EX_glc_D[e]), while other important carbon sources include glutamine, lysine, and glycine (EX_gln_L[e], EX_lys_L[e] and EX_gly[e]).

**Figure 10:**
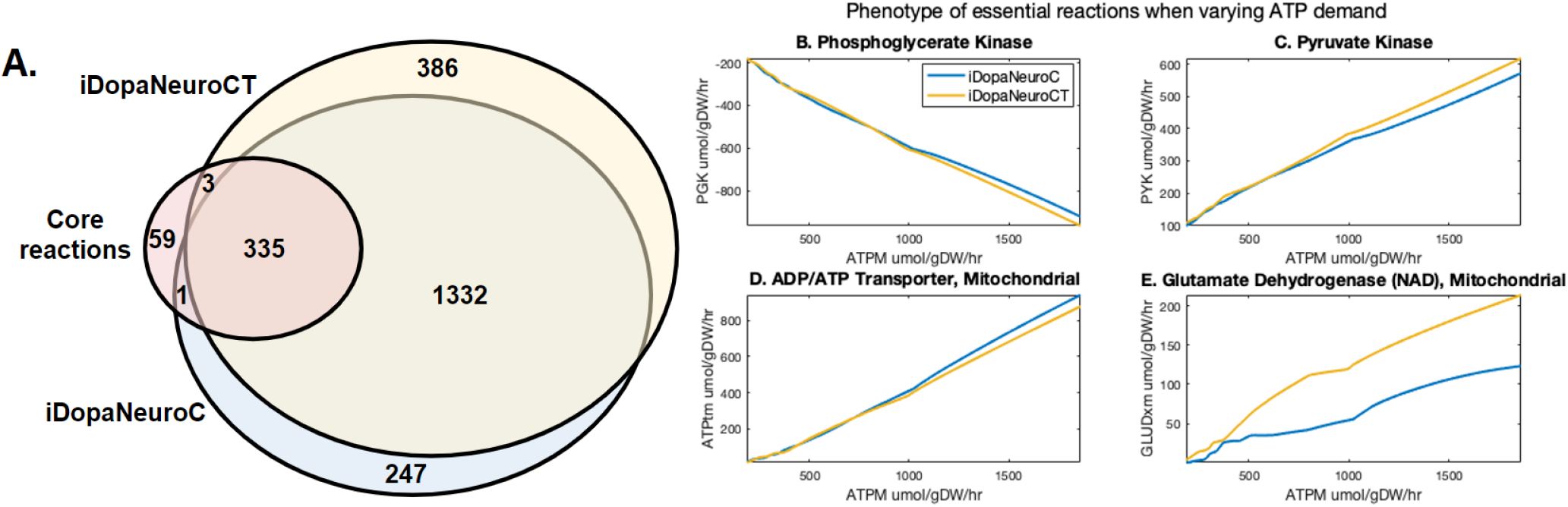
Comparison of condition-specific and cell-type-specific iDopaNeuro models. Comparison of the condition-specific model (iDopaNeuroC, blue) and cell-type-specific model (iDopaNeuroCT, yellow) with respect to: (**A**) reaction content, including overlap with core reactions (red); (**B**) predicted phosphoglycerate kinase (PGK) flux; (**C**) predicted pyruvate kinase (PYK) flux; (**D**) predicted ATP transport from the mitochondria (ATPtm) and (**E**) predicted mitochondrial glutamate dehydrogenase (GLUDXm) flux, as a function of varying demand for ATP (ATPM), using maximisation of internal flux entropy and quadratic penalisation of deviation from experimentally measured exchanges as the objective function.

**Figure 11:**
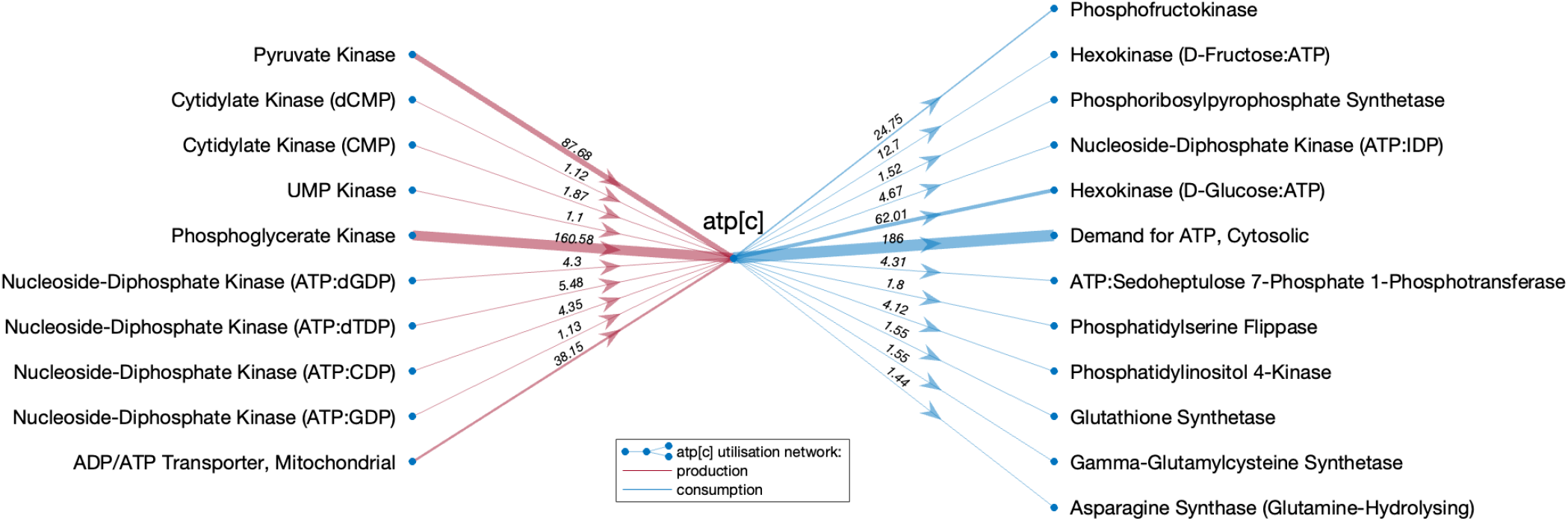
Cytosolic ATP utilisation network in the iDopaNeuro C model. Reactions producing (red) and consuming (blue) cytosolic ATP, above a threshold in flux magnitude (> 1 *μmol*/*gDW*/*hr*), with arrow thickness in proportion to magnitude. Fluxes were predicted using maximisation of internal flux entropy and quadratic penalisation of deviation from experimentally measured exchanges as the objective function. Glycolysis is the primary source of cytosolic ATP in the control model (pyruvate kinase and phosphoglycerate kinase) followed by the mitochondrial ATP production (ADP/ATP Transporter, Mitochondrial). The glycolytic phenotype is also reflected in the two key ATP consuming reactions - phosphofructokinase and hexokinase - which are also parts of glycolysis.

In total both models had 153 reactions involved in the metabolism of cytosolic ATP, out of which there were 16 and 17 reactions actively producing cytosolic ATP in iDopaNeuroC and iDopaNeuroCT respectively. Furthermore, 45 and 21 reactions were consuming cytosolic ATP in iDopaNeuroC and iDopaNeuroCT models respectively. The main sources of ATP in both models were oxidative phosphorylation (ATPtm) and glycolysis (PGK and PYK). Besides the demand for ATP (ATPM), which represents requirements for cellular maintenance and electrophysiological activity, the main consumers of cytosolic ATP are glycolytic reactions such as hexokinase (HEX1) and phosphofructokinase (PFK) (Figure 3.1.6).

Another key metabolite - mitochondrial NADH participates in a total of 51 and 62 reactions in iDopaNeuroC and iDopaNeuroCT models respectively. However, only 8 reactions in iDopaNeuroC and 10 reactions in iDopaNeuroCT are actively producing NADH, while 8 reactions in both models are consuming mitochondrial NADH. In both models, the two main producers of NADH are mitochondrial glutamate dehydrogenase and 2-oxyglutarate dehydrogenase, while mitochondrial complex I (NADH2_u10minew) and L-Glutamate 5-semialdehyde:NAD+ oxidoreductase and the key consumers of NADH in the iDopaNeuroC and iDopaNeuroCT models respectively (Figure 12).

**Figure 12:**
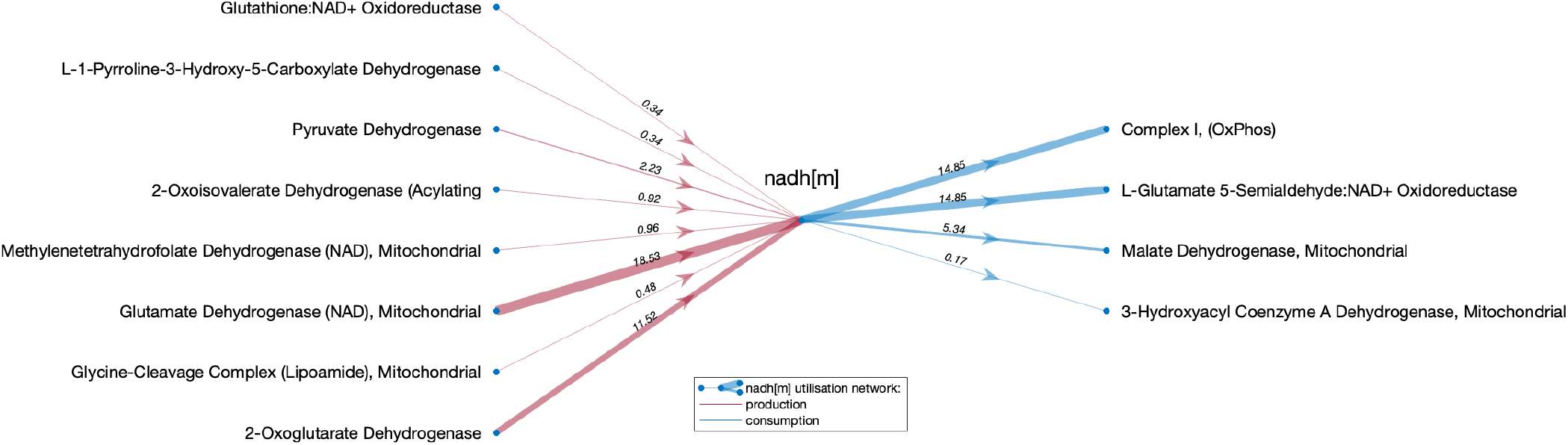
Mitochondrial NADH utilisation network in the iDopaNeuro C model. Reactions producing (red) and consuming (blue) mitochondrial NADH, above a threshold in flux magnitude (> 0.1*μmol*/*gDW*/*hr*), with arrow thickness in proportion to magnitude. Fluxes were predicted using maximisation of internal flux entropy and quadratic penalisation of deviation from experimentally measured exchanges as the objective function. Two key reactions generating mitochondrial NADH in the control model are glutamate dehydrogenase and 2-oxoglutarate dehydrogenase, responsible for the conversion of glutamate to alpha-ketoglutarate and a subsequent conversion to succinate-CoA, which are essential parts of the glutamate usage as a Krebbs cycle substrate. The mitochondrial NADH is then oxidised mostly via Complex I, providing electrons to the electron-transfer-chain and, eventually production of ATP. The second prominent reaction that oxidises NADH to NAD+ is L-Glutamate 5-semialdehyde:NAD+ Oxidoreductase which converts glutamate into the proline precursor - L-glutamate 5-semialdehyde.

Figure 13 illustrates that both models predict similar rates of exchange of unmeasured metabolites, when the models are fitted to measured exometabolomic exchange (Supp. Fig. 30) obtained from *in vitro* cultures of dopaminergic neurons in glucose maintenance medium. Both models were constrained to only permit uptake of metabolites present in the defined medium whereas the potential to secrete metabolites is either a consequence of manual literature curation, or a novel prediction. The condition-specific dopaminergic neuronal model contains 214 exchange reactions, of which 17 are for metabolite uptake, 137 are for metabolite secretion and 51 are open reversible exchange reactions, e.g., for the transport of water, (Table S-3). Out of the 188 metabolites with the potential to be secreted, most (112/188) were expected based on the assignment of corresponding transport reactions as active reactions during manual curation of the literature and exometabolomic measurements of spent media (Table S-1, S-2 and S-3), while the remainder (72/188) are novel predictions of metabolites with the potential to be secreted by dopaminergic neurons.

**Figure 13:**
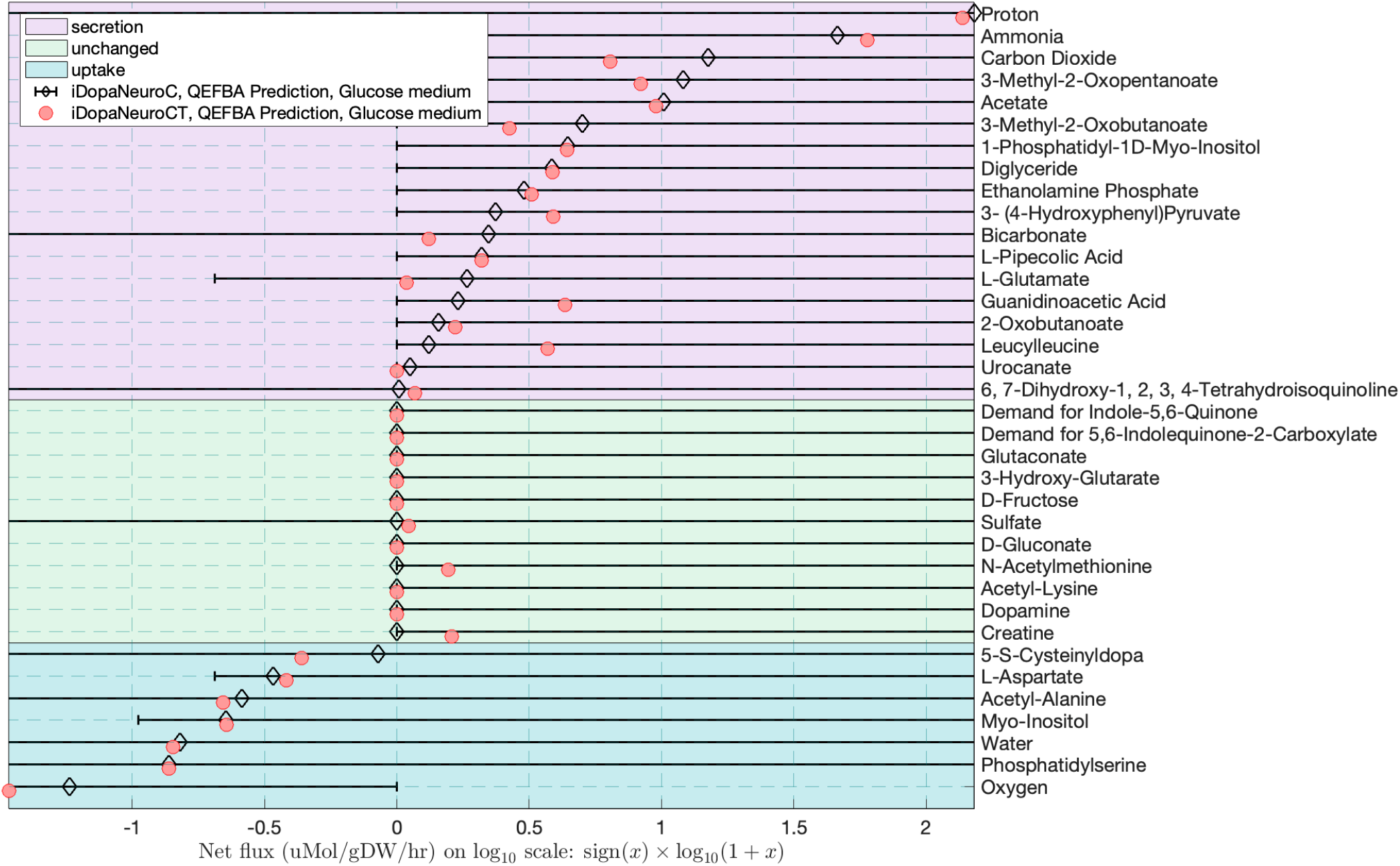
Prediction of unmeasured metabolite exchanges. The predicted rates of unmeasured exchange fluxes are very similar for the condition-specific (iDopaNeuroC) and cell-type specific (iDopaNeuroCT) models, when obtained by maximisation internal flux entropy and quadratic fitting to measured exchange fluxes (Supp Fig. 30) in glucose medium.

### 6.3 Model validation

The condition-specific dopaminergic neuronal model (iDopaNeuroC) was used to predict fluxes in response to several metabolic perturbations, namely complex I inhibition, complex V inhibition, and switching the carbon source from glucose to galactose. Leave-one-out cross validation was then used to compare these predictions with experimental data that was independent of the ensemble modelling process. Qualitative accuracy was evaluated for measured uptake and secretion, while semi-quantitative accuracy was evaluated compared to the mean of all measured exchange fluxes. Complex I inhibition predictions (Fig. 14) were qualitatively accurate (correct/total = 0.78, n = 9) and moderately semi-quantitatively accurate (*ρ* = 0.48, *pνal* = 0.018). Complex V inhibition predictions (Supplementary Fig. 22) were moderately qualitatively accurate (correct/total = 0.7, n = 13) and moderately semi-quantitatively accurate (*ρ* = 0.65, *pνal* = 0.0007). Predictions for switching from glucose to galactose (Supplementary Fig. 23) were moderately qualitatively accurate (correct/total = 0.68, n = 19) and moderately semi-quantitatively accurate (*ρ* = 0.55, *pνal* = 0.006). Supplementary Figure 26 illustrates the predicted consequences of complete inhibition of GBA1 on internal and external reaction fluxes.

**Figure 14:**
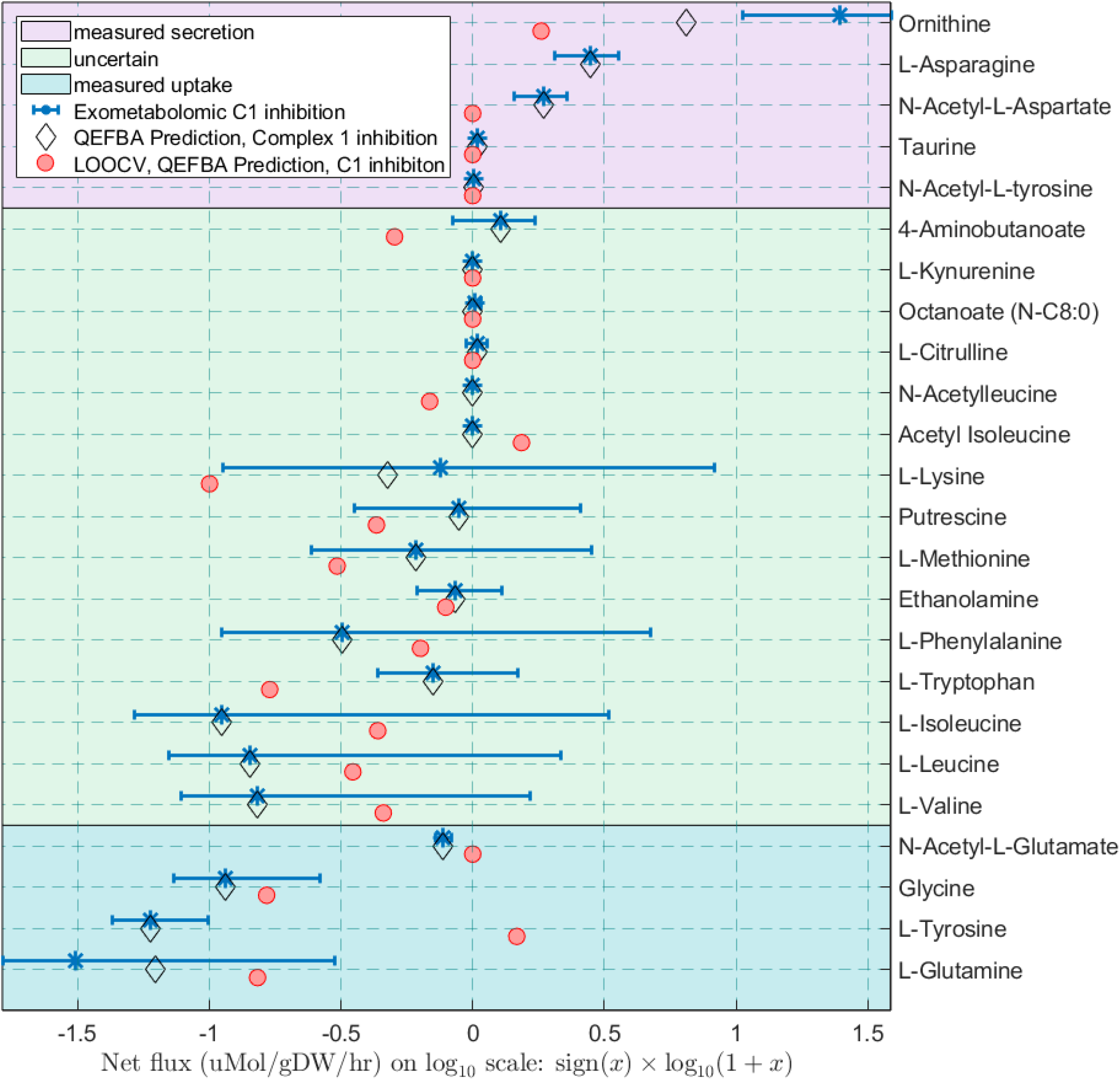
Comparison of measured and predicted exchange fluxes for complex I inhibition. Predictions were of high qualitative accuracy (correct/total = 0.78, n = 9) and moderately semi-quantitatively accurate (*ρ* = 0.48, *pνal* = 0.018). Quadratic penalisation of exchange flux deviation from measured exchanges with entropic flux balance analysis (QEFBA) is compared with the same approach except with omission of one experimentally measured exchange flux for each metabolite in the leave-one-out cross validation (LOOCV).

### 6.4 Biochemical interpretation of dopaminergic neuronal model perturbations

Following the analysis of the ATP and NADH utilisation in the iDopaNeuroC, which highlighted the model*’*s reliance on both glycolysis and mitochondrial ATP production to fulfill its metabolic needs, analysis of metabolic rearrangements in response to the various metabolic perturbations was performed. All perturbations (Complex I inhibition, Complex V inhibition, and a carbon source switch from glucose to galactose) caused an increase in flux though the glycolysis reactions in the iDopaNeuroC model (Fig. 15) with Complex V inhibition showing more than 20% increase when compared to the control model (Fig. 24. Similarly, all the perturbations resulted in the increase in pentose phosphate pathway with the pathway being only marginally active in the control model. However, the overall flux through the pentose phosphate pathway remained relatively low. However, all perturbations had a negative effect on mitochondrial energy metabolism, as seen through the changes to the reaction fluxes representing all the complexes of the mitochondrial electron transfer chain (Fig. 15. Complex V inhibition shows the most severe effect, with only Complex II (succinate dehydrogenase), Complex III (coenzyme Q : cytochrome c – oxidoreductase) and Complex IV (cytochrome c oxidase) still carrying a minimal flux. Interestingly, in Complex I inhibition and in the case of a carbon change from glucose to galactose, the flux through the Complex II was predicted to be slightly increased.

**Figure 15:**
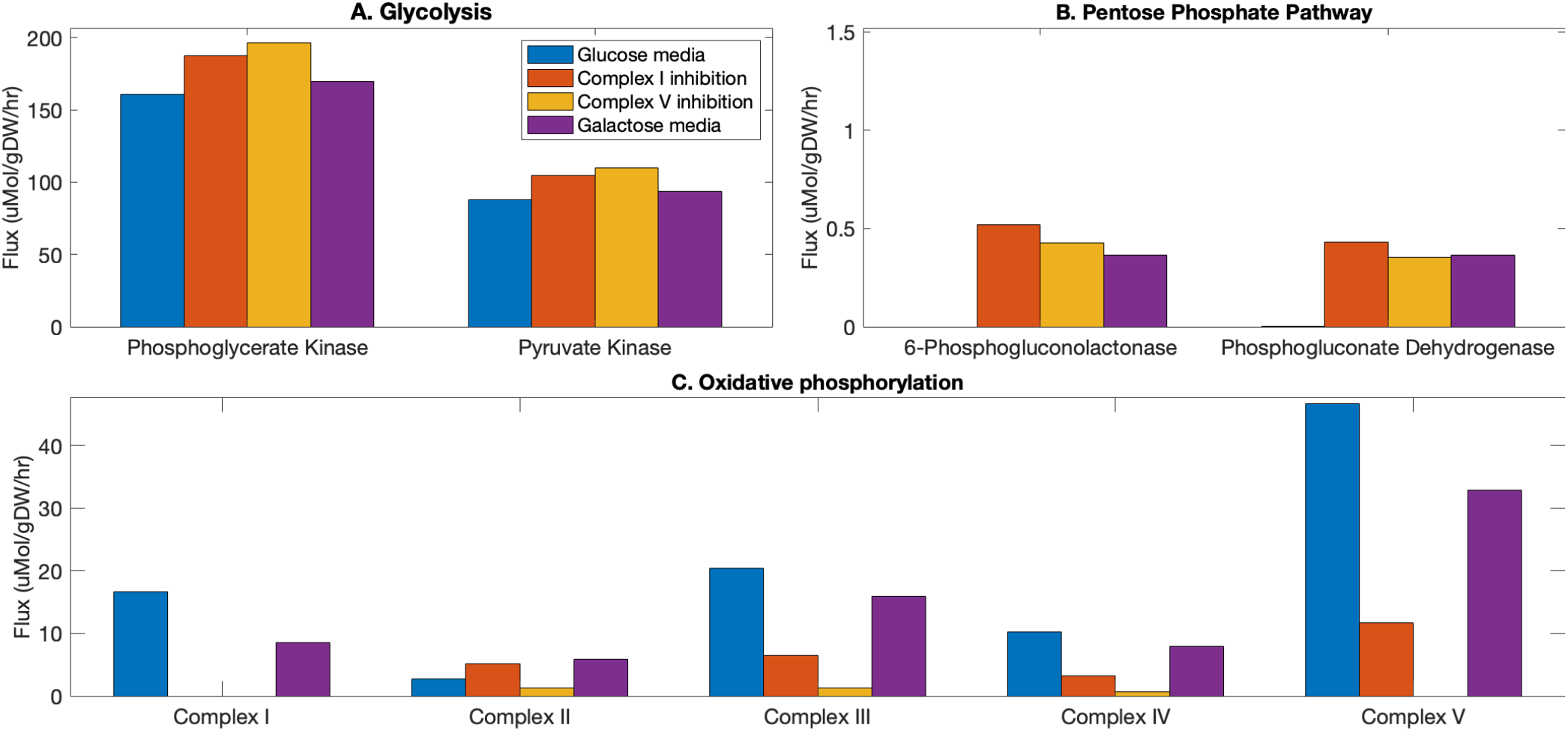
Predictions of the core energy-related reaction fluxes due to the metabolic perturbations. Predicted fluxes (*μmol*/*gDW*/*hr*) in control iDopaNeuroC model (blue) and in response to complex I inhibition (red), complex V inhibition (yellow), and change from glucose to galactose as carbon source (purple), obtained by internal flux entropy maximisation and quadratic penalisation of deviation from measured exchange fluxes. Reactions have been grouped by the metabolic pathway are part of (glycolysis, pentose phosphate pathway, and oxidative phosphorylation). Fluxes were predicted using maximisation of internal flux entropy and quadratic penalisation of deviation from experimentally measured exchanges as the objective function.

The net cytosolic ATP budget was further analysed to reveal key reactions that produce and consume cytosolic ATP after Complex I inhibition. Cytosolic ATP was predicted to be produced predominantly via two glycolysis reactions catalysed by pyruvate kinase (PYK, increase of 19% compared to control) and phosphoglycerate kinase (PGK, increase of 17%) and only marginally (reduction by 70%) via TCA and oxidative phosphorylation (ADP/ATP Transporter, Mitochondrial, Fig. 16). Following Complex I inhibition the biggest changes are seen in mitochondrial NADH utilisation (Fig. 17). Two NADH producing reactions have become inactive in response to the Complex I inhibition, such as pyruvate dehydrogenase glycine-cleavage complex, and 3-hydroxyacyl coenzyme A dehydrogenase. Furthermore, two reactions - glutathione:NAD+ oxidoreductase and methylenetetrahydrofolate dehydrogenase - are predicted to change their directionality from NADH producers to NADH consumers in response to teh Complex I inhibition. Among the major NADH producers, glutamate dehydrogenase (NAD) flux was significantly reduced (73% compared to the control), but 2-oxaloglutarate dehydrogenase flux was increased. Furthermore, several reactions that oxidise NADH back to NAD+ have become active, including bile acid-related reactions (e.g. CYP27A1), or have an increased flux (glycine synthase), with an exception of malate dehydrogenase reaction which has decreased flux (69% compared to the control) in response to the Complex I inhibition, and 3-hydroxyacyl coenzyme A dehydrogenase reaction, which became inactive.

**Figure 16:**
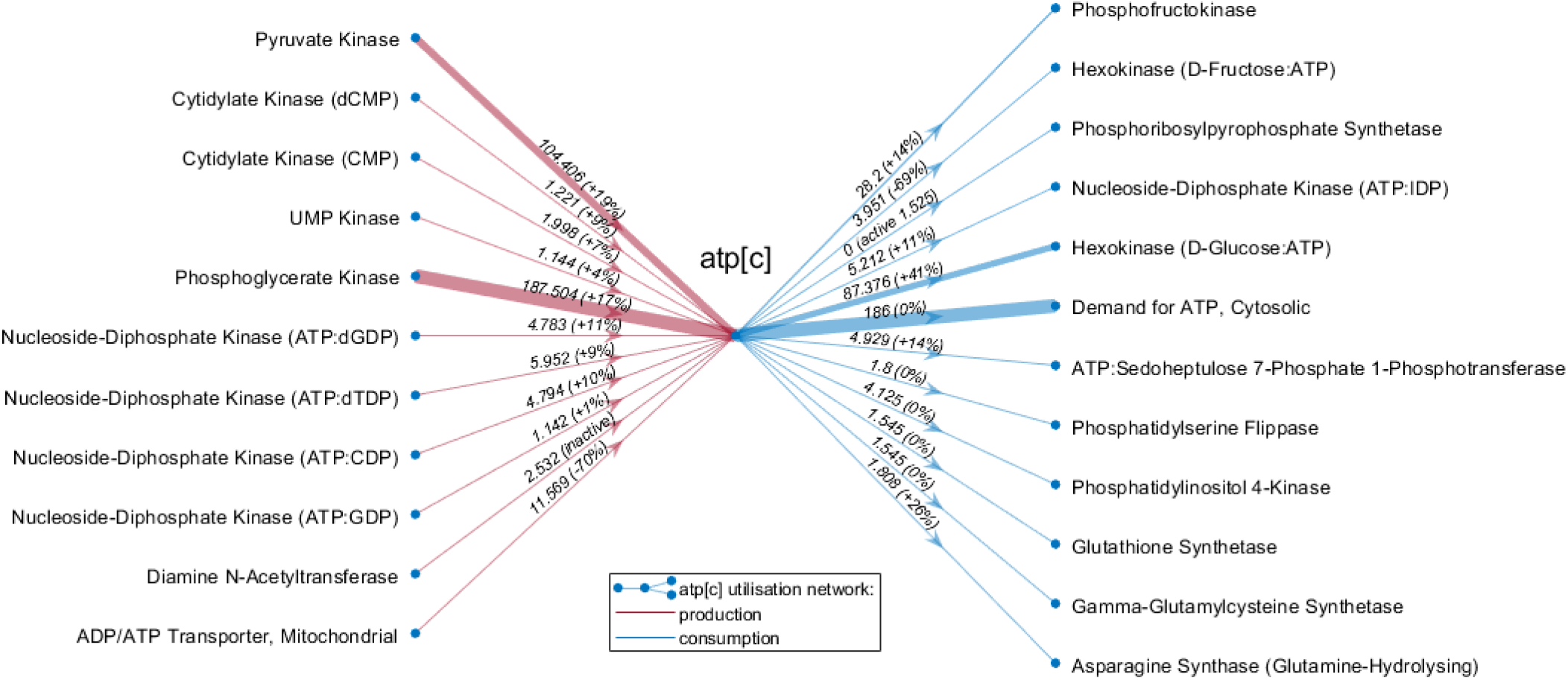
Cytoplasmic ATP utilisation after Complex I inhibition. Reactions producing (red) and consuming (blue) cytoplasmic ATP, above a threshold in flux magnitude (> 1 *μmol*/*gDW*/*hr*), with arrow thickness in proportion to magnitude of predicted flux in Complex I inhibition in the iDopaNeuroC model. Arrow*’*s labels show the flux value in Complex I inhibition and the flux % change compared to the control model. Fluxes were predicted using maximisation of internal flux entropy and quadratic penalisation of deviation from experimentally measured exchanges as the objective function. The major ATP producting reactions, pyruvate kinase and phosphoglycerate kinase, have both increased in flux in the Complex I inhibition prediction compared to the control model, while the net mitochondrial ATP production is strongly reduced (ADP/ATP Transporter, Mitochondria) afte Complex I inhibition. Similarly, glycolytic ATP-dependent reactions, phosphofructokinase and hexokinase, also are predicted to have an increased flux.

**Figure 17:**
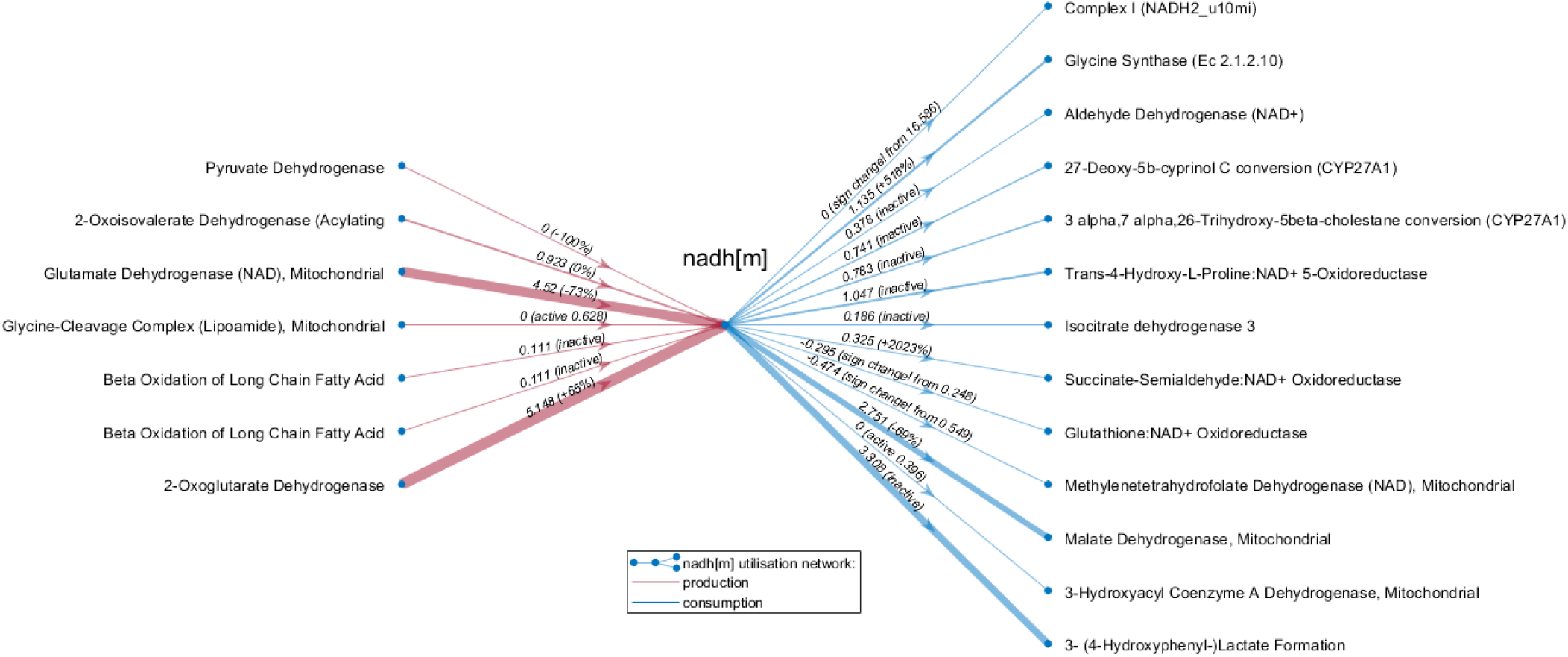
Mitochondrial NADH utilisation after Complex I inhibition. Reactions producing (red) and consuming (blue) mitochondrial NADH, above a threshold in flux magnitude (> 0.1*μmol*/*gDW*/*hr*), with arrow thickness in proportion to magnitude of predicted flux in Complex I inhibition. Arrow*’*s labels show the flux value in Complex I inhibition and the flux % change compared to the control model. Fluxes were predicted using maximisation of internal flux entropy and quadratic penalisation of deviation from experimentally measured exchanges as the objective function. All reactions significantly contributing to the production of mitochondrial NADH are predicted to be significantly affected by the Complex I inhibition. While pyruvate dehydrogenase and glutamate dehydrogenase (NAD) have both significantly reduced flux, 2-oxaloglutarate dehydrogenase*’*s flux is increased. Furthermore, two reactions - glutathione:NAD+ oxidoreductase and methylenetetrahydrofolate dehydrogenase - are predicted to change their directionality from NADH producers to NADH consumers in response to teh Complex I inhibition. On the NADH consumption side, several reactions that oxidise NADH back to NAD+ have become active, including bile acid-related reactions of CYP27A1, or have an increased flux (glycine synthase), with an exception of malate dehydrogenase reaction which has decreased flux (69% compared tot he control) in response to the Complex I inhibition, and 3-hydroxyacyl coenzyme A dehydrogenase reaction, which became inactive.

## 7 Prospective research design

Given the iDopaNeuroC model, a novel *uncertainty reduction* pipeline (Fig. 31A-C) predicted the metabolites whose corresponding external reactions would be the most important to constrain in future research, in order to maximally shrink the set of external reaction fluxes consistent with existing constraints (Ω ≔ {*ν*|*Sν* = 0, *l* ≤ *ν* ≤ *u*} in 3). Fig. 31D illustrates the top 20 most informative metabolites in exchange reactions to constrain to decrease the uncertainty in the iDopaNeuroC model. 5-S-Cysteinyldopamine and 5-S-Cysteinyldopa are formed by dopaquinone and dopamine-o-quinone, respectively, binding to free cysteine residues on proteins [80]. The next most important variables to constrain include water, acetate, protons bicarbonate and carbon dioxide, reflecting their general metabolic importance. Other metabolites point to opportunities for further development of quantitative metabolomic platforms, either the inclusion as new targets for quantification, e.g., (-)-Salsolinol, hypothesised to be formed by reaction of dopamine with acetaldehyde [81]. Supplementary Figure 31 illustrates the same result for the iDopaNeuroCT and Table S-12 contains the full list of prioritised metabolites for both models.

## Discussion

### Contribution overview

Degeneration and death of substantia nigra dopaminergic neurons in the midbrain is a hallmark of PD. Herein, we present, characterise and utilise comprehensive, high-quality, thermodynamically constrained, *in vitro* condition-specific (iDopaNeuroC) and cell-type-specific (iDopaNeuroCT), models of normal human dopaminergic neuronal metabolism. Their comprehensiveness reflects a synthesis of extensive manual curation of the biochemical literature on neuronal metabolism, together with novel, quantitative, transcriptomic and targeted exometabolomic data from human midbrain-specific dopaminergic neurons *in vitro*. The predictive fidelity of the models was evaluated by comparing their ability to quantitatively predict the rate of metabolite exchange in metabolically perturbed conditions. Leave-one-out cross validation of a condition-specific dopaminergic neuronal metabolic model of complex I inhibition was qualitative accurate (correct/total = 0.78, n = 9) and moderately semi-quantitatively accurate (*ρ* = 0.49, *pνal* = 0.018) at predicting secretion or uptake when compared with exometabolomic data (Fig. 14). Similar accuracy was achieved for complex V inhibition and for replacement of glucose with galactose as the main carbon source. The development and characterisation of the iDopaNeuro models also illustrates the application of a novel, scalable approach to thermodynamic constraint-based modelling. All reactions admit thermodynamically feasible fluxes and the practical utility of thermodynamic variational principles [82] are demonstrated at genome-scale for the first time.

### Relationship between *in vivo, in vitro*, and *in silico*

Manual curation of the literature was focused on the assembly of neuronal molecular composition, turnover fluxes, active genes, active reactions and inactive reactions specific to neurons, and substantia nigra dopaminergic neurons in particular (Tables S-3, S-4 and S-5). In parallel, we integrated transcriptomic and metabolomic data from an *in vitro* culture of midbrain dopaminergic neurons. As such, the iDopaNeuro models are *in silico* models that represent both *in vivo* substantia nigra dopaminergic neurons and *in vitro* midbrain-specific dopaminergic neurons, derived from human neuroepithelial stem cells [18]. To control for prioritisation of *in vivo* or *in vitro* data, we generated condition-specific (iDopaNeuroC)and cell-type-specific (iDopaNeuroCT) models, that prioritised *in vitro* and *in vivo* data, respectively. The protocol for generation of human midbrain-specific dopaminergic neurons *in vitro* is well established [18]. We observed that it generates tonically firing neurons that do have extensive neuronal projections. However, in rats, a *single* dopaminergic neuron emanating from the substantia nigra is characterised by a massive axonal arbour [83], much larger than other neuronal types, and projects to ∼200k terminals in the striatum [8].

Like this morphological divergence between *in vivo* and *in vitro*, there may be a molecular divergence between *in vivo* neurons, on which manual literature curation was based, and the *in vitro* neuronal culture used for generation of transcriptomic and metabolomic data. As such, it will be interesting to compare these versions of the iDopaNeuro models with future versions generated using data from improved protocols for generation of dopaminergic neuronal cultures.

### Metabolomics

The quality of the metabolomic analyses was assessed by comparing the measured and supplier reported concentration values (Fig. 7). Based on this comparison, for most of the measured metabolites, the measurements were obtained within a similar concentration value (±20%) of that specified by the manufacturer of the fresh culture media. However, some measurements (*e*.*g*. choline, ethanolamine, cysteine, histidine, threonine and putrescine) demonstrated larger differences between measured and concentrations specified by the manufacturer. There could be several explanations for the discrepancies observed. Some compounds may undergo spontaneous reactions with other metabolites in their environment due to their reductive, or oxidative nature, or both. For example, the thiol group of cysteine is reactive with oxidants and reductants and has high affinity for metals [84]. Therefore, it can be very difficult to determine the quantity of free cysteine residues after protein hydrolysis unless the thiol group of the molecule is stabilised by chemical derivatisation procedures [85]. This might explain the observation of a lower value for the measured concentration in comparison to the manufacturer specified value 7. The discrepancy could also be due to the incomplete knowledge on the composition and concentration of medium supplements, e.g., B21 medium supplement contains a confidential amount of putrescine which may have contributed to the large increase in the measured concentrations.

### Reconstruction versus model

The starting point for development of the dopaminergic neuronal model was a generic human metabolic reconstruction, Recon2 [50], and subsequently its successor Recon3D [35], which included new reactions specific to dopaminergic neuronal metabolism. It is critical to distinguish between a reconstruction and a model. The former is broader in coverage of metabolic pathways and may contain partially specified reaction stoichiometry, e.g., due to missing biochemical knowledge. A model is always a subset of a reconstruction as it must satisfy certain modelling assumptions. For example, a model is suitable for flux balance analysis [86] when all internal reactions are stoichiometric and flux consistent, and all external reactions are flux consistent [87]. However, when extracting a model, certain otherwise desirable reactions may be lost. As an example, the glutathione transferase reaction in dopamine metabolism was manually curated to be active since it is present in dopaminergic neurons [75], but it was excluded during model generation as it was not part of any thermodynamically flux consistent pathway in Recon3D model. This example illustrates that future manual curation of the literature is required in the next iteration of the generic human metabolic reconstruction.

### Model generation

The generation of the iDopaNeuro models required several advances in constraint-based reconstruction and analysis. Herein we present the first application of a novel model generation pipeline, XomicsToModel [36], that enables context-specific model generation by leveraging high-fidelity data obtained by literature curation and integration of high-throughput data obtained from multiple omics platforms (Fig. 1). In particular, we present the first application of thermoKernel, a novel model extraction algorithm that accepts weighted inputs on molecular abundance, e.g., confidence in presence of a metabolite, as well as weighted inputs on reaction fluxes, e.g., expression level of the corresponding gene, then simultaneously optimise these, perhaps conflicting, inputs to generate a minimal subnetwork that is thermodynamically consistent, i.e., each reaction admits a thermodynamically feasible net flux.

### Modelling non-growing cells

Constraint-based modelling is most commonly applied to biochemical systems where one predicts a steady-state flux vector that also satisfies a biologically-motivated cellular objective, e.g., for an exponentially growing culture of bacteria maximisation of biomass production flux subject to (external) uptake reaction constraints [86]. However, neither substantia nigra dopaminergic neurons nor *in vitro* differentiated dopaminergic neurons divide, and it is not known what the cellular objective is for such cell types. Therefore, we added new constraints that enforce certain internal reactions, or combinations thereof, to operate above a certain flux, e.g., lower bounds on metabolite turnover fluxes and constraints representing the energetic requirements for maintenance of biomass and tonic electrophysiological activity. Formulating a mathematical model that facilitates such constraints yet also admits a thermodynamically feasible net flux is an important prerequisite for modelling non-growing cells, such as neurons. This is because it enables the subsequent application of novel thermodynamic constraint-based modelling approaches that enforce thermodynamic principles, such as energy conservation, the second law of thermodynamics [88], and non-equilibrium thermodynamic force-flow relationships [89]. Reformulating the corresponding non-linear and/or non-convex constraints as convex optimisation problems [82, 90] enabled us to implement them in a mathematically and computationally tractable manner. The addition of such constraints also act to compensate for an absence of evolutionarily motivated objectives, such as maximisation of the rate of one or more external reactions.

### Ensemble modelling

It was not possible to use standard constraint-based modelling methods, such as flux balance analysis [86] to accurately predict metabolite exchanges. In part, this is because the cellular objective of a dopaminergic neuron is not known, so it is not clear what to optimise for. Leveraging the flexibility of the XomicsToModel pipeline, we generated an ensemble of candidate dopaminergic neuronal metabolic models, varying model generation parameters, including those known to substantially change the computational phenotype of an extracted model [62]. A set of candidate objective functions was applied to predict steady state fluxes with each of these candidate models. The most predictive objective was defined as the objective best able to predict uptake given secretion rates and secretion given uptake rates, on average over all models. Condition-specific and cell-type specific dopaminergic models were then defined models with the highest predictive capacity, using the most predictive objective, prioritising either condition-specific or cell type specific input data, respectively.

### Candidate cellular objectives

Candidate objectives tested included a variety of those already investigated in the literature, such as minimising two-norm flux [91] or one-norm flux [92, 93], and weighted versions of each, e.g., weighting one or two norm flux by the logarithm of the level of expression of the corresponding gene. Although other cellular objectives have been proposed in the literature [94, 95], we restricted our choice to those functions where a stationary point implies, and is implied by, attainment of a global minimum [96]. We consider that having mathematical guarantees, even if they are probabilistic, are vital when drawing conclusions from predictions with genome-scale models. Of the objectives tested, simultaneous maximisation of the entropy of unidirectional fluxes provided the most accurate prediction of metabolite exchanges. From an information theoretic perspective, entropy maximisation is the least biased prediction one can make given the available data [97]. This realisation came long after after entropy maximisation became established as a central tenet of equilibrium thermodynamics, including of systems of chemical reactions [98]. Maximising of flux entropy, subject to bounds on the direction of a subset of reactions, extends a theoretical consideration of flux entropy maximisation [82] and is also consistent with parallel developments in biophysics [99].

### Metabolic perturbations

Compared to other cells, neurons have low basal mitochondrial membrane potential, as well as low glycolytic activity, thus they rely on mitochondrial oxidative phosphorylation for their energy production [100]. Compared to other cells, neurons have been hypothesised to experience a higher degree of mitochondrial stress, and therefore, especially rely on a functional mechanism of mitochondrial quality for their survival [101]. In a well known PD familial mutation PINK1 one of the mitochondrial quality control systems is impaired and leads to a strong phenotype showing higher fluxes of glycolysis due to lower activity of the electron transport chain. This in consequence leads to a higher reliability on glycolysis in order to maintain cell viability, and a substantial increase in lactate release [102]. The loss of PINK1 function has been linked to NADH dehydrogenase and complex I dysfunction [103]. Complex I has also been linked to the development of idiopathic PD in people who experienced prolonged exposure to toxicants, e.g., rotenone and MPTP [104, 105]. The iDopaNeuroC model predictions suggest that complex I inhibition causes neurons to further increase their glycolytic flux by 19%, as seen in the ATP utilisation (Fig. 16) as well as in the predictions of increased glucose uptake and lactate secretion (Fig. 18) which is in line with PINK1 mutant observations [102]. Furthermore, the mitochondrial ATP production was severely impaired (−70%) in line with previous studies [102, 106]. Interestingly, the model predicted increased flux through the pentose phosphate pathway (6-P-gluconolactonase and phosphogluconate dehydrogenase reactions) as a consequence of complex I inhibition (Fig. 15). Since the pentose phosphate pathway plays an important role in protecting the cell from oxidative stress by reducing reactive oxygen species levels in the cell, an increased rate of the pentose phosphate pathway suggests higher levels of cellular stress in complex I inhibition. Similar response is see also in predictions of Complex V inhibition as well as a switch to galactose media.

**Figure 18:**
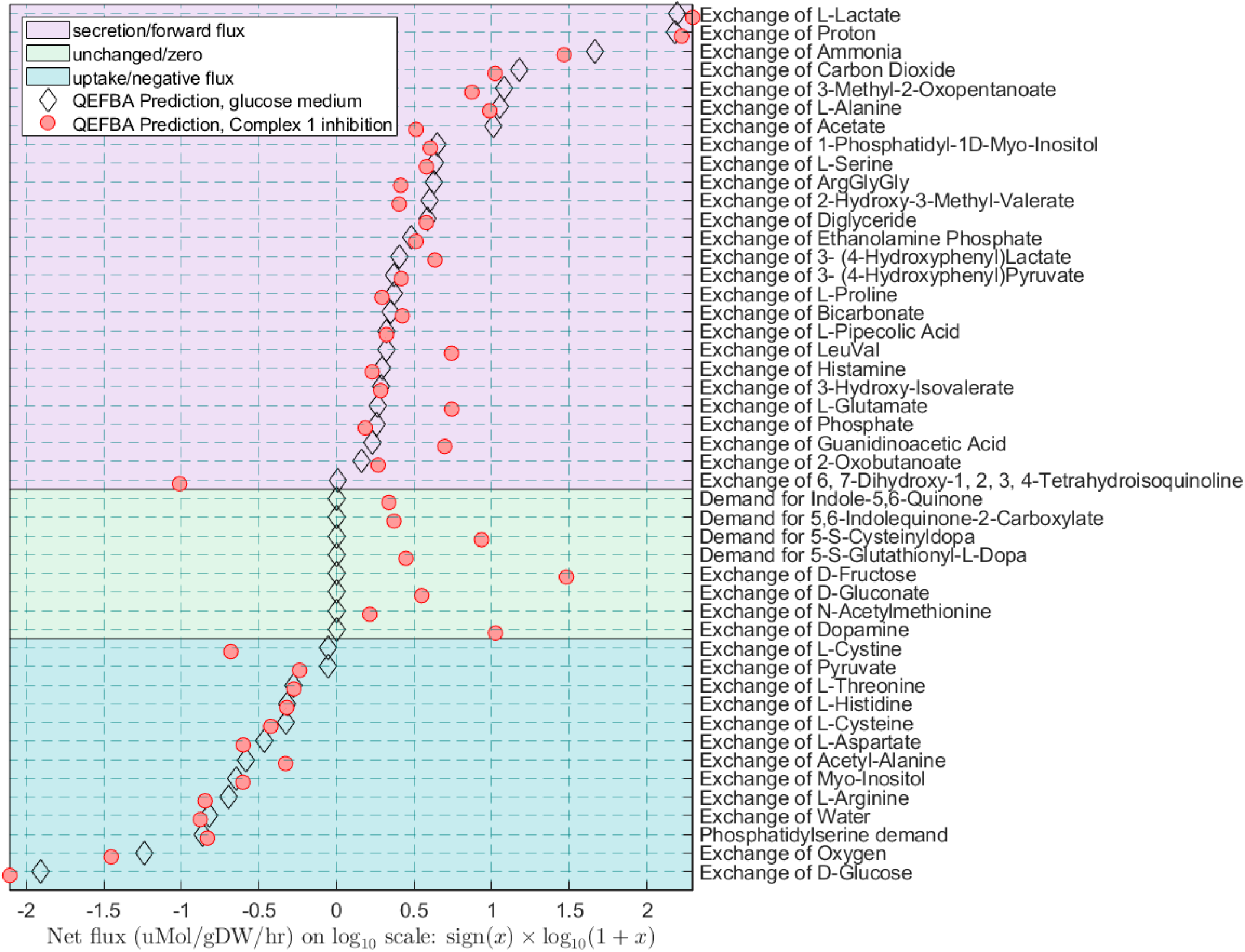
Predicted exchange reaction rates for unmeasured metabolites. Comparison between the predicted uptakes and secretion rates for exchange metabolites not measured experimentally in iDopaN-euroC model between control (black diamonds) and Complex I inhibition (red dots). Quadratic penalisation of exchange flux deviation from measured exchanges with entropic flux balance analysis (QEFBA) is compared with the same approach except with omission of one experimentally measured exchange flux for each metabolite in the leave-one-out cross validation (LOOCV).

**Figure 19:**
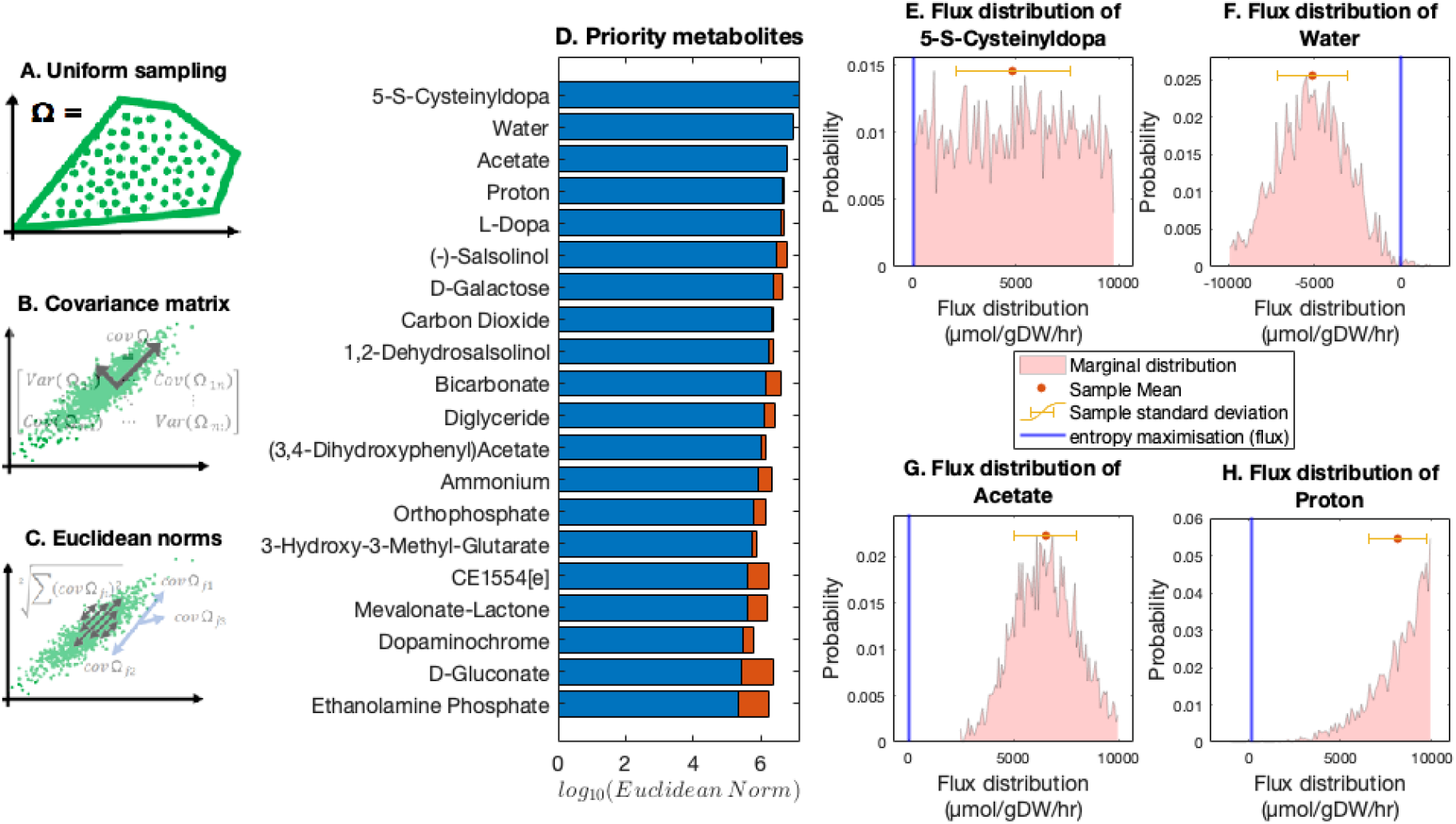
Prospective prioritisation of model variables to constrain. Uniform sampling of the steady-state flux space (Ω ≔ {*ν* | *Sν* = 0; *l* ≤ *ν* ≤ *u*} in 3), of the iDopaNeuroC model. (**B**) Computation of the covariance matrix of sampled external fluxes. (**C**) The Euclidean norm of each row of the covariance matrix identifies the exchange reaction with the highest degree of freedom. (**D**) The predicted most informative metabolites to measure, each corresponding to one external reaction flux, the iDopaNeuroC model. The variance reduction (to blue) due to cumulative constraints on higher-ranked metabolites (red) is taken into account in the ranking. (**E-H**) The flux distribution of the four most important exchanged metabolites to constrain in order to maximally shrink the steady-state flux space Ω, compared with the exchange fluxes predicted using the maximisation of the entropy of forward and reverse fluxes (blue, *ψ*_*ν*_ in 1).

**Figure 20:**
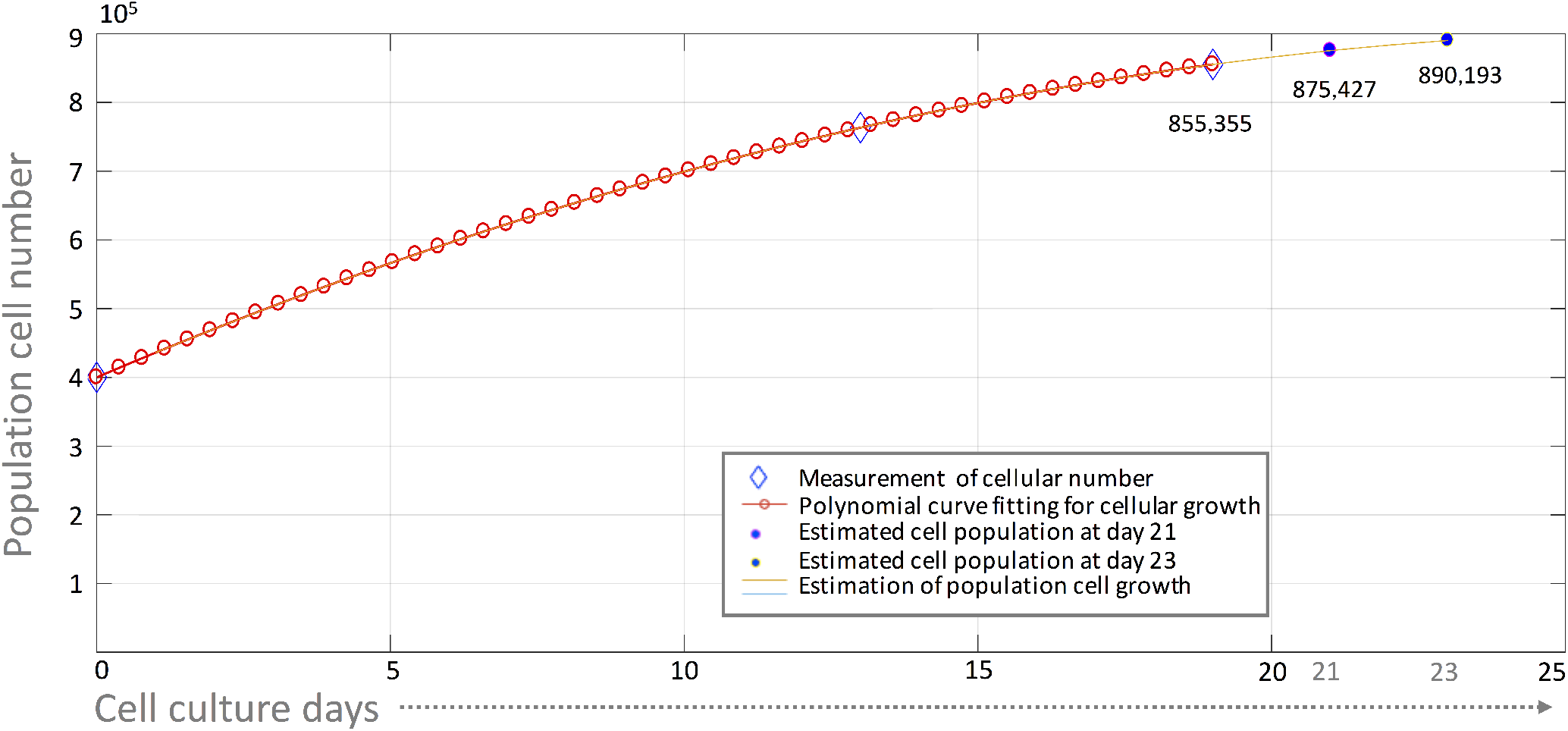
Measured and estimated cell numbers during neuronal differentiation. The cell number in each culture well was measured at day 0 (seeding, 400k cells per well.) and days 13, 19, but not day 21 or 23 in culture. The cell number at day 21 and 23 was estimated by interpolation in order to enable normalisation of metabolic uptake and secretion rates. Therefore, the evolution of cell number with respect to time was estimated using a cubic spline fit to the measured cell numbers. Exometabolomic data was collected at day 9, 13, 19 and 23. However, only exometabolomic data from day 19 and 23 were used to quantitatively constrain the models. This is consistent with the established differentiation protocol used, where a 30-45% increase in cell number is observed during the first five days and therefore an assumption that there was no biomass growth was not considered valid during the early period in cell culture.

**Figure 21:**
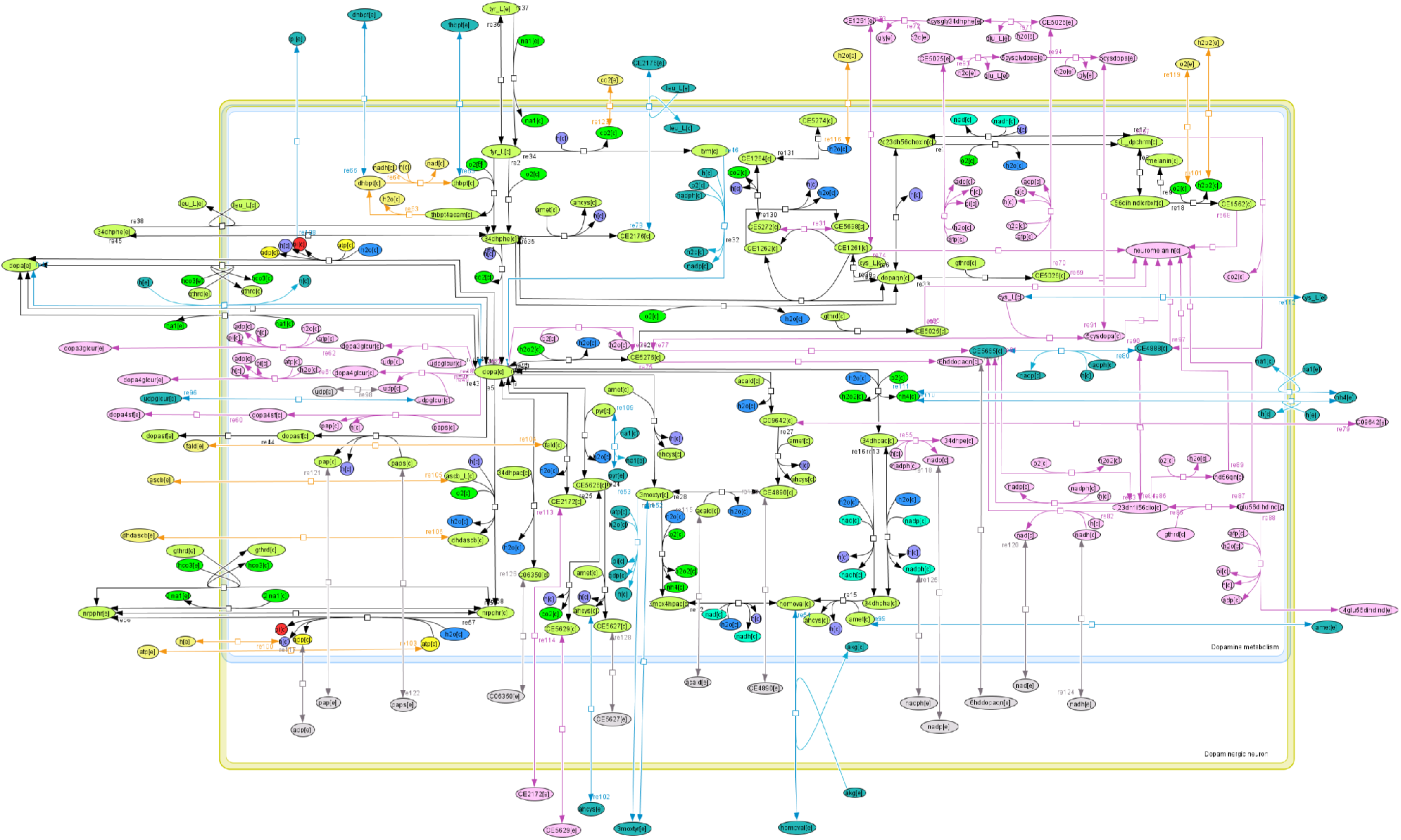
Reconstruction of dopamine metabolism. Dopamine metabolism in Recon2.04 (green, blue) was refined and updated with newly added reactions (pink).

**Figure 22:**
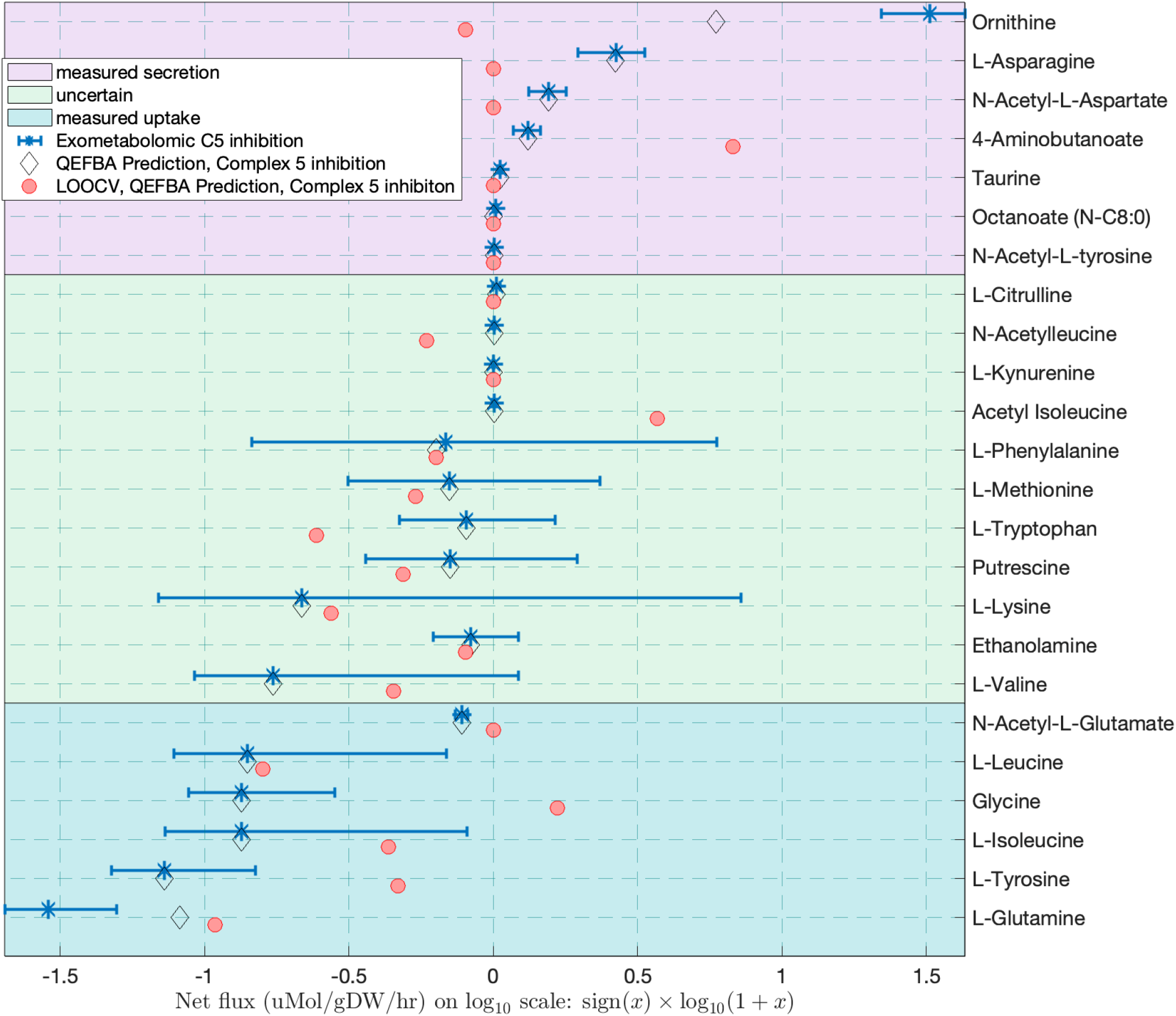
Comparison of measured and predicted exchange fluxes for complex V inhibition. When fresh versus spent media concentrations were compared, metabolites are grouped into those that were measured to be secreted (positive average minus one standard deviation, pink background), measured to be taken up (negative average plus one standard deviation, blue background) and measured with uncertain exchange (error bar includes zero, green background). Predictions were of high qualitative accuracy (correct/total = 0.7, n = 13) and moderately semi-quantitatively accurate (*ρ* = 0.65, *PνAL* = 0.0007). Quadratic penalisation of exchange flux deviation from measured exchanges with entropic flux balance analysis (QEFBA) is compared with the same approach except with omission of one experimentally measured exchange flux for each metabolite in the leave-one-out cross validation (LOOCV).

**Figure 23:**
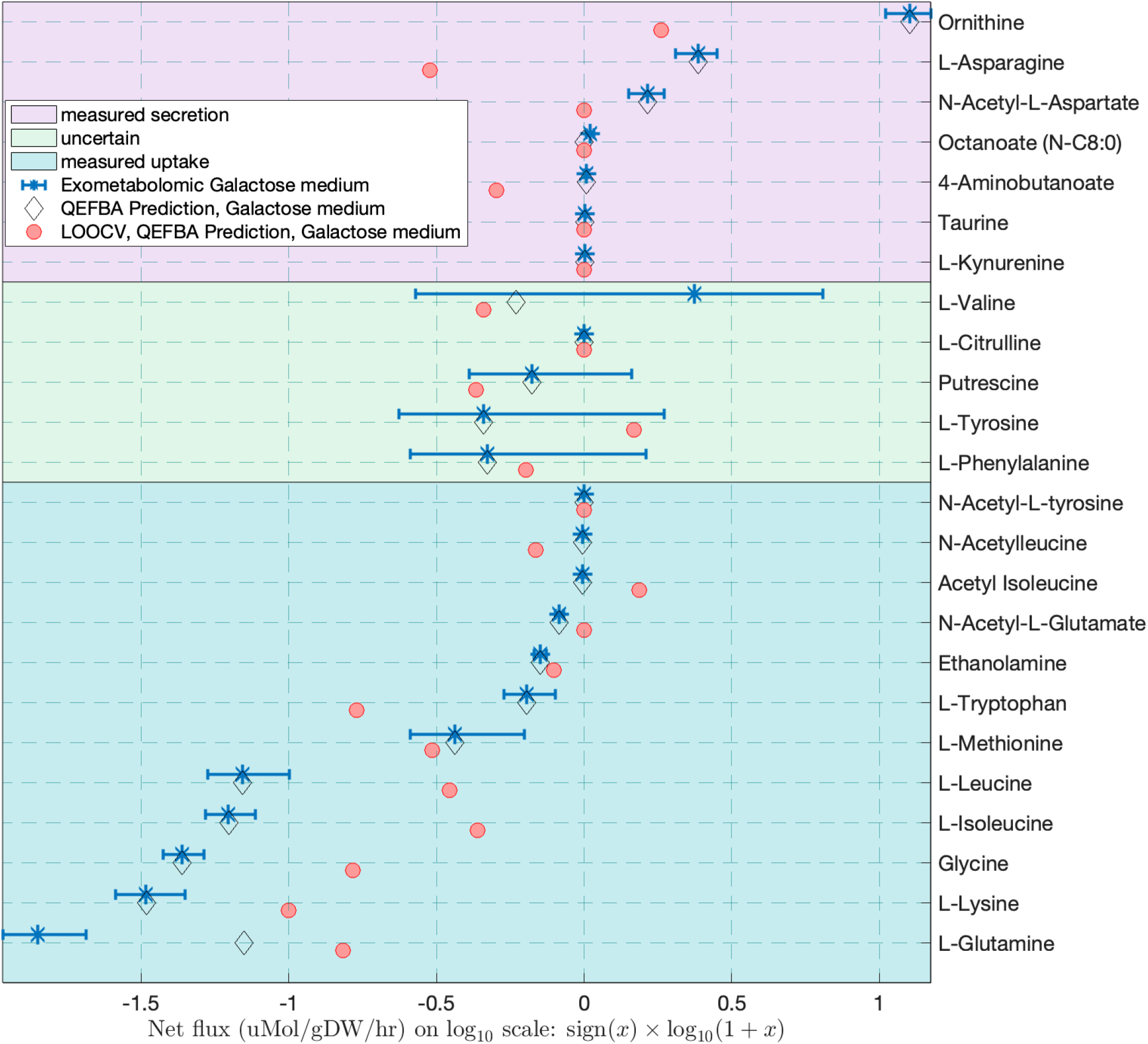
Comparison of measured and predicted exchange fluxes for switching from glucose to galactose. When comparing fresh versus spent media concentrations, metabolites are grouped into those that were measured to be secreted (positive average minus one standard deviation, pink background), measured to be taken up (negative average plus one standard deviation, blue background) and uncertain exchange direction (error bar includes zero, green background). Quadratic penalisation of exchange flux deviation from measured exchanges with entropic flux balance analysis (QEFBA) is compared with the same approach except with omission of one experimentally measured exchange flux for each metabolite in the leave-one-out cross validation (LOOCV). With leave-one-out cross validation, predictions were moderately qualitatively accurate (correct/total = 0.68, n = 19) and moderately semi-quantitatively accurate (*ρ* = 0.55, *PνAL* = 0.006).

**Figure 24:**
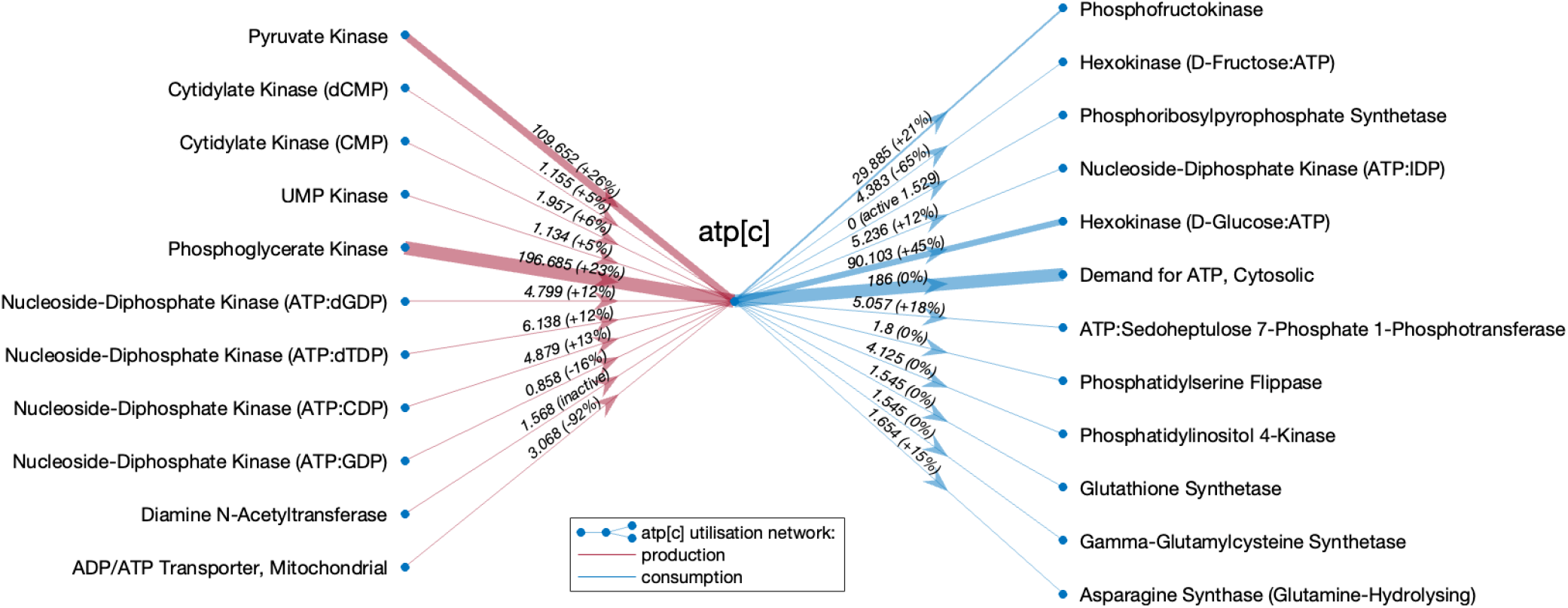
Cytoplasmic ATP utilisation network in the iDopaNeuroC model after Complex V inhibition. Reactions producing (red) and consuming (blue) cytoplasmic ATP, above a threshold in flux magnitude (> 1 *μmol*/*gDW*/*hr*), with magnitude of predicted flux in Complex V inhibition (in proportion to arrow thickness) and percentage change in Complex V inhibition compared to control (% change labels). Fluxes were predicted using maximisation of internal flux entropy and quadratic penalisation of deviation from experimentally measured exchanges as the objective function. The major ATP producting reactions, pyruvate kinase and phosphoglycerate kinase, are predicted to both increase in flux due to Complex V inhibition compared to the control model, while the net mitochondrial ATP production is strongly reduced (ADP/ATP Transporter, Mitochondria) after complex V inhibition. Similarly, the glycolytic ATP-dependent reactions, phosphofructokinase and hexokinase, also are predicted to have an increased flux after complex V inhibition.

**Figure 25:**
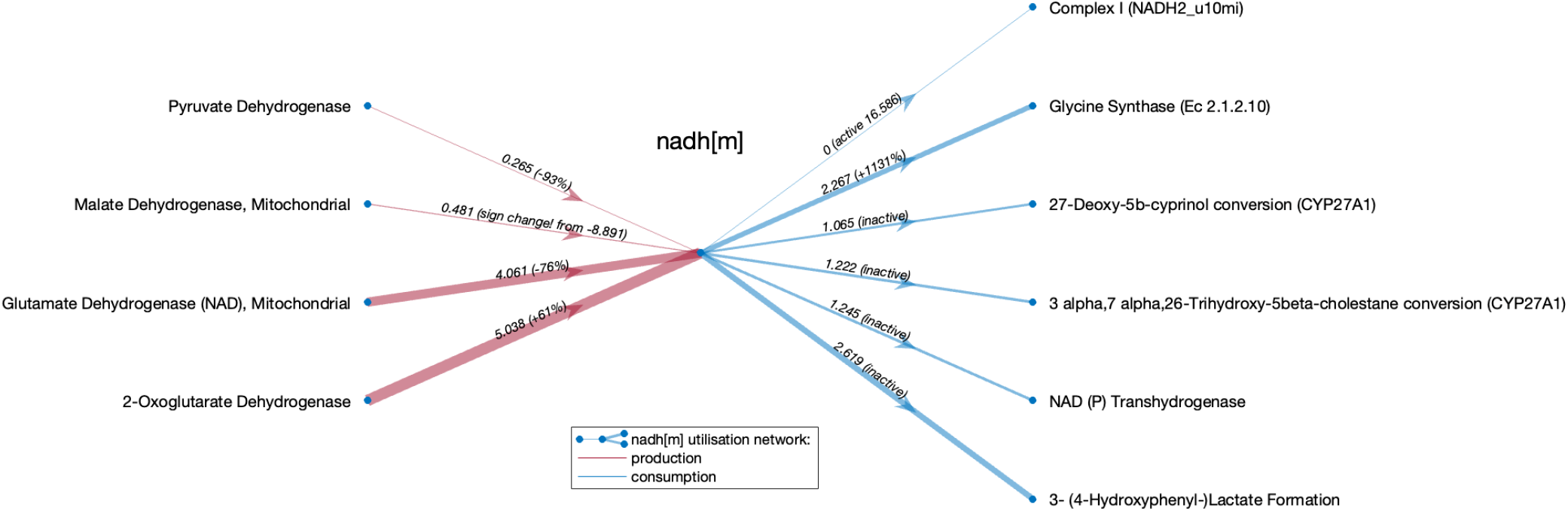
Mitochondrial NADH utilisation network in the iDopaNeuroC model after Complex V inhibition. Reactions producing (red) and consuming (blue) mitochondrial NADH, above a threshold in flux magnitude (> 0.1*μmol*/*gDW*/*hr*), with magnitude of predicted flux in Complex V inhibition (in proportion to arrow thickness) and percentage change in Complex V inhibition compared to control (% change labels). Fluxes were predicted using maximisation of internal flux entropy and quadratic penalisation of deviation from experimentally measured exchanges as the objective function. All reactions significantly contributing to the production of mitochondrial NADH are predicted to be significantly affected by the Complex V inhibition. While pyruvate dehydrogenase and glutamate dehydrogenase (NAD) have both significantly reduced flux, the 2-oxaloglutarate dehydrogenase flux is increased and malate dehydrogenase is predicted to change in reaction directionality (from consumer to producer of NADH) in response to the Complex V inhibition. Furthermore, several reactions that oxidise NADH back to NAD+ have become active, including bile acid-related reactions of CYP27A1, or have an increased flux (glycine synthase), with one exception - the reaction representing Complex I activity has became inactive in response to the Complex V inhibition.

Another interesting prediction is the increase of complex II flux in response to both complex I inhibition and galactose media (Fig. 15). Complex II, also known as a succinate dehydrogenase (SDH) creates a unique link between the TCA cycle and oxidative phosphorylation. Succinate dehydrogenase oxidises succinate to fumarate and transfers electrons via FAD clusters to ubiquinone, reducing it to ubiquinol. Therefore, in healthy cells both complex I and complex II reduce ubiquinone which donates electrons to complex III and leads to an increase in the membrane potential. However, following complex I inhibition succinate dehydrogenase upregulation is predicted to partially restore mitochondrial ATP production through complex V. Previously, upregulation of complex II was reported in PD patients with PINK1 mutations who had an unusually late onset and a mild progression of the disease [106, 107]. Furthermore, all perturbed variants of iDopaNeuroC model were able to secrete dopamine and serotonin, maintaining their dopaminergic phenotype.

In summary, the predicted metabolic changes in response to metabolic perturbations show important flux re-distribution in central energy metabolism. It shows several similarities between complex I inhibition and PD due to PINK1 mutation, and is in line with literature data. In the future, a similar methodology could be used to generate context-specific personalised models of different PD patients to unravel underlying metabolic similarities, and hopefully bring more insights into a PD mechanism.

### Experimental design

Algorithmic experimental design was used to propose experimental designs that optimise the information obtained in future exometabolomic experiments. Algorithmic design of exometabolomic experiments enables optimal selection and development of targeted mass spectrometry platforms for future analyses. This is important as one targeted analytical platform cannot quantify the concentration of all of the metabolites within the iDopaNeuroC model. Our uncertainty reduction pipeline rank orders unmeasured exchanged metabolites by the degree to which their measurement would shrink the feasible set of steady-state flux vectors. In addition to metabolites accessible to mass spectrometry, this analysis also identified protons, bicarbonate and oxygen as targets for additional experimental measurements. For example, the iDopaNeuroC model predicts that future exometabolomic experiments should assess acid-base balance, as well as gas exchange.

## Conclusions

Herein, we present data-driven, context-specific, genome-scale, constraint-based mechanistic models of dopaminergic neuronal metabolism. The combined results of literature curation and omics data generation were integrated together with comprehensive reconstruction of human metabolism, using a novel model generation pipeline to generate an ensemble of models. From this ensemble, we identified a condition-specific model (iDopaNeuroC) and a cell-type specific model (iDopaNeuroCT) as they had the highest predictive accuracy when evaluated against exometabolomic data from control dopaminergic neuronal cultures. Furthermore, independent exometabolomic data from four mass spectrometry platforms established that the condition-specific model predicts the consequences of metabolic perturbations with high qualitative accuracy and moderate quantitative accuracy, representing a breakthrough in predictive fidelity for modelling of human metabolism in non-growing cells. The iDopaNeuro models provide a validated platform for experimental data-driven mechanistic computational modelling, optimal design of experiments and ultimately, provides an objective, quantitative framework for development of drugs targeted toward the aetiopathogeneses of Parkinson*’*s Disease.

## Acknowledgements

The authors would like to thank Sylvain Arreckx, Laurent Heirendt and Thomas Pfau for helping with computational issues, Swagatika Sahoo explanation of the reconstruction process, and Sabine Dier at EURICE for SysMedPD project administration support. ELM, MO, and DE were supported by Aides à la FormationRecherche Training allowances granted to by the Fonds National de la Recherche Luxembourg. ZZ and EG were supported by the Fonds Nationale de la Recherche, Luxembourg, as part of the BMBF-funded e:Med project MitoPD (INTER/BMBF/13/04) and the CORE INTER project MiRisk-PD (FNR11676395). SV and YL were supported in part by NSF awards CCF-2007443 and CCF-2134105. The analysis of RNA sequencing presented in this paper were carried out in part using the HPC facilities of the University of Luxembourg. The provision of NESC from the StemBANCC project is gratefully acknowledged. GP, ELM, AW, CW, JM, FM, DE, MO, CG, HH, AH, JS, TH and RF received funding from the European Union*’*s Horizon 2020 research and innovation programme, for the project SysMedPD, under grant agreement No 668738.

## Author contributions

German Preciat: Methodology, Software, Investigation, Validation, Writing - Original Draft

Agnieszka Wegrzyn: Methodology, Software, Investigation, Validation, Writing - Original Draft

Edinson L. Moreno: Methodology, Investigation, Writing - Original Draft

Cornelius C.W. Willacey: Methodology, Investigation, Writing - Original Draft

Jennifer Modamio: Methodology, Software, Investigation, Writing - Original Draft

Fatima L. Monteiro: Methodology, Formal analysis, Investigation, Writing - Original

Draft Diana El Assal: Methodology, Investigation, Software, Writing – Review & Editing

Alissa Schurink: Methodology, Investigation

Miguel A.P. Oliveira: Methodology, Software, Investigation, Writing – Review & Editing, Funding acquisition

Zhi Zhang: Software, Writing – Review & Editing

Ben Cousins: Software, Formal analysis, Writing – Review & Editing

Hulda S. Haraldsdóttir: Software

Siham Hachi: Investigation

Susanne Zach: Investigation, Resources

German Leparc: Investigation, Resources

Yin Tat Lee: Software

Bastian Hengerer: Supervision, Investigation, Resources

Santosh Vempala: Supervision, Funding acquisition

Michael A. Saunders: Software, Writing - Review and Editing, Formal analysis

Amy Harms: Methodology, Writing - Review and Editing

Enrico Glaab: Investigation, Resources, Funding acquisition

Jens C. Schwamborn: Resources, Supervision, Funding acquisition

Ines Thiele: Conceptualization, Software, Writing - Review & Editing, Supervision, Funding acquisition

Thomas Hankemeier: Conceptualization, Methodology, Resources, Writing - Review & Editing, Supervision, Project administration, Funding acquisition

Ronan M.T. Fleming: Conceptualization, Methodology, Software, Validation, Formal analysis, Writing - Ori- ginal Draft, Writing - Review & Editing, Supervision, Visualization, Project administration, Funding acquisition

## Data Availability

All metabolite and reaction identifiers use the namespace of the Virtual Metabolic Human database [51]. Meta- bolomic data was preprocessed using scripts implemented in R. Constraint-based reconstruction, modelling and analysis was implemented in MATLAB (MathWorks Inc.). Computer code enabling the reproduction of all computational steps used for the generation, validation and prospective use of the iDopaNeuro models is available from https://github.com/opencobra/COBRA.papers/2023_iDopaNeuro. This code depends on the COBRA Toolbox [20] version 3.4+, which contains the XomicsToModel code for model generation and thermoKernel model extraction algorithm and entropicFluxBalanceAnalysis for maximisation of flux entropy, with the option of quadratic penalisation of deviation from measured experimental fluxes. Linear and quadratic optimisation problems were solved using Gurobi 9.1 (Gurobi Inc). Other nonlinear convex optimization problems were solved using the exponential cone solver within Mosek 10.0.30 (Mosek ApS) or the open source solver *Primal-Dual interior method for Convex Objectives* PDCO implemented in MATLAB (MathWorks Inc.). Each of these solvers is interfaced with the COBRA Toolbox.

## Supporting Information

### A Cell number

### B Reconstruction of dopamine metabolism

### C Constraint-based modelling: an introduction

All constraint-based modelling predictions are derived from optimisation problems, typically formulated in the form:

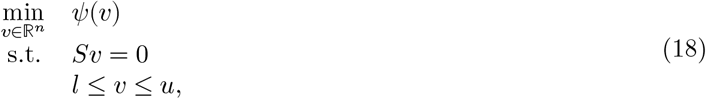

where *S* ∈ ℝ^*m*×*n*^ is a stoichiometric matrix of *m* metabolites and *n* reactions representing a biochemical network, *ν* ∈ ℝ^*n*^ is the vector representing the flux through all of the reactions in a network and *ψ* : ℝ^*n*^ → ℝ is an objective function, which is typically convex. In a constraint-based metabolic model of reaction fluxes, the set of feasible steady-state flux vectors forms a polyhedral convex solution space, defined by the equality and inequality constraints in Equation (18), enabling optimisation of a variety of convex objective functions over this set.

The matrix *S* can be split horizontally into two matrices corresponding to internal, *N* ∈ ℤ^*m*×*k*^, and external, *B* ∈ ℝ^*m*×(*n*−*k*)^, reactions, with corresponding internal and external rate vectors, *z* ∈ ℝ^*k*^ and *w* ∈ ℝ^*n*−*k*^. While all internal reactions are characterised by being mass and charge balanced, external reactions are, on the other hand, imbalanced reactions. External reactions are classified in sink, demand or exchange reactions. A demand reaction allows the accumulation of a compound. A sink reaction allows the production of a metabolite. Finally, an exchange reaction allows the exchange of a metabolite across the extracellular boundary of a system, providing a mechanism to transfer metabolites between the environment and the extra-cellular fluid. Such reactions are distinct from transport reactions, which transfer metabolites between compartments within the model, including the extracellular compartment. Exchange reactions are added to a model to allow certain metabolites to be exchanged across the boundary of the system at variable rates.

The linear equality, *Sν* = 0 in Equation (18), represents mass balance for all the metabolites. This means, for each metabolite the rate of metabolite consumption is equal to the rate of metabolite production. In Equation (18), *Sν* = 0 implies that *Nz* = −*Bw* where internal production plus external input equal internal consumption plus external output. For certain intracellular metabolites, those not exchanged across the boundary of the system, we assume they are at a steady-state, so we have *N*_*i*_*z* = 0, where *N*_*i*_ denotes the *i*^*th*^ row of the internal stoichiometric matrix. Additional linear inequalities keep reaction rates between lower and upper bounds, *l* and *u*, respectively.

#### Bounds on reaction rates

In each metabolic reaction, *ν*_*i*_, is constrained between a lower and an upper bound, *lb* ≤ *ν*_*i*_ ≤ *ub*. The default reaction lower and upper bounds are commonly set based on model characteristics and constraints value. Lower and upper bounds were set to include fluxes from metabolite concentration in the media, e.g., glucose flux rate based on media composition (−5,430.74 *μ*mol/gDW/hr). Reactions can be reversible or irreversible. A reaction is said to be reversible in the case where it has a negative *lb* and a positive *ub*. When the *lb* is set to zero and the *ub* is a positive number the reaction proceeds in the forward direction. Similarly, when the *ub* is zero and the *lb* is a negative number the reaction occurs in the backward direction. In a metabolic model, exchange of metabolites with its environment is represented by constraints on the corresponding exchange reactions, which define the boundary conditions of the model. If a metabolite is taken up, the corresponding exchange reaction has a negative number as *lb* and zero as *ub*, whereas if it is secreted, the *lb* is set to zero and the *ub* is a positive number.

### D Evaluation of different objectives

### E Perturbation-induced metabolic changes with iDopaNeuroC model

### F GBA1 knock out prediction

Figure 26 illustrates the predicted consequence of knocking out the GBA1 gene, corresponding to the Glucosylceramidase reaction (GBA) which is

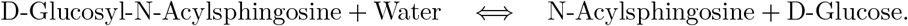

### G iDopaNeuro cell-type- and condition-specific results

#### G.1 Analytical chemistry

The iDopaNeuroCT model includes exchange reactions for 238 metabolites, the concentrations of 62 of which were known either from the cell media data or from exometabolomic measurements, therefore there still remains 152 metabolites to target with additional exometabolomic platforms to completely quantitatively constrain all exchanges (Fig. 27).

**Figure 26:**
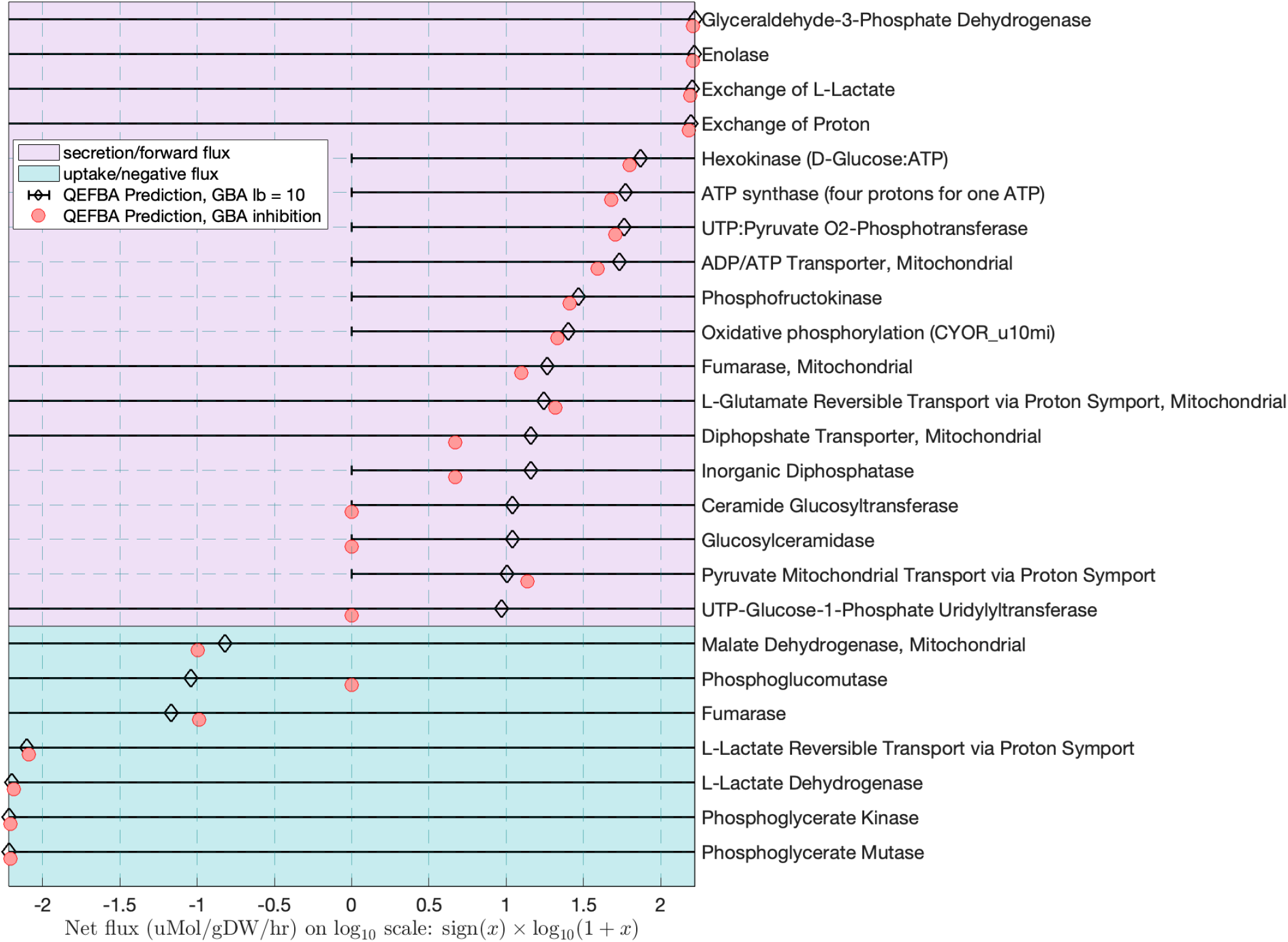
Predicted consequence of Glucosylceramidase inhibition. Comparison of predicted dopaminergic neuronal metabolic flux in the Glucosylceramidase reaction (GBA) in a normal dopaminergic neuron (diamonds), and a GBA1 knock-out (red dots), assuming 10 and 0 *μ*mol/gDW/hr flux, respectively. In both cases, fluxes were predicted using quadratic penalisation of exchange flux deviation from measured exchanges in glucose maintenance media and maximisation of internal reaction flux entropy as the objective (QEFBA). Reactions are only displayed where there is a difference of >3 *μ*mol/gDW/hr difference between control and GBA1 knock out. The bounds on reaction flux in the control model are illustrated by the bars.

**Figure 27:**
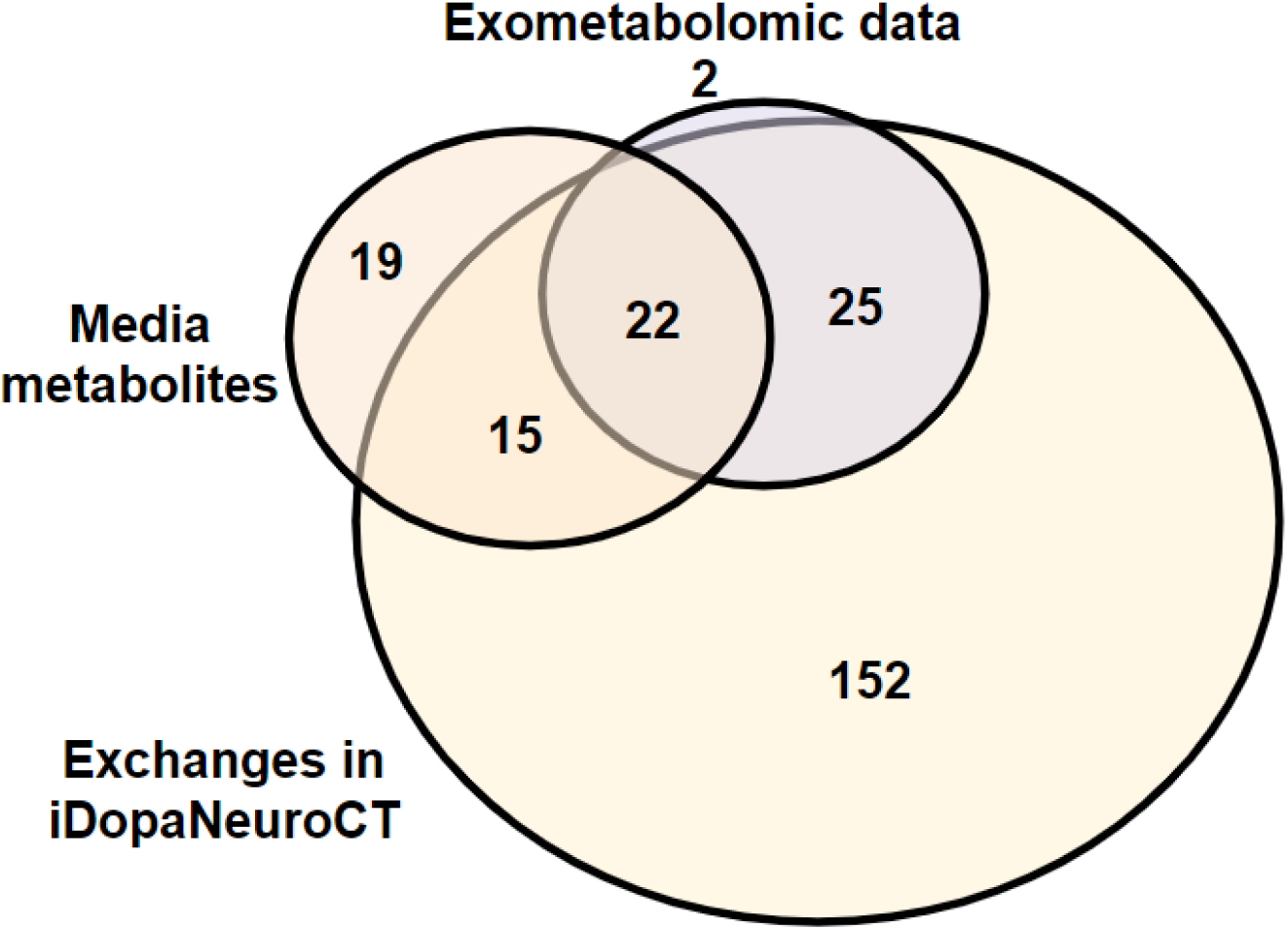
Venndiagram summarising metabolomic measurements. A total of 49 unique metabolites were targeted by the selected metabolomic platforms (purple), of which 47 could be used to constrain the model as 2 metabolites (glutarate, methylmalonate) were not present in the stoichiometrically and thermodynamically flux consistent subset of Recon3D. Of these 47 metabolites, 22 were present in the fresh medium (orange) and 25 were synthesised by the cells and secreted into the spent medium. The fresh media contained 56 metabolites, of which 37 corresponded to exchanges in the cell-type specific dopaminergic neuronal model, while 19 metabolites (mainly ions, e.g., Na2+) were either not in Recon3D or not present in the stoichiometrically and thermodynamically flux consistent subset of Recon3D.

#### G.2 Bibliomic data

The bibliomic context-specific data from dopaminegic neuronal metabolism integrated to the iDopaNeuroCT model is represented in the Figure 28

**Figure 28:**
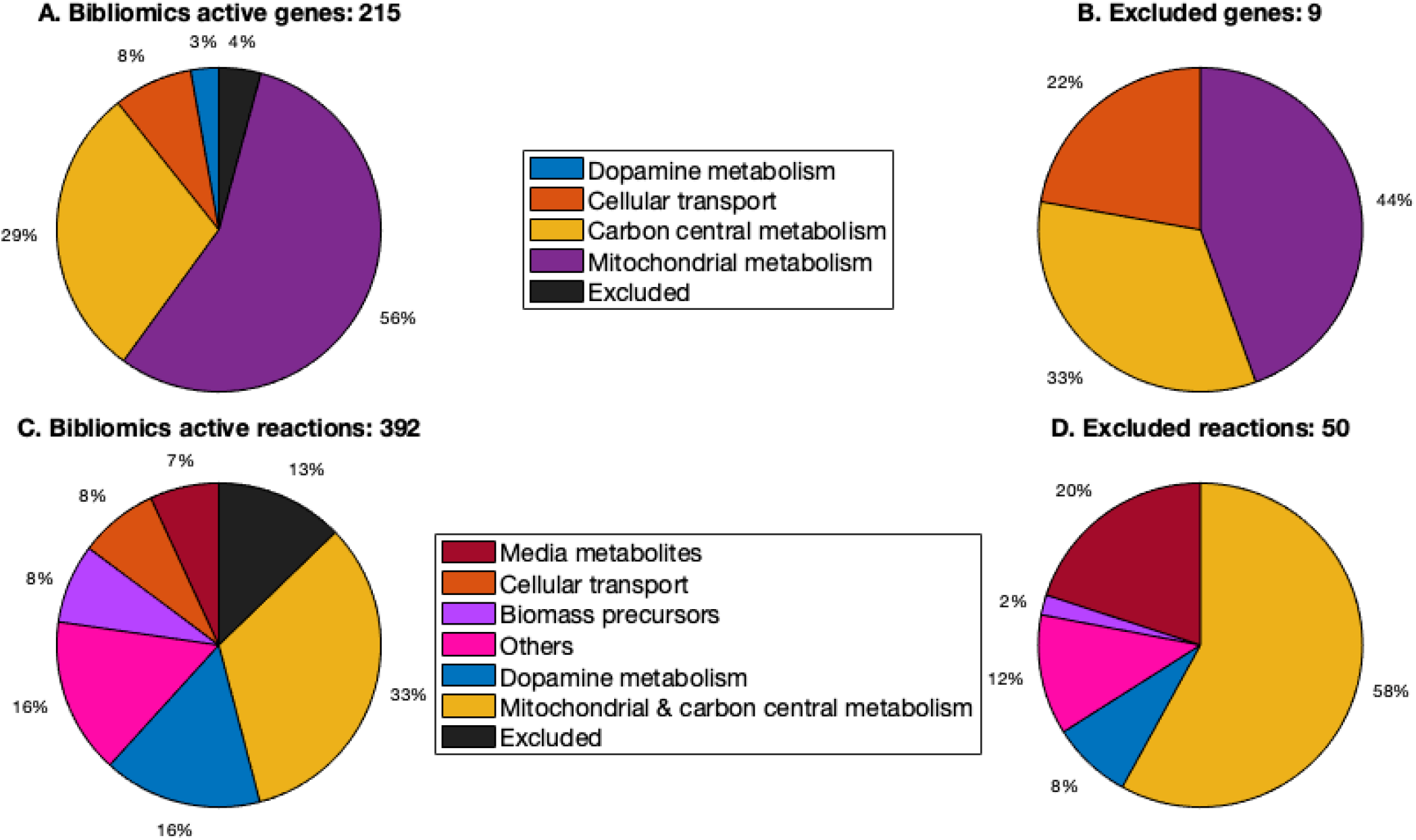
Classification of active reactions and genes by manual literature curation. The result is partly a reflection of the availability of biochemical information on certain pathways, e.g., central metabolism, and partly a reflection of the pathways that were targeted for curation, e.g., dopamine metabolism.

**Figure 29:**
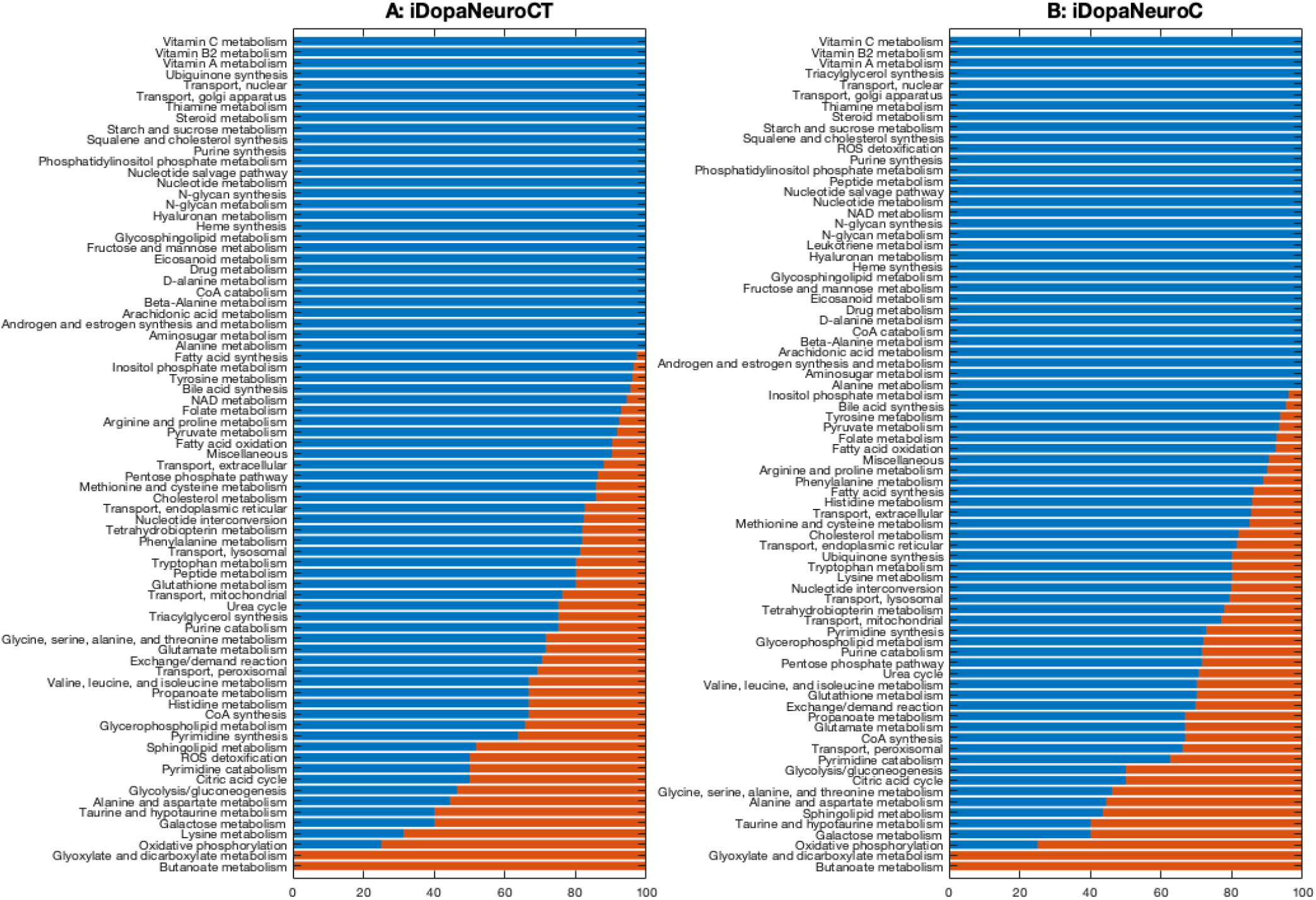
Minimal flux metabolic subsystems. Comparison of the fraction of active reactions (red) in the most sparse flux vector in each metabolic subsystem of (**A**) the iDopaNeuroCT model and (**B**) the iDopaNeuroC, obtained by minimising the function *ψ*(*ν*) := ‖*ν*‖_0_, to predict the minimum number of reactions that are required to be active to satisfy dopaminergic neuron specific constraints on the steady-state flux space Ω in glucose maintenance medium.

**Figure 30:**
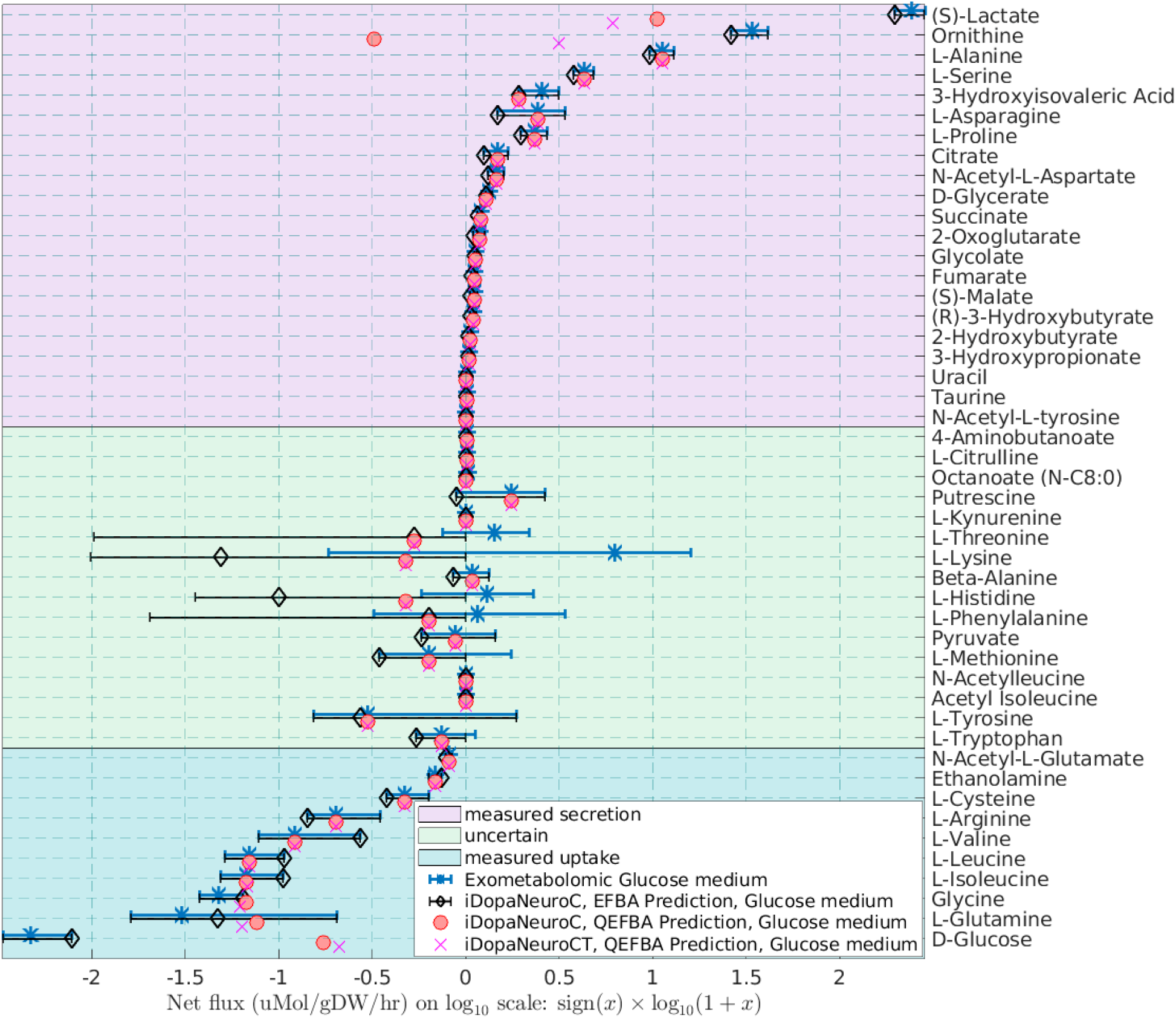
Fitting of measured metabolite exchanges. When fresh versus spent media concentrations were compared, metabolites were grouped into those that were measured to be secreted (positive average minus one standard deviation, pink background), measured to be taken up (negative average plus one standard deviation, blue background) and measured with uncertain exchange (error bar includes zero, green background). During the dopaminergic neuronal model generation process, exchange fluxes were adjusted from the generic bounds in Recon3 to bounds to intervals derived from measured exchange fluxes (mean +/-standard deviation), except in the case that it was infeasible to obtain a steady state flux and not secreting any essential metabolite, whereupon quadratic minimisation was used to minimise the deviation from the experimentally measured interval (blue). Thus, certain dopaminergic neuronal exchange flux bounds (black) were allowed to be relaxed, especially in the case that measurement interval spanned zero. Maximisation internal flux entropy alone (EFBA) results in predictions within these bounds. Maximisation internal flux entropy and quadratic penalisation from experimentally measured fluxes (QEFBA) generally results in predictions within the aforementioned bounds but may deviate outside (e.g. D-glucose). The results of fitting are very similar for the condition-specific (iDopaNeuroC) and cell-type specific (iDopaNeuroCT) models, using maximisation internal flux entropy and quadratic fitting to measured exchange fluxes in glucose medium. Note that measured exchange fluxes extend from −213 ± 86 *μmol*/*gDW*/*hr* for glucose to 241 ± 44 *μmol*/*gDW*/*hr* for lactate and are significant over ∼5 orders of magnitude, e.g., Uracil is secreted at a rate of 0.0118 ± 0.0048 *μmol*/*gDW*/*hr*.

**Figure 31:**
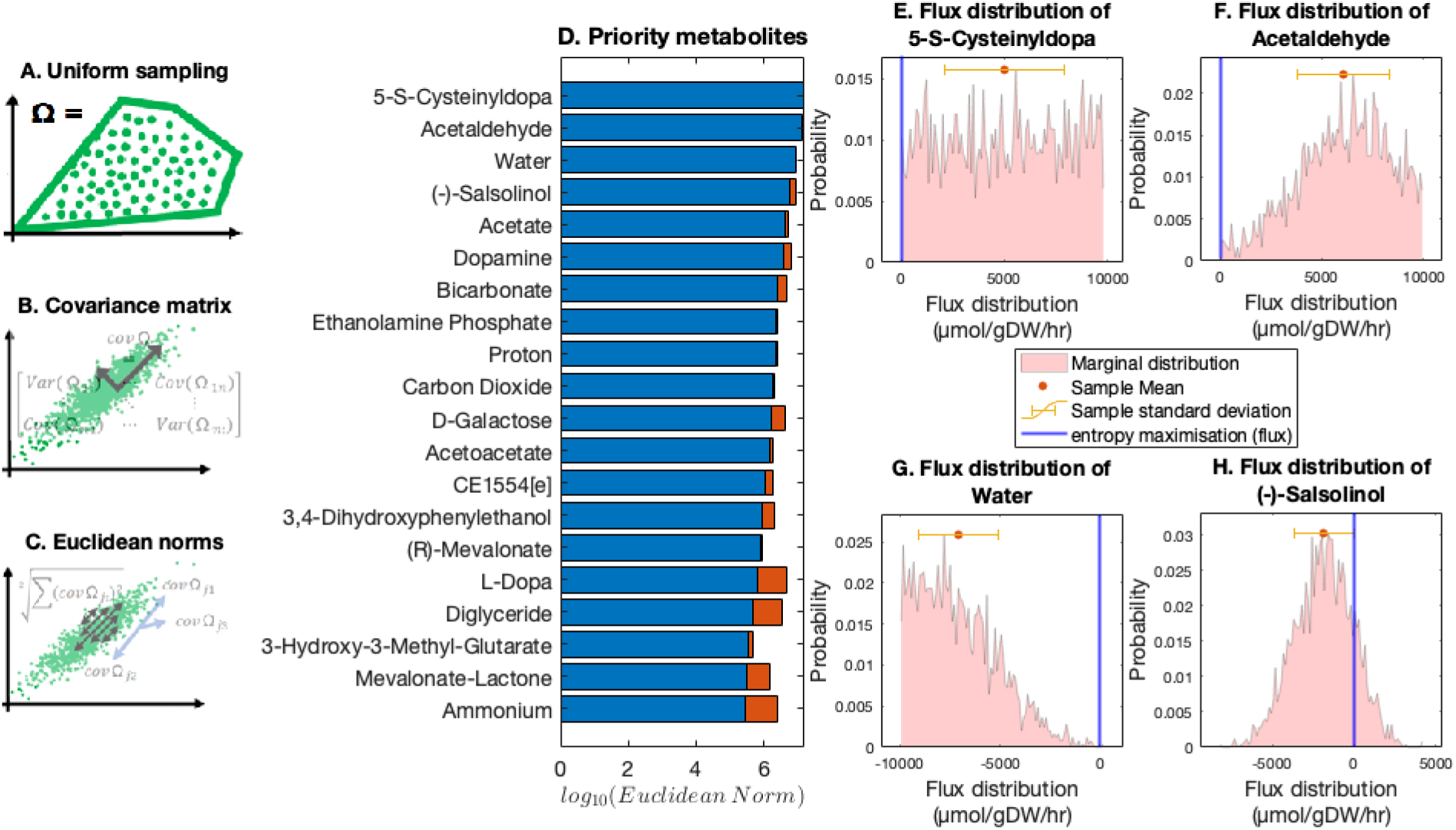
Prospective prioritisation of model variables to constrain for the iDopaNeuroCT model. **A**. Uniform sampling of the steady-state flux space (Ω ≔ {*ν* | *Sν* = 0; *l* ≤ *ν* ≤ *u*} in 3), of the iDopaNeuroCT model. **B**. Computation of the covariance matrix of sampled external fluxes. **C**. The Euclidean norm of each row of the covariance matrix identifies the exchange reaction with the highest degree of freedom. **D**. The predicted most informative metabolites to measure, each corresponding to one external reaction flux, the iDopaNeuroCT model. The variance reduction (to blue) due to cumulative constraints on higher-ranked metabolites (red) is taken into account in the ranking. **E-H**. The flux distribution of the four most important metabolites to constrain in order to optimally shrink the steady-state flux space Ω. Additionally, the fluxes of the metabolites were estimated using the maximisation of the entropy of all forward and reverse net fluxes (*ψ*_*ν*_ in 1; blue).

#### G.3 Model characteristics

**Minimal reaction set**. Sparse Flux Balance Analysis [108] was used to predict the number of active reactions in the iDopaNeuro models that are consistent with the constraints on the feasible set of fluxes, that is, a solution to the following problem

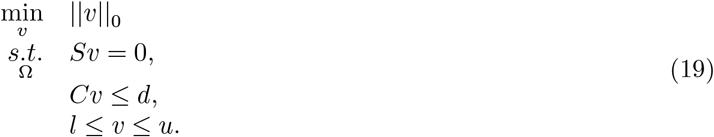

Each active reaction in the optimal solution is then tested to see if it is essential to satisfy the constraints.

The iDopaNeuroCT model represents the activity of 1,229 metabolic genes from 77 biological pathways, comprised of 2,065 biochemical reactions that interconvert 1,313 metabolites. The sparseFBA solution contains 375 active reactions in 50 subsystems, each reaction of which is essential to be active to satisfy the constraints. The iDopaNeuroC model represents the activity of 1,212 metabolic genes from 78 biological pathways, comprised of 1,915 biochemical reactions that interconvert 1,244 metabolites. The sparseFBA solution contains 360 active reactions in 45 subsystems, all but 3 reactions of which are essential to be active satisfy the constraints.

#### G.4 Fitting to measured metabolite exchanges

#### G.5 Uncertainty reduction

Metabolites that would most shrink the flux space of the iDopaNeuroCT model identified using uniform sampling [66] are shown in Figure 31.

